# Fully integrative species delimitation with machine learning and diverse data types in *delimSOM* 2.0

**DOI:** 10.64898/2026.10.04.756574

**Authors:** Daniel Schönberger, R. Alexander Pyron, Julian R. Dupuis

## Abstract

Species delimitation increasingly relies on multiple sources of evidence, but most frameworks are still limited to genetics and morphology or analyze data types separately and compare results qualitatively. We present *delimSOM* 2.0, an R package that leverages unsupervised machine learning through multilayer self-organizing maps. Any kind of input data – including genomic, phenotypic, ecological, and spatial – is analyzed jointly, retaining each data type as a separate layer and balancing their contributions through layer-specific weighting. Our package provides functions for preprocessing, replicate training, multivariate clustering, hyperparameter optimization, model-quality assessment, and variable- and layer-importance analyses. We evaluated our approach using two simulation sets and eight empirical studies spanning diverse taxa and data types. Overall, performance was strongest with larger sample sizes (>10 individuals per lineage) and modest numbers of lineages (K < 5), but unlike most species delimitation methods, it can reliably infer K = 1. Across two-population demographic simulations of genetic data, our method performed similarly to the well-established *DAPC* across gene-flow and sampling regimes, was more conservative than *STRUCTURE* and *sNMF* in inferring K = 2 and showed mixed support for K = 1 and K = 2 at intermediate differentiation levels and with gene flow. Crucially, our method is robust to missing data and variation in hyperparameters. Empirical examples showed that genetic data were typically most influential, but phenotypic, ecological, and spatial layers often contributed substantial complementary information. By combining diverse data types into a single unsupervised analysis, *delimSOM* 2.0 enables fully integrative species delimitation, reduces over-splitting, and advances efforts toward more objective and standardized taxonomy and speciation biology.

## INTRODUCTION

Integrative species delimitation is the process of evaluating candidate species and estimating their boundaries using multiple complementary sources of evidence (Padial et al., 2010; Schlick-Steiner et al., 2010). It has become a central objective in systematics because speciation is a multidimensional evolutionary process in which different attributes of lineage separation—such as genetic differentiation, phenotypic divergence, ecological specialization, and geographic and reproductive isolation—can arise through different mechanisms at different rates (de Queiroz, 2007). Consequently, species boundaries often emerge gradually along a continuum of divergence rather than at discrete points (Stankowski & Ravinet, 2021), making species delimitation inherently challenging. This view is consistent with the unified species concept, under which species are separately evolving metapopulation lineages diagnosed using various biological properties (de Queiroz, 2007). As a result, robust species hypotheses are more likely to emerge when complementary sources of evidence are analyzed together, limiting biases from any single data type (Dayrat, 2005; Padial et al., 2010; Schlick-Steiner et al., 2010). This also facilitates the identification of lineages with subtle or uneven differentiation across multiple dimensions, while reducing the risk of oversplitting where intraspecific variation and population-level structure are mistaken for species-level divergence (Chambers et al., 2025; Chan et al., 2017, 2020; Dufresnes et al., 2023; Sukumaran & Knowles, 2017).

Although integrative species delimitation has been advocated for over two decades (Dayrat, 2005), most approaches described as “integrative” have combined evidence comparatively or iteratively rather than analyzing multiple data types jointly. In a comparative workflow, different lines of evidence are analyzed separately, and the resulting partitions are compared *a posteriori* for congruence (Carstens et al., 2013; Pyron et al., 2023; Raxworthy et al., 2007). For example, genetic lineages may be inferred from genomic data, morphometric clusters from morphology, and niche differences from climatic data. In an iterative workflow, candidate lineages are first identified from one source of information and then successively evaluated, refined, or rejected, usually first with genetic data and then with morphology, ecology, or behavior (Chan et al., 2020; van Elst et al., 2024; Yeates et al., 2011). Although these approaches are not mutually exclusive (Schlick-Steiner et al., 2010), both integrate conclusions across separate analyses rather than estimating species boundaries jointly.

Few species delimitation approaches have analyzed multiple data types in ways that might be described as fully integrative. Guillot et al. (2012) developed a unified Bayesian model for phenotypic and genetic data that incorporates spatial information, but its computational demands limit scalability to large datasets and its model is restricted to these three data types. Edwards & Knowles (2014) presented a statistically integrative method for species detection and individual assignment using genetic, morphological, and ecological data, but it was designed to generate putative species assignments for subsequent evaluation rather than a complete species delimitation workflow. Solís-Lemus et al. (2015) introduced a Bayesian approach that integrates gene sequence data and quantitative traits for species delimitation, but it requires predefined candidate species and a guide tree. Consequently, there remains a need for approaches that can jointly analyze broad combinations of data within a single workflow while retaining robustness, ease, and computational efficiency.

Machine-learning (ML) approaches offer a promising path toward species delimitation because they can accommodate complex heterogeneous datasets that differ in scale, variance, and completeness, without requiring explicit parametric models and making fewer assumptions about distributions and statistical properties (Derkarabetian et al., 2019; Karbstein et al., 2024; Pyron, 2023). Recent work has shown that ML algorithms can be effective species delimitation tools in both supervised settings, where classifiers are trained on simulated data to distinguish alternative species or demographic scenarios (Pei et al., 2018; Smith & Carstens, 2020), and in unsupervised settings, where lineage structure is inferred from observed data without *a priori* labels (Derkarabetian et al., 2019; Pyron, 2023). Furthermore, some recently developed multimodal deep-learning methods have combined molecular data and morphological images (Kösters et al., 2025; Yang et al., 2022), but these are supervised species-identification rather than *de novo* species-delimitation tools. Therefore, most ML applications in species delimitation remain focused on genetic or morphological data, while computationally tractable methods that jointly integrate multiple heterogeneous data types are still in their infancy (Karbstein et al., 2024).

Recently, Pyron et al. (2023) introduced an individual-based, unsupervised ML approach using self-organizing maps (SOMs) to delimit candidate species from genetic data. SOMs project complex multivariate relationships onto a two-dimensional map grid while preserving local topological structure (Kohonen, 1998, 2014). Candidate lineages are inferred by clustering the trained map units without requiring predefined species assignments. Pyron (2023) subsequently introduced *delimSOM*, which extended this approach to multilayer SOMs capable of integrating multiple data types. This linked unsupervised clustering with integrative species delimitation through a shared competitive-learning framework. While the original *delimSOM* implementation consisted of analysis scripts for basic SOM-based analyses, a comprehensive workflow and flexible *R* package remained lacking.

Here, we present *delimSOM* 2.0, an *R* package with user-friendly functions for fully integrative species delimitation. Any data type that can be represented quantitatively for a common set of individuals can be incorporated, including count, binary, continuous, and categorical variables from genomic, morphological, behavioral, host-association, ecological, and spatial data. Different lines of evidence are retained as separate layers but jointly inform a shared map within a single analytical framework through layer-specific weighting that balances their contributions. We provide a workflow for data preprocessing, SOM training, candidate lineage assignment, hyperparameter optimization, output extraction, model-quality assessment, visualization, and comparison of alternative clustering strategies (Supplementary Table S1). Relative to the previous implementation, *delimSOM* 2.0 advances the framework through updated training and clustering functions, alternative clustering and K-selection strategies, new data-processing functions, revised metrics of variable and layer importance, and improved computational scalability for large datasets.

We comprehensively test *delimSOM* 2.0 using eight empirical studies and two simulation sets to assess its performance across diverse species-delimitation scenarios. We ask whether SOMs recover species-level divergence rather than intraspecific population structure, which SOM parameters most reliably recover known structure, how accuracy is affected by missing data, sample size, and uneven sampling, and how *delimSOM* 2.0 performs across divergence-time and gene-flow scenarios relative to other commonly used population-genetic methods. We provide a step-by-step tutorial (https://github.com/rpyron/delim-SOM).

## METHODS

### SOM framework

A Self-Organizing Map (SOM) organizes many different data types from specimens into a two-dimensional grid map so that specimens with similar overall patterns tend to be represented near one another (Kohonen, 1998, 2014). By summarizing many individual observations into fewer representative local patterns, SOMs can reduce sensitivity to noise and reveal broad structure in high-dimensional biological datasets where relationships are often non-linear, partly conflicting, or differently distributed across variables (Kohonen, 1998). Different types of data, such as genetics and climate, are kept as separate “layers” but contribute jointly to the map. The map consists of a grid of cells, each storing a representative local “average” pattern (called a codebook vector) for the variables in each data layer. During training, each specimen is compared with all grid cells and assigned to the cell whose pattern is most similar to its overall combination of measurements – the best-matching unit. The best-matching unit and nearby cells are then adjusted to become slightly more similar to that specimen. After repeated updates (“steps”), the grid becomes organized so that neighboring cells represent specimens with similar overall measurements, whereas dissimilar specimens occur farther apart on the map. After the map has been trained, the representative map cells (i.e., the codebook vectors) are grouped into clusters, and the best-supported number of clusters (K) is evaluated using various methods (Supplementary Methods M1). These clusters are then interpreted as candidate lineages.

### Data processing

We provide preprocessing functions to prepare different input data for SOM training and clustering (Supplementary Methods M2). Genetic inputs are converted to biallelic dosage matrices and categorical variables to binary indicators. Highly correlated or low-variation variables are removed. Only individuals shared across all layers are retained. Within each layer, variables with excessive missingness or no variance are removed. Retained variables are min-max scaled to a range from zero to one so that different scales contribute comparably to training. Missing values are accommodated by calculating sample-to-unit distances and codebook-vector updates only across observed dimensions, allowing partially incomplete samples to contribute (Samad & Harp, 1992).

### SOM training

Training (Supplementary Methods M3) is performed using hexagonal grids, which support neighborhood relationships and facilitate visualization (Kohonen, 1998). Grid dimensions are determined automatically based on sample size and the primary axes of variation in the data, so that map geometry reflects the data structure while controlling computational cost and topological distortion (Kohonen, 2014). This is important because grid dimensions influence performance and resolution: large grids can increase computation and produce sparsely occupied units, whereas small grids can merge distinct multivariate structures (Kohonen, 1998).

Codebook vectors are initialized by randomly sampling observations from the input data (Wehrens & Kruisselbrink, 2018), so that initial prototypes reflect observed data combinations. At each training step, the codebook vectors of the best-matching and surrounding units are moved toward an individual’s values, with the magnitude of the update determined by the learning rate and neighborhood function. Two neighborhood functions are implemented, which control how the codebook vectors of the best-matching and neighboring units are updated (Kohonen, 1998): the bubble function applies equal influence on all units within the neighborhood radius, whereas the Gaussian function applies influence that declines smoothly with grid distance from the best-matching unit. For both, the neighborhood radius decreases during training, shifting learning from broad map organization to finer local refinement (Wehrens & Kruisselbrink, 2018).

Because training is sensitive to random initialization and input order, multiple replicate maps are trained to assess stochastic variation and summarize recurrent structure across runs. Replicates can be executed in parallel to reduce runtime. Learning proceeds with a linearly decreasing learning rate, with stronger adjustments of the codebook vectors during early iterations supporting broad map organization and progressively weaker adjustments enabling finer local refinement (Kohonen, 1998). Learning-rate tuning can be used to evaluate candidate initial and final rates and select the combination producing the best fit (lowest mean quantization error) across replicates.

### Multi-layer integration

Different datasets are retained as separate layers with layer-specific distance functions while contributing jointly to training. Distances between input samples and codebook vectors are calculated separately for each layer and combined into a weighted overall distance. Normalization of these distances prevents layers with larger numerical scales from dominating the analysis. In addition, user-defined layer weights can be supplied, which are combined multiplicatively with the internal normalization weights to determine each layer’s contribution to the final aggregated distance. These user-defined weights can be used to emphasize or down-weight data types according to analytical goals, prior hypotheses, or measurement uncertainty. For example, layers with uncertain measurements or lower information may be down-weighted, whereas layers expected to contain stronger evolutionary signal may be emphasized.

### Clustering and inference of candidate lineages

After training, the codebook vectors are clustered, the best-supported number of clusters is determined, and each sample is assigned to the cluster containing its best-matching unit (Supplementary Methods M4). Since the performance of clustering and K-selection methods depends on data structure and cluster geometry (Omran et al., 2007; Rodriguez et al., 2019), we implemented and tested six approaches: centroid-based k-means with BIC-elbow selection, centroid-based k-means with BIC-threshold selection, probabilistic Gaussian mixture modelling with BIC-threshold selection, hierarchical clustering with the Davies–Bouldin index, density-based HDBSCAN, and density-based OPTICS with silhouette-score selection. Importantly, all approaches allow K = 1 when subdivision is insufficiently supported. Poorly fitting replicate maps are removed based on quantization error (measuring sample-to-best-matching-unit fit) and topographic error (measuring neighborhood preservation). Clustering is repeated across retained replicates to evaluate support for alternative values of K and calculate replicate-consensus “species coefficients” (Pyron, 2023) which summarize assignment stability across replicates.

### Evaluation and visualization of results

*delimSOM* 2.0 provides functions to evaluate and visualize results (Supplementary Methods M4–M5). Neighbor-distance plots highlight local discontinuities and potential boundaries in map space, while additional plots summarize sample mappings and cluster boundaries on the SOM, replicate-consensus species coefficients, geographic assignment maps, learning trajectories, support for alternative values of K, and relative layer-distance scales.

We developed complementary variable and layer importance metrics to evaluate how individual variables and data layers contribute to lineage delimitation. These replace the importance metrics used in the original implementation (Pyron, 2023), which relied on median codebook values and layer-distance weights rather than directly quantifying contributions to inferred lineage structure. Two variable-importance metrics are calculated within each layer and summarized across replicates: weighted map variance measures variation across the trained map, whereas weighted η^2^ measures how strongly each variable separates the inferred clusters. Layer importance is assessed using within-layer variable importance and replicate-matched leave-one-layer-out analyses. The latter reruns SOM training and clustering while omitting one layer at a time and evaluates how its omission affects the number of identified clusters, which samples are grouped together, and how clearly samples are assigned to those clusters.

### Simulation tests

We tested the performance of *delimSOM* 2.0 using two simulation sets (Supplementary Methods M6– M7). Simulation set 1 used synthetic multilayer datasets with either a low (K = 3) or high (K = 7) number of simulated groups. This simulation set was used to test which parameter settings (clustering and K-selection method, number of training iterations, neighborhood function, missing-data proportion, and learning-rate tuning) most reliably recovered the true clusters. These results informed our implemented default settings in *delimSOM* 2.0. Each dataset contained four layers representing SNPs, morphology, climate, and host-association, designed to mimic common data types found in empirical datasets. Multivariate differentiation was introduced between groups within each layer.

Simulation set 2 evaluated *delimSOM* 2.0 using genomic data simulated under two-population demographic scenarios spanning a range of divergence times and migration rates. This simulation set tested whether the method can infer K = 1 when differentiation is absent, at what point support shifts toward K = 2 as divergence increases, and how gene flow affects performance. We evaluated complete isolation (no gene flow) and three symmetric secondary-contact migration treatments. Datasets comprising 1,000 unlinked SNPs were simulated in *fastsimcoal2* (Excoffier et al., 2021). Divergence time varied gradually from no to deep divergence (200,000 generations). We analyzed total sample sizes of 48, 32, or 16 diploid individuals, allocated either evenly between populations or with a 3:1 sampling imbalance. Effective population size, mutation rate, and growth rate were held constant. Performance was compared with three commonly used population-genetic methods: *DAPC* (Jombart et al., 2010), *sNMF* (Frichot et al., 2014), and *STRUCTURE* (Pritchard et al., 2000).

### Empirical examples

We evaluated *delimSOM* 2.0 across eight empirical multi-layer case studies spanning amphibians, corals, insects, shrubs, mammals, and sea slugs. Datasets comprised 2–8 complementary data layers, 1–27 candidate lineages, and 22–367 individuals (Supplementary Methods M8 and Tables S2–S3). Layers included genomic SNPs, microsatellite markers, mitochondrial and nuclear marker haplotypes, mitochondrial sequences, continuous and categorical morphology, spatial and environmental variables, symbiont community composition, host association, and developmental mode. The case studies included three eastern North American *Desmognathus* salamander systems (Pyron et al., 2022, 2023, 2024), Indo-Pacific *Pocillopora* corals (Oury et al., 2023), western Canadian *Polygonia* anglewing butterflies (Dupuis et al., 2018), the eastern North American *Viburnum nudum* species complex (Spriggs et al., 2019), Malagasy *Microcebus* mouse lemurs (van Elst et al., 2024), and western Atlantic *Elysia* sea slugs (Krug et al., 2026). When spatial data were available, we used *NicheDiv* (Schönberger et al., 2026) to extract a comprehensive set of environmental variables for inclusion as an ecological layer to evaluate multivariate environmental differentiation.

We highlight two case studies. The first examined four broadly sympatric and morphologically similar *Polygonia* anglewing butterfly species. Dupuis et al. (2018) used mitochondrial DNA, genome-wide SNPs, and visually and digitally scored wing characters to evaluate species boundaries, supporting four species despite overlapping morphology and cytonuclear discordance in relationships. This system provided a strong test of *delimSOM* 2.0’s ability to integrate six diverse layers: genomic, mitochondrial, continuous morphological, categorical morphological, environmental, and spatial. The second investigated the seepage salamander *Desmognathus aeneus*, a small, low-dispersal habitat specialist with deeply structured geographic populations and extensive admixture. Pyron et al. (2024) used genomic, demographic, environmental, geographic, and morphometric evidence, recovering deeply divergent genetic lineages but concluding that recurrent migration, admixture, and limited phenotypic differentiation were consistent with a single species. We included this system to test whether *delimSOM*

2.0 could distinguish strong population structure from species-level divergence (Chambers et al., 2025; Chan et al., 2020; Dufresnes et al., 2023) using genomic, spatial, environmental, watershed, and morphological layers.

## RESULTS

### Simulation set 1: Effects of parameters on performance

Simulation set 1 used synthetic four-layer datasets with known group membership representing low (K = 3) and high (K = 7) lineage numbers to test how clustering method, training iterations, neighborhood function, missing data, and learning-rate tuning affected recovery of the true K, assignment accuracy, SOM map quality, and runtime. *delimSOM* 2.0 performed strongly in recovering the simulated number of lineages and accurately assigning individuals under the K = 3 scenario (Figure 1; Supplementary Figure S1), whereas it resolved the true group structure less reliably under K = 7 (Supplementary Figures S2–S3).

**Figure 1:**
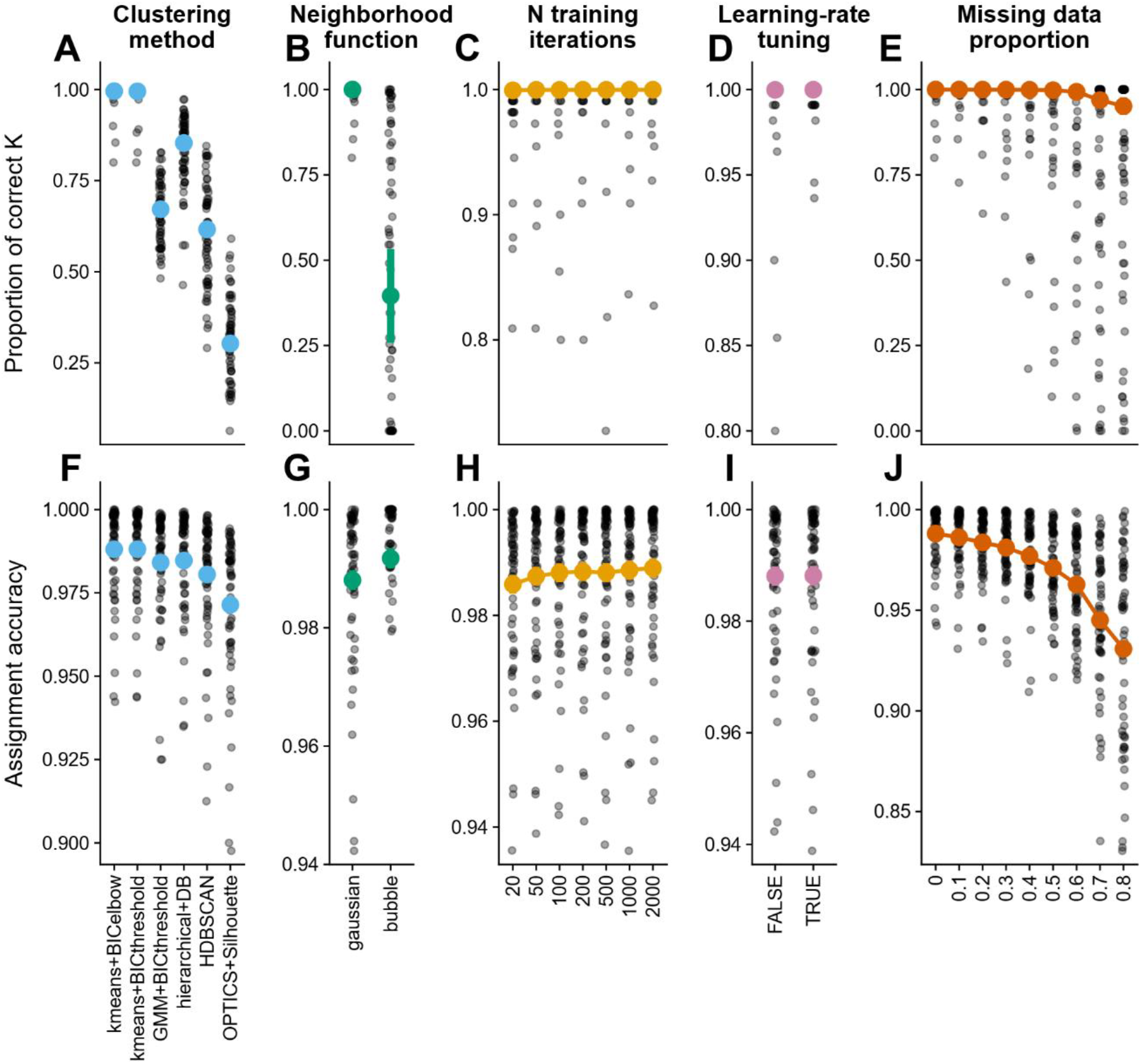
Effects of SOM parameter settings on *delimSOM* 2.0 performance in simulation set 1 with simulated K = 3. Each dataset comprised 20 individuals per group and four layers representing SNPs, morphology, climate, and host association. Upper panels show the proportion of simulations recovering the correct K, and lower panels show assignment accuracy (proportion of correctly assigned individuals among simulations that recovered K = 3). Panels show clustering and K-selection method **(A, F)**, neighborhood function **(B, G)**, training steps **(C, H)**, learning-rate tuning **(D, I)**, and missingness **(E, J)**. Small gray points represent simulation-level results, whereas larger colored points and vertical bars represent estimated marginal means ± 1 standard error from binomial generalized (upper panels) and Gaussian (lower panels) linear mixed models. Abbreviations: BIC = Bayesian information criterion, DB = Davies–Bouldin index, GMM = Gaussian mixture model, HDBSCAN = hierarchical density-based spatial clustering of applications with noise, OPTICS = ordering points to identify the clustering structure.

Performance also depended strongly on the clustering method. Both k-means approaches with BIC-based K-selection consistently recovered K = 3, achieved high assignment accuracy, showed the greatest agreement with the true assignments, and were the computationally fastest overall (Figure 1; Supplementary Figures S1–S3). Although both approaches selected K = 1 rather than K = 7 in the K = 7 scenario, inspection of the BIC curves revealed a distinct elbow at K = 7 that was not automatically selected because of the small initial improvement in BIC between K = 1 and K = 2 (Supplementary Figure S4). GMM and hierarchical clustering also showed high recovery and assignment accuracy but were slower, whereas HDBSCAN and OPTICS were faster but less reliable, particularly under K = 7 (Figure 1; Supplementary Figures S1–S3).

The Gaussian outperformed the bubble neighborhood function in recovering the correct K and produced lower topographic error, whereas the bubble function yielded lower quantization error and shorter runtimes; results for assignment-based metrics were mixed (Figure 1B & G; Supplementary Figures S1– S3). Despite this trade-off, we recommend the Gaussian function because it more reliably recovered the correct K and better-preserved map topology. Increasing the number of training iterations beyond 100 primarily increased computational time, with little effect on lineage recovery, assignment accuracy, or quantization error (Figure 1C & H; Supplementary Figures S1–S3). Likewise, learning-rate tuning improved performance only marginally while substantially increasing runtime (Figure 1D & I; Supplementary Figures S1–S3). *delimSOM* 2.0 was also relatively robust to missing data, as performance declined only marginally as the proportion of missing data increased to approximately 0.5 but decreased more sharply beyond this threshold (Figure 1E & J; Supplementary Figures S1–S3). These results indicate that SOM performance is relatively insensitive to hyperparameter settings and does not require extensive fine-tuning.

### Simulation set 2 – Effects of divergence, migration, and sampling on lineage recovery

In Simulation set 2, we simulated two-population demographic scenarios across increasing divergence times and migration rates under three sample sizes with even and uneven sampling, assuming an effective population size of 250,000 diploid individuals per deme. *delimSOM* 2.0 correctly supported K = 1 at low divergence and shifted toward K = 2 with increasing divergence (Figure 2A). Without gene flow, this transition occurred at approximately 55,000 generations (Figure 2A; purple dashed line). Introducing gene flow progressively delayed the transition, and at the highest migration rate, only K = 1 was supported across the simulated divergence range. Reducing sample size or introducing uneven sampling delayed the transition from K = 1 to K = 2, indicating reduced power to detect divergence (Figure 2A–B; Supplementary Figures S5A–B, S6A–B).

**Figure 2:**
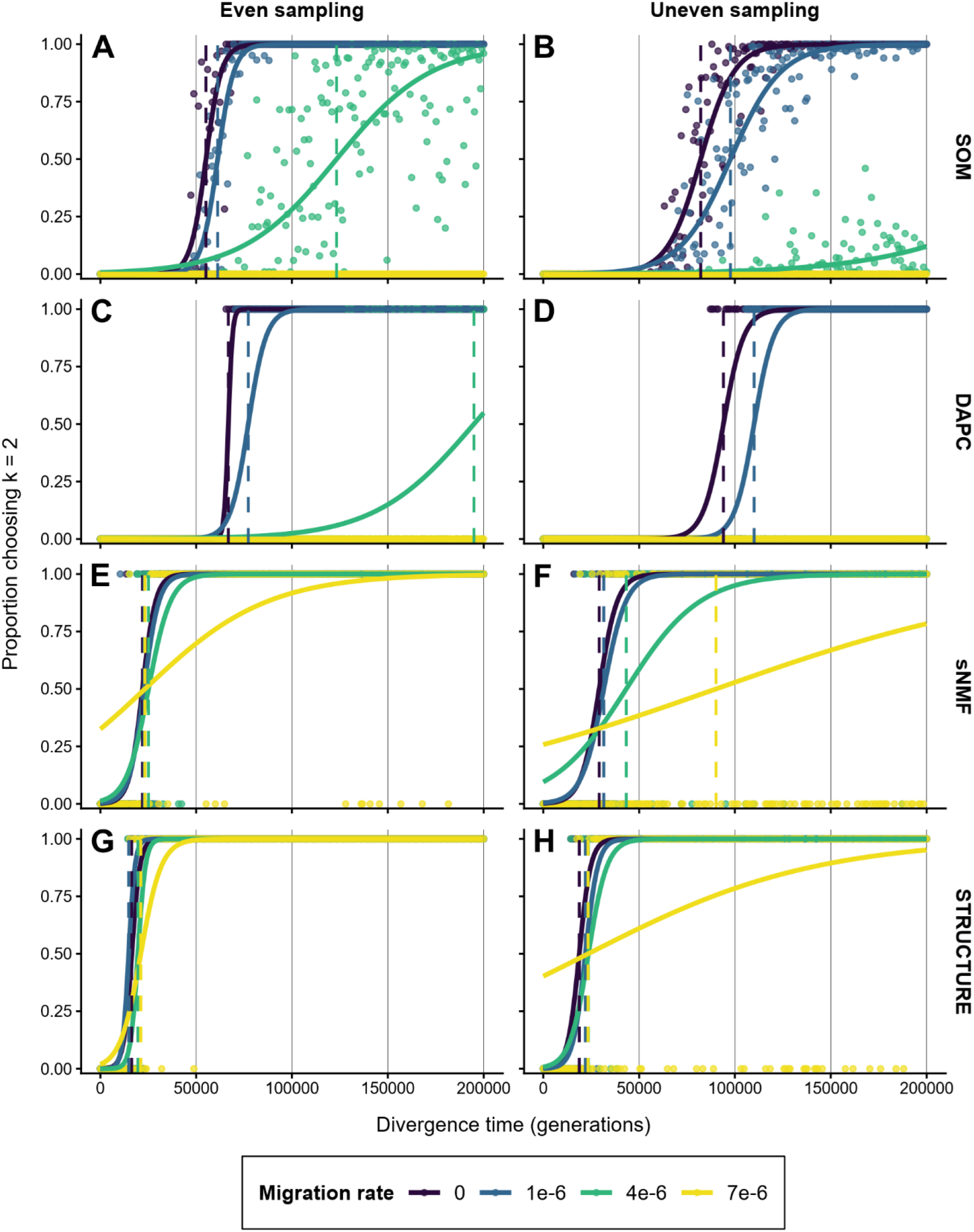
Effects of divergence time, migration rate, and uneven sampling on recovery of two simulated populations (K = 2) in simulation set 2. Datasets comprising 1,000 SNPs were simulated with gradually increasing divergence between two populations and analyzed using *delimSOM* 2.0 **(A, B)**, *DAPC* **(C, D)**, *sNMF* **(E, F)**, and *STRUCTURE* **(G, H)**. Left and right columns show even (24 individuals per population; 1:1 ratio) and uneven sampling (36 and 12 individuals per population; 3:1 ratio), respectively. Points show observed proportions selecting K = 2, solid colored lines show fitted probabilities from binomial logistic regression models, and dashed vertical lines mark the divergence time at which the fitted probability reaches 0.5.

At intermediate divergence along the simulated divergence continuum, many SOM replicates supported either K = 1 or K = 2 (Figure 2A; Supplementary Figures S5A, S6A). This mixed support for both K occurred even in the absence of gene flow, consistent with incomplete lineage differentiation at intermediate divergence, but was especially frequent under migration, where gene flow produced less discrete lineage boundaries. Evaluation of genetic differentiation between the simulated populations showed that transitions between K = 1 and K = 2 generally corresponded to an F_ST_ of approximately

0.11 (Supplementary Figure S7).

Comparison with three alternative methods revealed the same general effects of divergence, gene flow, sample size, and uneven sampling but differences in the timing of the transition from K = 1 to K = 2. Because K selection relied on method-specific thresholds, we refrain from interpreting transition points as fixed quantitative differences among methods but rather focus on general performance trends. *DAPC* produced patterns most similar to *delimSOM* 2.0, although its transition occurred at slightly higher divergence times (Figure 2C–D; Supplementary Figures S5C–D, S6C–D). In contrast, *sNMF* and particularly *STRUCTURE* supported K = 2 at substantially lower divergence times (Figure 2E–H; Supplementary Figures S5E–H, S6E–H).

### Empirical analyses recover supported species while avoiding over-splitting

Overall, *delimSOM* 2.0 recovered the number of species recognized by the original studies in several empirical systems, depending on the clustering and K-selection method (Figures 3 & 5; Supplementary Tables S4–S5 and Figures S8–S13). Recovery of delimited species was strongest in systems with fewer taxa and relatively large samples, whereas species-rich complexes with limited sampling per species were generally resolved into fewer, broader lineages. Furthermore, closely related and recently diverged species were often combined into a single lineage (e.g., Figure 3).

**Figure 3:**
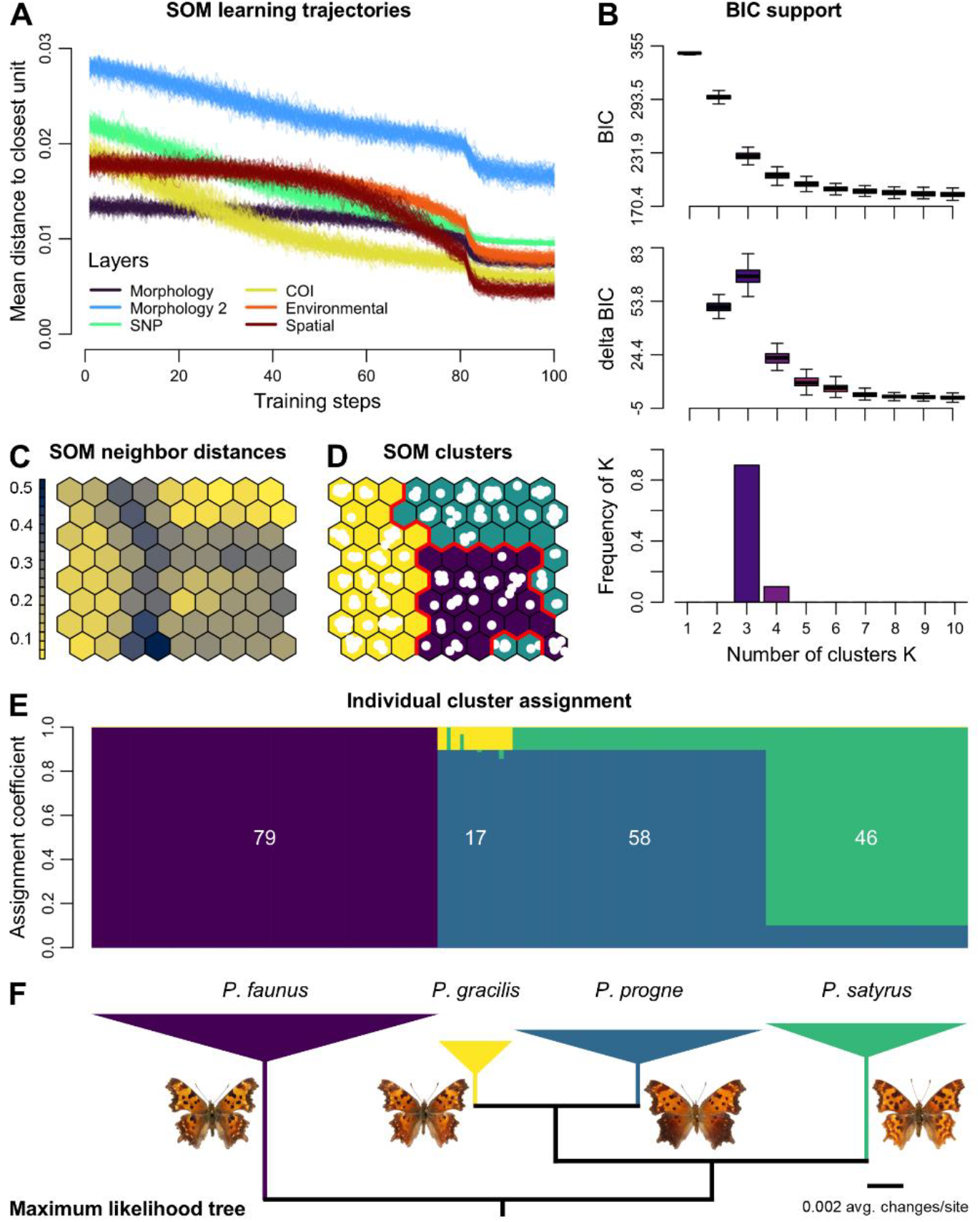
*delimSOM* 2.0 results from the empirical focus study comprising 200 *Polygonia* anglewing butterflies representing four recognized species across six layers (Dupuis et al., 2018). SOM analyses recovered three candidate lineages (K = 3), with *P. gracilis* and *P. progne* grouped into a single lineage. Morphology comprised a continuous layer with wing-color measurements and a categorical layer with wing characters. **A)** Learning trajectories showing mean distance to the closest map unit across training steps for each layer, where decreasing and stabilizing values indicate improved representation of the data and training convergence. **B)** Support for alternative K based on BIC values, changes in BIC between successive K values, and the frequency with which each K was selected across SOM replicates, with lower BIC indicating greater support and positive ΔBIC indicating improvement after adding a cluster. **C)** Neighbor distances among adjacent SOM units for a representative replicate, with dark areas showing larger distances between neighboring codebook vectors that indicate group boundaries (“ridges”) and light areas showing lower distances that form “valleys,” which indicate candidate lineages when separated by ridges. **D)** Recovered candidate lineages on the SOM map, with white points representing mapped individuals and red lines indicating lineage boundaries. **E)** *STRUCTURE*-like barplot of replicate-consensus assignment coefficients, with each bar representing one individual and numbers indicating individuals per species. **F)** Maximum-likelihood phylogeny from Dupuis et al. (2018) based on a 121,513-bp alignment from genotyping-by-sequencing, with colored triangles representing collapsed intraspecific branches (outgroups and other *Polygonia* not shown). Photo credits: Kim Davis, Mike Strangeland, and Andrew Warren.

**Figure 4:**
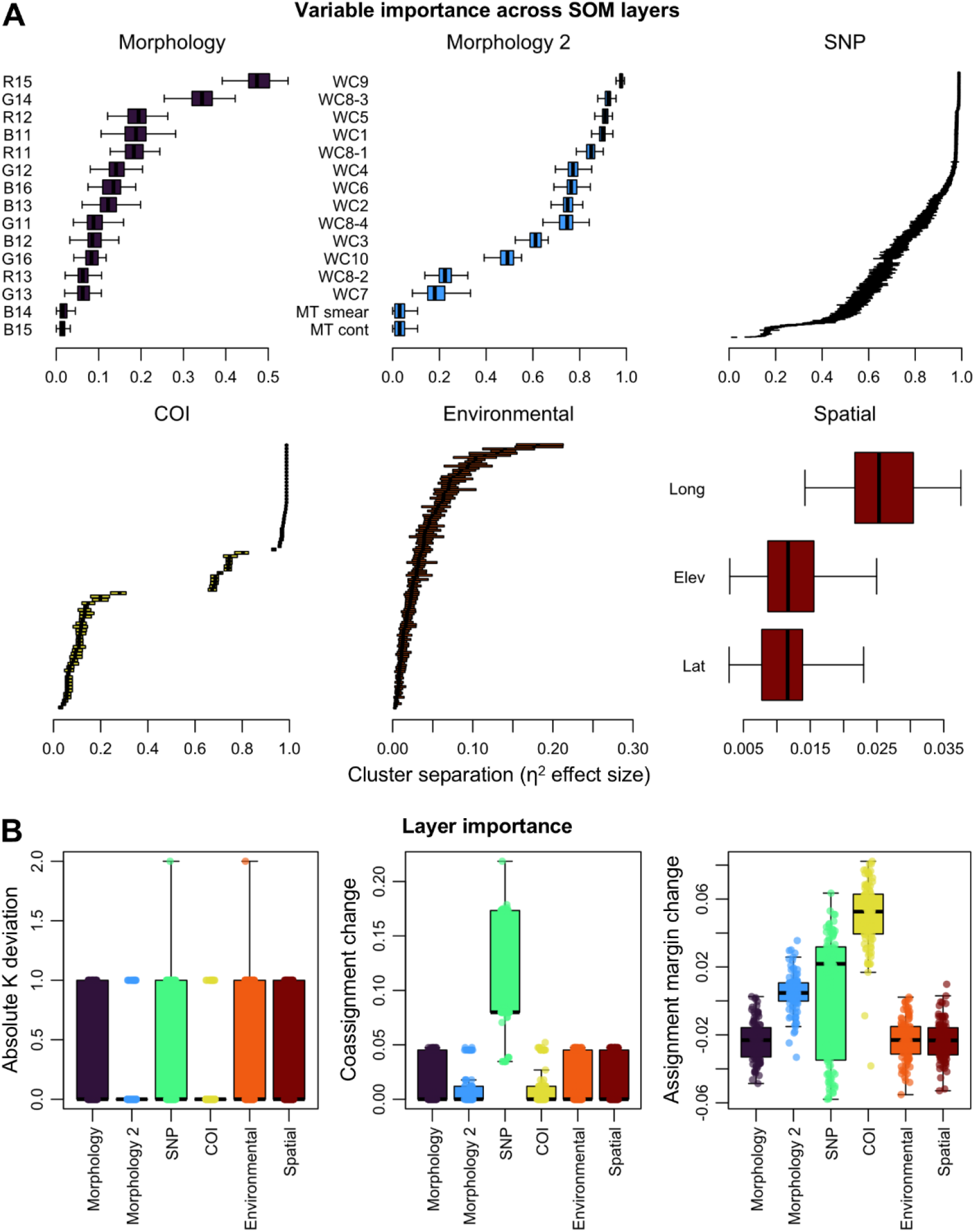
Variable and layer importance for the empirical focus study of 200 *Polygonia* anglewing butterflies (Dupuis et al., 2018). **A)** Variable importance across SOM layers based on an ANOVA-like η^2^ effect size representing the proportion of variation in each variable associated with the inferred clusters, with boxplots showing the distribution of η^2^ values across SOM replicates for each variable and higher values indicating stronger associations with cluster separation. **B)** Layer importance from leave-one-layer-out analyses measuring changes in the inferred number of clusters (left), cluster composition (which individuals are grouped together; middle), and confidence of individual cluster assignments (how much support for the best cluster decreases relative to the second-best cluster; right) after omitting each layer, with points representing replicate-matched comparisons and boxplots summarizing variation across SOM replicates. For all metrics, higher values indicate greater importance for inferring cluster structure. Morphology was represented by a continuous layer comprising RGB values for wing spots 11–16 (“Morphology”) and a categorical layer comprising wing characters (WC) 1–10 and two morphotypes (MT) (“Morphology_2”)

**Figure 5:**
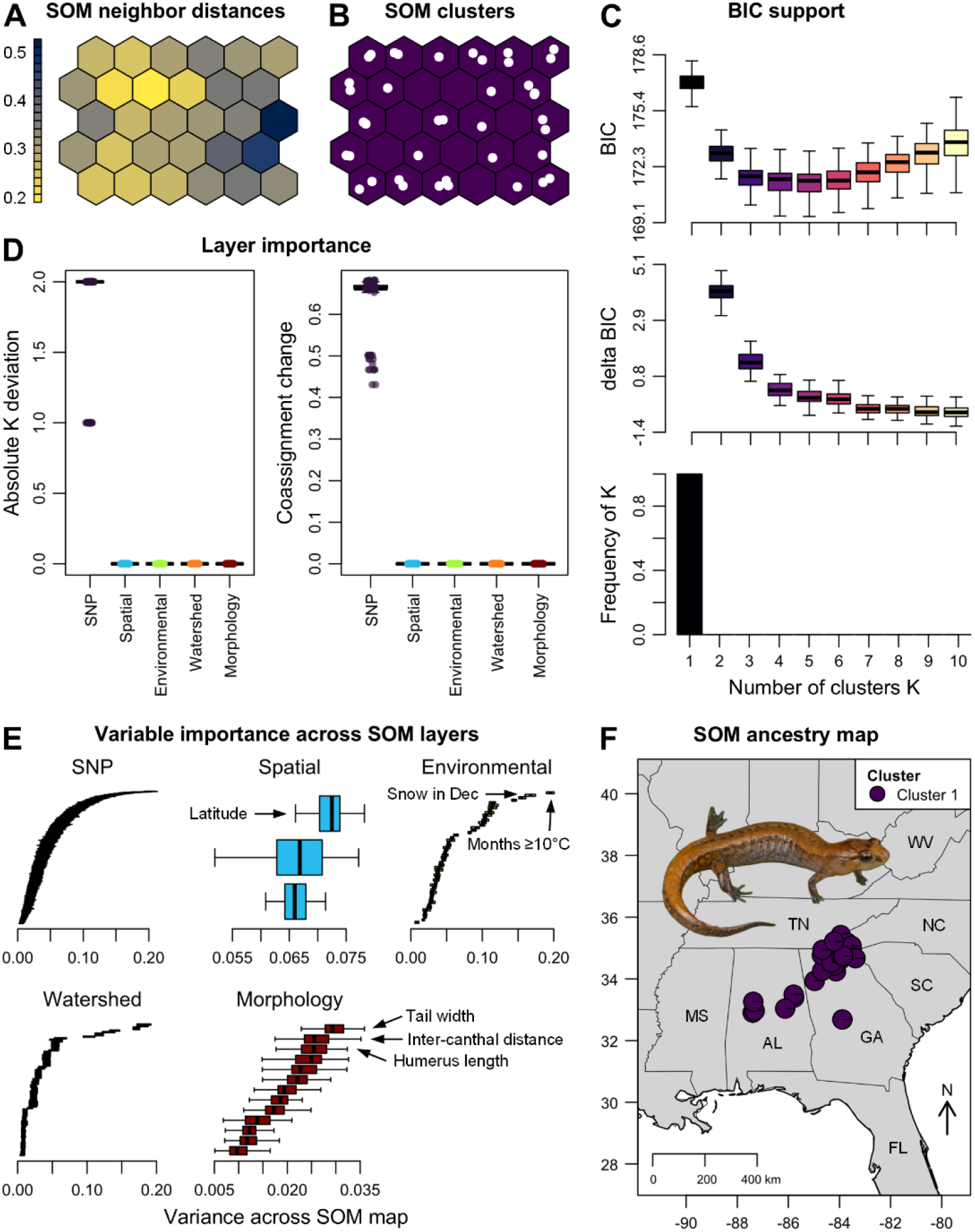
*delimSOM* 2.0 results from the empirical focus study of 40 *Desmognathus aeneus* seepage salamanders across five data layers (Pyron et al., 2024). SOM analyses recovered a single candidate lineage (K = 1). **A)** Neighbor distances among adjacent SOM units for a representative replicate, with dark areas showing larger distances between neighboring codebook vectors that indicate group boundaries (“ridges”) and light areas showing lower distances that form “valleys,” which indicate candidate lineages when separated by ridges. **B)** Recovered SOM clustering for the same replicate, with white points representing mapped individuals. **C)** Support for alternative K based on BIC values, changes in BIC between successive K values, and the frequency with which each K was selected across SOM replicates. **D)** Layer importance from leave-one-layer-out analyses measuring changes in the inferred number of clusters (left) and cluster composition (which individuals are grouped together; right) after omitting each layer, with points representing replicate-matched comparisons and boxplots summarizing variation across SOM replicates. **E)** Variable importance based on variance across the SOM map, with boxplots showing replicate-level map-variance distributions for individual variables (see Supplementary Table S7 for the most important variables). **F)** Species distribution in the southeastern USA, with replicate-consensus assignment coefficients showing all individuals assigned to the single inferred cluster. Photo credit: Max A. Seldes.

Although genomic layers consistently had much larger raw distance scales than other layers before SOM training (Supplementary Figure S14), our implemented layer weighting reduces the dominance of such layers during training. Consistent with this, layer-importance analyses indicated that many data types contributed substantially to delimitation outcomes. Genomic data were strongly associated with lineage separation in most systems, but other layers were also influential in many cases, particularly spatial and environmental layers (Supplementary Figures S8–S13 and Tables S6–S13).

In the first focal study of *Polygonia* anglewing butterflies, our method supported three clusters (Figure 3). Two clusters corresponded to *P. faunus* and *P. satyrus*, while the third grouped the closely related sister species *P. gracilis* and *P. progne* into a single lineage. Nevertheless, replicate-consensus assignment coefficients indicated that *P. gracilis* and *P. progne* were separated in 10% of SOM replicates, and hierarchical reanalysis also resolved them as distinct, indicating that *delimSOM* 2.0 detected a weak signal of differentiation within this combined lineage. No hierarchical subdivision was recovered within *P. faunus* or *P. satyrus*. Variable-level importance analyses identified the variables most strongly associated with separation among the inferred clusters, including key morphological characters (Figure 4A; Supplementary Table S6). Leave-one-layer-out analyses further showed that SNPs had the strongest effect on the inferred clustering, followed by wing-pattern characters and mitochondrial DNA, whereas environmental and spatial layers had weaker effects (Figure 4B).

In the second empirical study of the *D. aeneus* seepage salamander, our approach strongly supported a single candidate lineage (K = 1; Figure 5), despite pronounced geographic and genomic structure within the species. Leave-one-layer-out analysis identified the genomic layer as having the strongest effect on the delimitation outcome, with its removal substantially altering the number and composition of inferred clusters. Variable-level map-variance analyses indicated that the most important environmental variables were the number of months above 10°C, December precipitation as snow, degree days below 0°C, July climatic moisture deficit, and monthly burned area in winter; map variation for morphology was lower, with tail width, intercanthal width, and humerus length appearing most important (Supplementary Table S7).

## DISCUSSION

For decades, integrative species delimitation has been dominated by comparative and iterative approaches (Dayrat, 2005; Schlick-Steiner et al., 2010). The *delimSOM* framework moves the field toward fully integrative delimitation by incorporating heterogeneous sources of evidence to jointly inform candidate lineage hypotheses within a single unsupervised analysis. Building on SOMs that can capture complex non-linear multivariate structure (Kohonen, 1998), it leverages ML to synthesize multidimensional evidence of lineage divergence (Derkarabetian et al., 2019; Karbstein et al., 2024; Kösters et al., 2025). Our approach does not assume population-genetic models such as Hardy–Weinberg equilibrium, nor does it require an explicit evolutionary or demographic model or predefined species assignments (Pyron, 2023). Across simulations and empirical examples, *delimSOM* 2.0 generally recovered well-supported candidate lineages, retained K = 1 when differentiation reflected population rather than species structure, responded predictably to divergence, gene flow, and sampling, and remained relatively robust to missing data.

### Performance across simulated and empirical scenarios

*delimSOM* 2.0 performed well in recovering the expected number of candidate lineages across our simulation and empirical analyses, particularly when lineage numbers were relatively low and within-lineage sample sizes were large. Nevertheless, recovery declined as the number of lineages increased and sample sizes decreased. Uneven sampling also reduced the power to recover the expected structure. Thus, we recommend applying our method to systems with few candidate lineages (approximately K ≤ 4) and at least fifteen individuals per lineage to obtain robust conclusions. Species-rich and sparsely or unevenly sampled complexes require more cautious interpretation or additional analyses.

Our simulation and focal empirical analyses demonstrate the desirable tendency of *delimSOM* 2.0 to require relatively strong differentiation before supporting separate lineages. In simulation set 2, K = 1 was retained at low differentiation and under high migration, while mixed support for K = 1 and K = 2 occurred around the transition between population structure and lineage divergence (Figure 2). In *Polygonia*, closely related sister species were initially merged into one lineage but separated in some SOM replicates and through hierarchical reanalysis (Figure 3). In *D. aeneus*, our method supported K = 1 (Figure 5) despite pronounced genomic and geographic structure, consistent with the interpretation of this structure as intraspecific differentiation maintained by migration and admixture (Pyron et al., 2024). This distinction is particularly important in geographically structured taxa, where isolation by distance and restricted dispersal can generate pronounced genetic differentiation without corresponding to species boundaries (Chambers et al., 2025; Chan et al., 2017, 2020). This conservative behavior is intentional: *delimSOM* 2.0 was designed to retain K = 1 when evidence for multiple lineages is weak, an important advantage over approaches that cannot evaluate K = 1 or favor K > 1 even without clear structure (Janes et al., 2017; Pyron et al., 2023). This mitigates the risk of interpreting population structure as species-level divergence, thereby reducing over-splitting and inflation of biodiversity estimates (Dufresnes et al., 2023; Sukumaran & Knowles, 2017).

Compared with *DAPC, sNMF*, and *STRUCTURE*, our method supported K = 2 at substantially later divergence times than *sNMF* and *STRUCTURE*, but slightly earlier than *DAPC*. The broadly comparable behavior of *delimSOM* 2.0 and *DAPC* is promising, suggesting that our framework performs similarly to this well-established multivariate method (Jombart et al., 2010), with both requiring stronger species-level differentiation before supporting K = 2. By contrast, the earlier transitions of *sNMF* and *STRUCTURE* are consistent with their model-based design to detect fine-scale population structure (Frichot et al., 2014; Pritchard et al., 2000).

A further strength of *delimSOM* 2.0 is that it explicitly quantifies uncertainty at both the lineage- and individual-assignment levels: replicate support can be distributed among alternative values of K, while replicate-consensus assignment coefficients quantify how consistently each individual is assigned to the inferred clusters (e.g., Figure 3). Mixed support among K values or intermediate individual assignments may indicate weak differentiation, recent divergence, admixture, or discordance among data sources. For closely related or recently diverged taxa, we recommend hierarchical reanalysis of broadly supported clusters and inspection of support curves and replicate-consensus assignment coefficients to identify weaker differentiation (Janes et al., 2017).

### Clustering and SOM parameterization

Determining the optimal number of clusters remains challenging because no single clustering or K-selection approach performs best across all data structures (Omran et al., 2007; Rodriguez et al., 2019).

We evaluated six clustering and K-selection approaches using simulated and empirical datasets. This showed that the two k-means-based BIC approaches provided the best balance of expected-lineage recovery, assignment accuracy, and computational efficiency. Because the elbow approach performed well in the original *delimSOM* implementation (Pyron, 2023; Pyron et al., 2023) and is conceptually related to the well-established *diffNgroup* elbow criterion in *DAPC* (Jombart et al., 2010), we recommend using k-means with BIC-elbow selection.

In *delimSOM* 2.0, we revised the original elbow-detection procedure to explicitly compare K = 1 and K

= 2, retaining K = 1 when the improvement from adding a second cluster is below the specified BIC threshold. This conservative safeguard reduces the risk that weak population structure or stochastic variation is interpreted as support for multiple lineages. However, the K = 7 simulations showed that this safeguard can prevent the automatic recovery of more complex structure when the initial improvement from K = 1 to K = 2 is small, even when a clear elbow occurs at the expected K. Users should therefore inspect the BIC curve rather than relying solely on the automatically selected K. For such cases or when a particular K value is of interest, users can rerun the clustering and K-selection step with K fixed to that value.

Alternative clustering methods may be useful when larger numbers of lineages or more complex cluster geometries are expected. GMM with BIC-threshold selection also performed well and represents a suitable complementary approach, although it required longer runtimes. By contrast, HDBSCAN and OPTICS were less reliable, and hierarchical clustering had prohibitively long runtimes, making these methods less suitable.

### Toward fully integrative species delimitation

Species delimitation approaches commonly infer genomic, phenotypic, or environmental structure through separate analyses and then reconcile the resulting partitions qualitatively (Schlick-Steiner et al., 2010; Yeates et al., 2011). *delimSOM* instead retains each data type as a distinct layer and integrates them through a shared competitive-learning process. The contributions of non-genomic layers across our empirical systems highlight the value of including additional layers beyond genomics as complementary evidence of lineage divergence (Edwards & Knowles, 2014; Padial et al., 2010; Schlick-Steiner et al., 2010). Since variable-and layer-importance analyses identify the biological axes most strongly associated with inferred lineage structure, they can also generate hypotheses about the processes contributing to lineage divergence: strong genomic differentiation may be consistent with restricted gene flow, morphological differentiation with phenotypic divergence, environmental differentiation with niche divergence, and behavioral differentiation with potential isolating mechanisms (de Queiroz, 2007; Raxworthy et al., 2007; van Elst et al., 2024). Our fully integrative framework also reduces analytical complexity and subjectivity, in line with repeated calls for more objective and standardized species delimitation (Karbstein et al., 2024; Schlick-Steiner et al., 2010). By contrast, comparative and iterative approaches require multiple specialized analyses across data types, demanding substantial time and methodological expertise, while reconciling separate results may depend on researcher judgment (Schlick-Steiner et al., 2010; Yeates et al., 2011).

One constraint is that individuals must be represented across all layers. Genetic and morphological data are frequently collected from different field or museum specimens. Behavioral, reproductive, and life-history observations may be available only at population or species level, while environmental data are often extracted from citizen-science occurrence records that do not correspond to the individuals used for genomic or morphological analyses. Similarly, older studies often provide only species means or diagnostic characters rather than individual-level matrices. Intersecting independently collected datasets can therefore cause substantial sample loss and limit the number of useable datasets for analysis, as evidenced in our empirical analyses. Consequently, *delimSOM* is suited to prospective studies where genomic, phenotypic, ecological, and spatial information is collected for the same specimens. Future expansions could examine the possibility of merging heterogeneous datasets and allowing missingness across layers.

Nonetheless, *delimSOM* 2.0 can accommodate substantial missingness within individuals. In Simulation set 1, performance declined only marginally up to approximately 50% missingness and more sharply at higher levels. The robustness of *delimSOM* is facilitated by how missing data are handled during SOM training: after samples and variables with high missingness are filtered, best-matching units are identified using only observed dimensions, and codebook updates are restricted to those dimensions, allowing incomplete observations to contribute without global imputation across heterogeneous data (Samad & Harp, 1992). Some other integrative approaches also accommodate missingness, including those of Guillot et al. (2012) and Solís-Lemus et al. (2015). However, many population-genetic workflows rely on substantially lower levels of missing data or prior filtering, including *STRUCTURE* (Pritchard et al., 2000) and *sNMF* (Frichot et al., 2014). Similarly, *DAPC* (Jombart et al., 2010) and PCA require prior removal or mean imputation of incomplete observations.

## Acknowledgments

We thank Patrick J. Krug (California State University, USA) for assistance with data compilation for the *Elysia* empirical case study, Tobias van Elst (ETH Zürich, Switzerland) for assistance with reprocessing RADseq data to expand genomic coverage in the *Microcebus* reanalysis, and the University of Kentucky Center for Computational Sciences and Information Technology Services Research Computing for support and access to the Morgan Compute Cluster.

## References

Carstens, B. C., Pelletier, T. A., Reid, N. M., & Satler, J. D. (2013). How to fail at species delimitation. Molecular Ecology, 22(17), 4369–4383. 10.1111/mec.12413

Chambers, E. A., Lara-Tufiño, J. D., Campillo-García, G., Cisneros-Bernal, A. Y., Dudek, D. J., León-Règagnon, V., Townsend, J. H., Flores-Villela, O., & Hillis, D. M. (2025). Distinguishing species boundaries from geographic variation. Proceedings of the National Academy of Sciences, 122(19). 10.1073/pnas.2423688122

Chan, K. O., Alexander, A. M., Grismer, L. L., Su, Y., Grismer, J. L., Quah, E. S. H., & Brown, R. M. (2017). Species delimitation with gene flow: A methodological comparison and population genomics approach to elucidate cryptic species boundaries in Malaysian Torrent Frogs. Molecular Ecology, 26(20), 5435–5450. 10.1111/mec.14296

Chan, K. O., Hutter, C. R., Wood, P. L., Grismer, L. L., Das, I., & Brown, R. M. (2020). Gene flow creates a mirage of cryptic species in a Southeast Asian spotted stream frog complex. Molecular Ecology, 29(20), 3970–3987. 10.1111/mec.15603

Dayrat, B. (2005). Towards integrative taxonomy. Biological Journal of the Linnean Society, 85(3), 407–415. 10.1111/j.1095-8312.2005.00503.x

de Queiroz, K. (2007). Species concepts and species delimitation. Systematic Biology, 56(6), 879–886. 10.1080/10635150701701083

Derkarabetian, S., Castillo, S., Koo, P. K., Ovchinnikov, S., & Hedin, M. (2019). A demonstration of unsupervised machine learning in species delimitation. Molecular Phylogenetics and Evolution, 139, 106562. 10.1016/j.ympev.2019.106562

Dufresnes, C., Poyarkov, N., & Jablonski, D. (2023). Acknowledging more biodiversity without more species. Proceedings of the National Academy of Sciences, 120(40). 10.1073/pnas.2302424120

Dupuis, J. R., Peigler, R. S., Geib, S. M., & Rubinoff, D. (2018). Phylogenomics supports incongruence between ecological specialization and taxonomy in a charismatic clade of buck moths. Molecular Ecology, 27(22), 4417–4429. 10.1111/mec.14883

Edwards, D. L., & Knowles, L. L. (2014). Species detection and individual assignment in species delimitation: Can integrative data increase efficacy? Proceedings of the Royal Society B: Biological Sciences, 281(1777), 20132765. 10.1098/rspb.2013.2765

Excoffier, L., Marchi, N., Marques, D. A., Matthey-Doret, R., Gouy, A., & Sousa, V. C. (2021). fastsimcoal2: Demographic inference under complex evolutionary scenarios. Bioinformatics, 37(24), 4882–4885. 10.1093/bioinformatics/btab468

Frichot, E., Mathieu, F., Trouillon, T., Bouchard, G., & François, O. (2014). Fast and efficient estimation of individual ancestry coefficients. Genetics, 196(4), 973–983. 10.1534/genetics.113.160572

Guillot, G., Renaud, S., Ledevin, R., Michaux, J., & Claude, J. (2012). A unifying model for the analysis of phenotypic, genetic, and geographic data. Systematic Biology, 61(6), 897–911. 10.1093/sysbio/sys038

Janes, J. K., Miller, J. M., Dupuis, J. R., Malenfant, R. M., Gorrell, J. C., Cullingham, C. I., & Andrew, R. L. (2017). The K= 2 conundrum. Molecular Ecology, 26(14), 3594–3602. 10.1111/mec.14187

Jombart, T., Devillard, S., & Balloux, F. (2010). Discriminant analysis of principal components. A new method for the analysis of genetically structured populations. BMC Genetics, 11(1), 94. 10.1186/1471-2156-11-94

Karbstein, K., Kösters, L., Hodač, L., Hofmann, M., Hörandl, E., Tomasello, S., Wagner, N. D., Emerson, B. C., Albach, D. C., Scheu, S., Bradler, S., de Vries, J., Irisarri, I., Li, H., Soltis, P., Mäder, P., & Wäldchen, J. (2024). Species delimitation 4.0: Integrative taxonomy meets artificial intelligence. Trends in Ecology & Evolution, 39(8), 771–784. 10.1016/j.tree.2023.11.002

Kohonen, T. (1998). The self-organizing map. Neurocomputing, 21(1–3), 1–6. 10.1016/S0925-2312(98)00030-7

Kohonen, T. (2014). MATLAB implementations and applications of the self-organizing map. Unigrafia Oy.

Kösters, L. M., Karbstein, K., Hofmann, M., Hodač, L., Mäder, P., & Wäldchen, J. (2025). Data fusion for integrative species identification using deep learning. Systematic Biology. 10.1093/sysbio/syaf026

Krug, P. J., Trathen, D. Y., Retana, J., Rico, D. M., Bronola, J., & Ellingson, R. A. (2026). Integrating host use and dispersal ability with species delimitation to unravel a cryptic radiation of photosynthetic sea slugs. Biological Journal of the Linnean Society, 147(3). 10.1093/biolinnean/blaf111

Omran, M. G. H., Engelbrecht, A. P., & Salman, A. (2007). An overview of clustering methods. Intelligent Data Analysis, 11(6), 583–605. 10.3233/IDA-2007-11602

Oury, N., Noël, C., Mona, S., Aurelle, D., & Magalon, H. (2023). From genomics to integrative species delimitation? The case study of the Indo-Pacific Pocillopora corals. Molecular Phylogenetics and Evolution, 184, 107803. 10.1016/j.ympev.2023.107803

Padial, J. M., Miralles, A., De la Riva, I., & Vences, M. (2010). The integrative future of taxonomy. Frontiers in Zoology, 7(1), 16. 10.1186/1742-9994-7-16

Pei, J., Chu, C., Li, X., Lu, B., & Wu, Y. (2018). CLADES: A classification-based machine learning method for species delimitation from population genetic data. Molecular Ecology Resources, 18(5), 1144–1156. 10.1111/1755-0998.12887

Pritchard, J. K., Stephens, M., & Donnelly, P. (2000). Inference of population structure using multilocus genotype data. Genetics, 155(2), 945–959. 10.1093/genetics/155.2.945

Pyron, R. A. (2023). Unsupervised machine learning for species delimitation, integrative taxonomy, and biodiversity conservation. Molecular Phylogenetics and Evolution, 189, 107939. 10.1016/j.ympev.2023.107939

Pyron, R. A., Kakkera, A., Beamer, D. A., & O’Connell, K. A. (2024). Discerning structure versus speciation in phylogeographic analysis of seepage salamanders (Desmognathus aeneus) using demography, environment, geography, and phenotype. Molecular Ecology, 33(2). 10.1111/mec.17219

Pyron, R. A., O’Connel, K. A., Lamb, J. Y., & Beamer, D. A. (2022). A new, narrowly endemic species of swamp-dwelling dusky salamander (Plethodontidae: Desmognathus) from the Gulf Coastal Plain of Mississippi and Alabama. Zootaxa, 5133(1), 53–82. 10.11646/zootaxa.5133.1.3

Pyron, R. A., O’Connell, K. A., Duncan, S. C., Burbrink, F. T., & Beamer, D. A. (2023). Speciation hypotheses from phylogeographic delimitation yield an integrative taxonomy for seal salamanders (Desmognathus monticola). Systematic Biology, 72(1), 179–197. 10.1093/sysbio/syac065

Raxworthy, C. J., Ingram, C. M., Rabibisoa, N., & Pearson, R. G. (2007). Applications of ecological niche modeling for species delimitation: A review and empirical evaluation using day geckos (Phelsuma) from Madagascar. Systematic Biology, 56(6), 907–923. 10.1080/10635150701775111

Rodriguez, M. Z., Comin, C. H., Casanova, D., Bruno, O. M., Amancio, D. R., Costa, L. da F., & Rodrigues, F. A. (2019). Clustering algorithms: A comparative approach. PLOS ONE, 14(1), e0210236. 10.1371/journal.pone.0210236

Samad, T., & Harp, S. (1992). Self–organization with partial data. Network: Computation in Neural Systems, 3(2), 205–212. 10.1088/0954-898X/3/2/008

Schlick-Steiner, B. C., Steiner, F. M., Seifert, B., Stauffer, C., Christian, E., & Crozier, R. H. (2010). Integrative taxonomy: A multisource approach to exploring biodiversity. Annual Review of Entomology, 55(1), 421–438. 10.1146/annurev-ento-112408-085432

Schönberger, D., MacDonald, Z. G., Schmidt, B. C., & Dupuis, J. R. (2026). NicheDiv: A DAPC framework to quantify niche divergence across highly multivariate environmental space. BioRxiv. 10.64898/2026.06.19.733388

Smith, M. L., & Carstens, B. C. (2020). Process-based species delimitation leads to identification of more biologically relevant species. Evolution, 74(2), 216–229. 10.1111/evo.13878

Solís-Lemus, C., Knowles, L. L., & Ané, C. (2015). Bayesian species delimitation combining multiple genes and traits in a unified framework. Evolution, 69(2), 492–507. 10.1111/evo.12582

Spriggs, E. L., Eaton, D. A. R., Sweeney, P. W., Schlutius, C., Edwards, E. J., & Donoghue, M. J. (2019). Restriction-site-associated DNA sequencing reveals a cryptic Viburnum species on the North American coastal plain. Systematic Biology, 68(2), 187–203. 10.1093/sysbio/syy084

Stankowski, S., & Ravinet, M. (2021). Defining the speciation continuum. Evolution, 75(6), 1256– 1273. 10.1111/evo.14215

Sukumaran, J., & Knowles, L. L. (2017). Multispecies coalescent delimits structure, not species. Proceedings of the National Academy of Sciences, 114(7), 1607–1612. 10.1073/pnas.1607921114

van Elst, T., Sgarlata, G. M., Schüßler, D., Tiley, G. P., Poelstra, J. W., Scheumann, M., Blanco, M. B., Aleixo-Pais, I. G., Rina Evasoa, M., Ganzhorn, J. U., Goodman, S. M., Hasiniaina, A. F., Hending, D., Hohenlohe, P. A., Ibouroi, M. T., Iribar, A., Jan, F., Kappeler, P. M., Le Pors, B., … Salmona, J. (2024). Integrative taxonomy clarifies the evolution of a cryptic primate clade. Nature Ecology & Evolution. 10.1038/s41559-024-02547-w

Wehrens, R., & Kruisselbrink, J. (2018). Flexible self-organizing maps in kohonen 3.0. Journal of Statistical Software, 87(7). 10.18637/jss.v087.i07

Yang, B., Zhang, Z., Yang, C.-Q., Wang, Y., Orr, M. C., Wang, H., & Zhang, A.-B. (2022). Identification of species by combining molecular and morphological data using convolutional neural networks. Systematic Biology, 71(3), 690–705. 10.1093/sysbio/syab076

Yeates, D. K., Seago, A., Nelson, L., Cameron, S. L., Joseph, L., & Trueman, J. W. H. (2011). Integrative taxonomy, or iterative taxonomy? Systematic Entomology, 36(2), 209–217. 10.1111/j.1365-3113.2010.00558.x

