## Supplement for "Fully integrative species delimitation with machine learning and diverse data types in *delimSOM* 2.0"

### Supplementary Materials for:

#### Fully integrative species delimitation with machine learning: Using self-organizing maps to infer candidate lineages from genetic, phenotypic, spatial, and environmental data in *delimSOM* 2.0

**Daniel Schönberger[1], R. Alexander Pyron[2], Julian R. Dupuis[1]**

[1]Department of Entomology, University of Kentucky, Lexington, Kentucky, USA

[2]Department of Biological Sciences, George Washington University, Washington, DC, USA

**This PDF includes:**

Supplementary Methods: M1–M9
Supplementary Figures: S1–S14
Supplementary Tables: S1–S15
Supplementary References

### Supplementary Methods

#### M1: Lay summary of the *delimSOM* framework

A Self-Organizing Map (SOM) is a way of turning a complicated table of measurements into a two-dimensional grid map that is easier to understand. The grid consists of cells, where each cell stores a representative "average" pattern (called a codebook vector) for the variables in each data layer. Each specimen is then compared with all these grid cells, and it is assigned to the cell whose pattern is most like its own overall combination of measurements. This cell is called the best-matching unit. During training, the winning and nearby cells on the map are adjusted to become slightly more similar to that specimen. After many repeated updates across all specimens, the grid becomes organized so that similar specimens tend to occupy the same or neighboring parts of the map, whereas more different specimens tend to fall farther apart. SOM does not preserve distances exactly, but local similarity relationships. Another useful feature is that the map summarizes many individual observations into a smaller set of representative local averages, which can reduce sensitivity to noise and make broad structure easier to identify. It also tends to allocate more representative cells to densely sampled regions of multivariate data space than to sparse regions. These attributes make SOM especially helpful for complex biological datasets in which relationships may be nonlinear, partly conflicting across variables, or distributed across different data types. Different types of information (e.g., genetics, morphology, climate) are kept as separate "layers" but contribute jointly to the same map. Their contributions are balanced so that a dataset with many variables, such as genomic data, does not outweigh a smaller dataset, such as morphology or environment. After the map has been learned, the representative map cells (codebook vectors) themselves are grouped into clusters, and the optimal number of clusters (K) can be evaluated. These clusters are then interpreted as candidate species. Because the map can vary each time it is trained due to random initialization, many SOM replicates are run, and their results are then combined to identify the groupings that are most consistently recovered.

#### M2: *delimSOM* framework

Our *delimSOM* 2.0 *R* package implements a SOM framework that supports fully integrative species delimitation using genomic, phenotypic, environmental, spatial, and other multivariate data. The approach consists of two main steps. First, the original high-dimensional data are represented by SOM codebook vectors arranged on a two-dimensional topology-preserving grid. These codebook vectors are learned local representations within the multivariate space (or, in simple words, local "averages"). Second, those codebook vectors are clustered into groups that are interpreted as candidate species. This two-stage strategy follows the classical use of SOMs as competitive, unsupervised neural networks that represent high-dimensional observations by codebook vectors arranged on a low-dimensional grid, thereby emphasizing local neighborhood preservation rather than global distance preservation (Kohonen, 1990, 1998, 2001; Wehrens & Buydens, 2007). Because map training and clustering are unsupervised, predefined group assignments do not need to be provided, and the candidate delimitation structure is inferred from multivariate similarities among observations and the resulting codebook vectors (Pyron, 2023; Pyron et al., 2023). During training, each observation is matched to the map unit whose codebook vector is most similar to it, which is called the best-matching unit. The best-matching unit and nearby units are updated so that similar observations become organized into neighboring regions of the map. In this framework, neighboring map units contain relatively similar codebook vectors, whereas more distant regions of the map summarize broader differences in multivariate structure (Kohonen, 1998; Wehrens & Buydens, 2007). Because SOM codebook vectors are learned local representations rather than individual observations, the SOM acts as a regularized representation that can reduce sensitivity to noise and individual-level irregularities while preserving major topological relationships (Kohonen, 2001, 2014; Vesanto & Alhoniemi, 2000). This makes SOMs especially useful for complex biological datasets in which structure may be nonlinear, subtle, partially conflicting across variables, or unevenly distributed across different data types (Lek et al., 1996; Lek & Guégan, 2000; Zamani et al., 2013). SOMs are also well-suited to integrative delimitation because multilayer SOMs can accommodate heterogeneous data types through layer-specific distance functions and weighting, allowing signals from genetics, phenotype, environment, and space to contribute jointly to the learned map without requiring a single explicit generative model for all data types (Pyron, 2023; Wehrens & Buydens, 2007; Wehrens & Kruisselbrink, 2018). The subsequent clustering of codebook vectors provides a second level of abstraction by identifying major groups (Akman et al., 2019; Kohonen, 2013; Vesanto & Alhoniemi, 2000), which are then interpreted as candidate species (Pyron, 2023; Pyron et al., 2023). Because SOM training depends on the stochastic initialization of the grid map, multiple independent replicate maps are trained, and the resulting delimitation assignments are summarized across replicates (Pyron, 2023; Pyron et al., 2023).

**M3: Processing of data and preparation of SOM input matrices**

*delimSOM* 2.0 provides several preprocessing functions for preparing raw input data for SOM training and clustering. For genomic inputs, including SNP dosage data and aligned sequence data such as COI, preprocessing is performed by our *process.SNP.data.SOM* function (Supplementary Methods M3.1–3.5). For categorical non-genetic inputs (e.g., habitat type, host type, morphological characters), preprocessing can be performed with our *make.cols.binary.SOM* function (Supplementary Methods M3.6), which converts categorical variables into binary matrices. For numeric matrices (e.g., climatic or morphological data), our *remove.lowCV.multicollinearity.SOM* function removes variables with low variation and high correlations (Supplementary Methods M3.7). These preprocessing functions facilitate data handling and maximize comparability among individuals and layers. The normalization of the processed matrices is handled directly by our *train.SOM* function (Supplementary Methods M4).

1. **Input and setup for genetic data.** Our *process.SNP.data.SOM* function facilitates the preprocessing of commonly used genetic data formats into matrices suitable for SOM training. The function accepts one of eight inputs: 1) a VCF file path (*vcf.path* argument), 2) a genind object (*genind.input* argument), 3) a genlight object (*genlight.input* argument), 4) a numeric SNP dosage matrix or data frame (*snp.matrix.input* argument), 5) a PLINK .raw dosage file path (*plink.raw.path* argument), 6) a NEXUS alignment file path (*nexus.path* argument), 7) a FASTA alignment file path (*fasta.path* argument), or 8) a PHYLIP alignment file path (*phylip.path* argument). Additional arguments control the PHYLIP reading format (*phylip.format* argument; *"sequential"* or *"interleaved"*), PLINK metadata columns to exclude (*plink.raw.metadata.columns* argument), the ploidy of numeric SNP dosage matrices and genlight inputs (*snp.matrix.ploidy* argument), the treatment of two-base IUPAC ambiguity codes in sequence alignments (*alignment.ambiguity.mode* argument; *"heterozygote"* or *"missing"*), the maximum proportions of missing data permitted per locus during an initial lenient and final stringent pass (*missing.loci.cutoff.lenient* and *missing.loci.cutoff.final* arguments), the maximum proportion of missing data allowed per individual (*missing.individuals.cutoff* argument), whether singleton and invariant loci are removed (*singleton.loci.filter* and *invariant.loci.filter* arguments), and whether filtering summaries are printed (*verbose* argument).

2. **Genetic data import.** If NEXUS, FASTA, or PHYLIP alignments are provided, the function reads the aligned sequences, converts them into a sample-by-site character matrix, standardizes case, assigns row names from the sequence identifiers, and standardizes missing and ambiguous symbols. The symbols *"?"*, *"-"*, *"N"*, *"B"*, *"D"*, *"H"*, *"V"*, and *"X"* are treated as missing. When *alignment.ambiguity.mode = "heterozygote"* (the recommended default), the two-base IUPAC

codes *"R"*, *"Y"*, *"S"*, *"W"*, *"K"*, and *"M"* are interpreted as heterozygous genotypes when they match the two alleles at a biallelic site. When *alignment.ambiguity.mode = "missing"*, these codes are treated as missing. For NEXUS input, reading is performed with the *read.nexus.data* function in *ape*. Because *"."* can represent a NEXUS match character rather than true missing data, the function first expands *"."* relative to the first sequence and then treats any unresolved *"."* symbols as missing. FASTA and PHYLIP alignments are read with *read.dna* in *ape*, using *format = "fasta"* for FASTA files and the user-supplied *phylip.format* value for PHYLIP files. If a VCF file is provided, the function imports it via the *read.vcfR* and *extract.gt* functions in the *vcfR v.1.16 R* package (Knaus & Grünwald, 2017). This VCF processing retains single-base biallelic variants, extracts diploid genotype strings, standardizes phased genotype separators, treats fully or partially missing genotypes as missing values, and recodes supported genotypes as numeric dosages. If unsupported genotype coding is encountered, the function falls back to the genind conversion. If a genind object is supplied directly, the VCF import step is skipped. If a genlight object is supplied, it is converted to a dosage matrix while preserving individual and locus names where available. For numeric SNP dosage matrices, ploidy is specified with *snp.matrix.ploidy* (1 = haploid, 2 = diploid; default = 2) rather than inferred from the observed dosage values. For *genind* inputs, ploidy is obtained from the object, whereas PLINK *.raw* and standard VCF inputs are treated as diploid. If a PLINK .raw file is supplied, the columns specified by *plink.raw.metadata.columns* are excluded and the remaining columns are interpreted as SNP dosage variables.

3. **Biallelic encoding.** Our *process.SNP.data.SOM* function processes genetic datasets to retain only biallelic loci. Multiallelic loci are removed rather than expanded into a number of allele-dosage columns equal to one fewer than the number of alleles because such an encoding would cause a single biological locus to contribute multiple variables and could make the resulting representation and distances depend on which allele was selected as the omitted reference state. This could distort distance calculations, particularly for binary or Tanimoto-based comparisons. Standard haploid dosage data are encoded as 0/1, and standard diploid genotype data are encoded as 0/1/2. For each retained biallelic site in a sequence alignment, the alphabetically first allele is encoded as 0 and the second allele as 2. When *alignment.ambiguity.mode = "heterozygote"*, a matching two-base IUPAC ambiguity code is encoded as heterozygote dosage 1. When *alignment.ambiguity.mode = "missing"*, ambiguity codes are treated as missing.

4. **Locus and individual missingness filtering.** After allele-state processing and biallelic filtering, loci and individuals are filtered in a staged manner. The function first applies a lenient locus-missingness filter (*missing.loci.cutoff.lenient* argument; default: *0.7*), then removes individuals exceeding the individual missingness threshold (*missing.individuals.cutoff* argument; default: *0.5*), optionally removes singleton loci (*singleton.loci.filter = TRUE*), then applies the stricter locus-missingness filter (*missing.loci.cutoff.final* argument; default: *0.5*), and finally optionally removes invariant loci (*invariant.loci.filter = TRUE*). The singleton filter is placed after individual filtering because singletons are defined relative to the retained individuals. Removing singleton loci does not affect the missing-data proportion of other loci because locus missingness is calculated independently per column. The same filtering order is applied to direct dosage matrices, PLINK, genlight, genind, VCF, and sequence-alignment inputs. A lenient first pass removes the most incomplete loci before individual-level filtering, so that subsequent assessment of individual missingness is not driven by a relatively small number of incomplete loci. The removal of loci and individuals with high proportions of missing data follows recommendations for population-genomic data processing (Andrews et al., 2016; O'Leary et al., 2018). The final stricter pass is applied after incomplete individuals have been removed, so that locus-level missingness is recalculated among the retained individuals. The default thresholds provide a permissive starting point intended to remove poorly genotyped loci and heavily incomplete specimens while retaining

sufficient samples and loci for SOM analysis. These thresholds should be adjusted according to the characteristics of the dataset and the trade-off among sample retention, locus retention, and missingness (Andrews et al., 2016; Nazareno & Knowles, 2021). When the number of loci is limited, a more permissive locus-missingness threshold may be preferable, whereas datasets with very large numbers of loci can often tolerate stricter filtering. Missing values retained after filtering are handled through NA-aware SOM training rather than global imputation. Partial-data SOM training allows observations with missing variables to contribute information without being discarded. Best-matching units are identified using the observed dimensions, so missing variables do not contribute to unit selection. During training, codebook updates are likewise restricted to observed dimensions, while codebook values corresponding to missing variables are not updated. This partial-data treatment applies both during SOM training and when observations are mapped to the trained map, thereby avoiding artificial intermediate genotypes that can be introduced by simple mean imputation, which can reduce variation and distort the learned topology (Cottrell & Letrémy, 2005; Kohonen, 2001; Samad & Harp, 1992; Wehrens & Buydens, 2007). Although there are more specialized SOM-based imputation methods for continuous data (Folguera et al., 2015; La Torre et al., 2024; Rejeb et al., 2022), we use filtering combined with NA-aware training because *delimSOM* accommodates heterogeneous data types, including discrete and binary variables.

5. **Singleton and invariant locus filtering.** Optional singleton and invariant locus filters are performed by default (*singleton.loci.filter = TRUE*; *invariant.loci.filter = TRUE*). Singleton loci are removed after the lenient locus-missingness filter and individual missingness filter, because singleton status depends on the set of individuals retained after missingness filtering. In dosage-matrix workflows, genlight, PLINK .raw, and VCF-derived matrices, singleton loci are identified from the minor allele count, calculated as the smaller of the alternate- and reference-allele counts among called chromosomes. For sequence alignments analyzed with *alignment.ambiguity.mode = "heterozygote"*, singleton loci are identified using minor-allele counts on a diploid called-chromosome scale, so a heterozygote dosage of 1 contributes one alternate allele. When *alignment.ambiguity.mode = "missing"*, singleton loci are identified using allele counts among observed sequence states. In the genind workflow, singleton loci are identified by calculating the minimum observed allele count per locus from the allele-count matrix. Invariant loci are removed only at the final filtering step, after missing-data and singleton filters have been applied, because removing individuals can turn previously polymorphic loci into invariant loci. In dosage-matrix workflows, invariant loci are identified as columns where the finite minimum and maximum values are identical after excluding missing values. In alignment workflows, invariant loci are identified as encoded SNP columns whose finite minimum and maximum values are identical after filtering. In the genind workflow, invariant loci are identified by counting observed allele columns per locus from the allele-count matrix. Invariant loci do not contribute to delimitation because they carry no among-individual information and only inflate dimensionality. Singleton loci can also be disproportionately influenced by sequencing or calling artifacts, because very rare variants are especially difficult to distinguish from error in sequencing-based datasets (Johnston et al., 2015; Pfeifer, 2017). We therefore remove them by default to reduce sensitivity to potentially erroneous or weakly supported rare variants in the SOM input. Our *process.SNP.data.SOM* function returns a final SNP matrix with samples as rows and SNPs as columns.

6. **Preparation of binary matrices from categorical variables.** For non-genetic data layers that contain categorical variables, our *make.cols.binary.SOM* function can be used to convert selected factor or character columns in a data frame into binary 0/1 (presence/absence) indicator variables before SOM training. The categorical columns to be converted are specified with the *make.binary.cols* argument. The function extracts those selected columns and expands each level

into a separate binary indicator column. When *append.to.original = TRUE*, the *remove.original.cols* argument controls whether the original categorical columns are retained or removed after the indicator columns are appended. The *append.to.original* argument controls whether the output consists only of the binary indicator columns or of the original data frame with the binary columns appended. This workflow provides a standardized way to prepare categorical non-genetic variables for inclusion in single- or multilayer SOM analyses, such as phenotypic traits (e.g., color morph, pattern class, sex, or developmental stage), ecological variables (e.g., habitat type, host-plant category, substrate type, pollinator type, or microhabitat class), and geographic descriptors (e.g., drainage basin, island, or biogeographic region). We use this binary expansion because *train.SOM* expects numeric input. Converting categorical states into separate 0/1 indicators allows categorical variables to be incorporated without imposing an arbitrary numeric ranking among levels. Missing values are retained as NA. For example, consider a habitat-type variable. If a given column named "habitat" contains the values forest, grassland, wetland, and NA, the function generates separate binary columns such as habitat_forest, habitat_grassland, and habitat_wetland. A sample scored as forest would be coded 1, 0, 0; a sample scored as grassland would be coded 0, 1, 0; a sample scored as wetland would be coded 0, 0, 1; and a sample with missing habitat would remain NA, NA, NA across all derived columns.

7. **Removal of low-variation and correlated predictors.** For numeric predictor matrices, our *remove.lowCV.multicollinearity.SOM* function can be used to reduce noise and redundancy before SOM training. The input matrix or data frame is supplied with the *input.dataframe* argument. Additional arguments control the minimum coefficient of variation required for retention (*CV.threshold* argument), the maximum allowed absolute pairwise correlation among retained predictors (*cor.threshold* argument), the minimum prevalence required for rare binary states and count-variable nonzero values (*prevalence.threshold* argument), whether selected columns are excluded from filtering (*exclude.cols* argument), and whether filtering summaries are printed (*verbose* argument). The function first sets aside any columns listed in *exclude.cols* so that they bypass all filtering steps and are retained in the final output. Binary 0/1 variables are identified and filtered by minor-state prevalence rather than coefficient of variation, because their dispersion is constrained by binary coding and the coefficient of variation is not an appropriate measure of relative information content for this class of variables. Binary variables are removed when the rarer state occurs in fewer observations than the smallest whole number greater than or equal to the number of finite observations multiplied by the prevalence threshold. Non-negative integer-valued non-binary variables are identified as count variables, and count variables are removed when nonzero values occur in fewer observations than the smallest whole number greater than or equal to the number of finite observations multiplied by the prevalence threshold. This prevalence filter reduces the influence of very rare binary states and highly zero-inflated count variables on SOM distance calculations. For non-binary variables, the coefficients of variation are calculated after excluding non-finite values as the standard deviation divided by the absolute value of the mean after removal of missing values (Pearson, 1896; Sokal & Rohlf, 2012; Zar, 2010). Variables with coefficients of variation less than or equal to the specified *CV.threshold* are removed because variables with no or very low variability contribute little discriminatory information, while increasing dimensionality without meaningfully improving sample separation. Retaining such near-constant variables can worsen numerical conditioning in downstream analyses by inflating redundancy among predictors (Dormann et al., 2013; Greenacre & Primicerio, 2014). We recommend a default of *CV.threshold = 0.05*, corresponding to variables with less than 5% relative variability compared to their mean. We use the coefficient of variation rather than variance because it is unitless and scale-invariant, allowing variability to be compared directly across predictors measured in different units and magnitudes. When the mean is effectively zero, the function min–

max standardizes the variable and uses the standard deviation of the standardized values as a fallback variability measure, because the coefficient of variation becomes unstable when its denominator approaches zero. The function also calculates an absolute Spearman correlation (Spearman, 1904) matrix among the retained variables using pairwise complete observations and iteratively removes predictors from highly correlated pairs until no pair exceeds the specified *cor.threshold*. Strongly correlated predictors can overweight the same underlying gradient in distance-based learning and thereby inflate redundancy in the SOM input (Dormann et al., 2013; Gregorich et al., 2021). We use pairwise correlation rather than variance inflation factors because the latter are regression-oriented diagnostics defined relative to the full predictor set, whereas pairwise correlation provides a simpler rule for identifying near-duplicate variables (Dormann et al., 2013; Vatcheva & Lee, 2016). We recommend a default threshold of *cor.threshold = 0.9* so that highly correlated variables are removed while retaining moderately correlated variables. The function automatically selects which variable to remove from a highly correlated pair: it first removes the variable with more missing values, because the more complete variable preserves information for a larger number of samples. If missingness is equal, it removes the variable with lower variance, thereby retaining the variable with more variation across its observed range. If variability is effectively equal, it removes the variable with the larger mean absolute correlation to the remaining variables. Exact ties are resolved by removing the second variable in the selected pair. Any columns listed in *exclude.cols* are recombined with the retained filtered variables in their original column order, and the function returns a filtered data frame suitable for use in single- or multilayer SOM analyses.

8. **Preparation of single- or multilayer SOM inputs.** Our *train.SOM* function accepts a single matrix or data frame, or a list containing multiple matrices or data frames (*input_data* argument). These matrices may represent genetic, phenotypic, environmental, spatial, or other numeric variables, provided that rows represent individuals and row names consistently identify individuals across layers. Generally, we recommend supplying matrices that have been preprocessed using the functions described above, although unprocessed numeric datasets can also be used. The output of the preprocessing stage is a sample-by-variable numeric matrix or a list of such matrices. These processed data constitute the input to the *delimSOM* species-delimitation workflow implemented by our *train.SOM* function.

**M4: SOM training**

Our *train.SOM* function implements the core learning stage of *delimSOM* using the *kohonen v.3.0.13 R* package (Wehrens & Buydens, 2007; Wehrens & Kruisselbrink, 2018). It trains either a single- or a multilayer SOM, depending on whether the input consists of one or multiple matrices. The function performs input validation, sample and variable filtering, normalization, layer-type inference, automatic or user-defined grid construction, optional learning-rate tuning, and replicated SOM training with reproducible random seeds. This provides suitable default settings, reduces reliance on a single random initialization, and allows the stability of the learned structure to be assessed across replicate maps.

1. **Input validation and main training arguments.** The main SOM-training arguments are the number of training iterations (*N.steps* argument), the number of replicate SOM runs (*N.replicates* argument), whether to run replicates in parallel (*parallel* argument), the number of cores used when parallelizing (*N.cores* argument), the SOM grid dimensions (*grid.size* argument), the grid-size scaling parameter (*grid.multiplier* argument), the initial and final learning rates (*learning.rate.initial* and *learning.rate.final* arguments), whether to tune learning rates automatically (*learning.rate.tuning* argument), the neighborhood function (*training.neighborhoods* argument), and the maximum

allowed missing-data fractions per sample and variable (*max.NA.row* and *max.NA.col* arguments). The function also accepts optional user-specified layer distance functions (*layer.distance.functions* argument), optional user-defined layer weights (*manual.layer.weights* argument), saving options (*save.SOM.results*, *save.SOM.results.name*, and *overwrite.SOM.results* arguments), progress settings (*verbose* and *message.N.replicates* arguments), and a random seed (*set.seed.N* argument).

2. **Harmonization of rows and variables across layers.** If the input data is a list containing multiple matrices or data frames, the function first checks that all layers have row names and identifies the shared set of individuals across layers. Samples not shared across all matrices are removed. Each layer is then filtered to remove individuals exceeding the missing-data fraction specified by *max.NA.row*, and the shared-row intersection is performed again so that all retained layers contain the same final set of individuals. Within each layer, the function then removes nonnumeric columns, all-missing columns, variables exceeding the missing-data fraction specified by *max.NA.col*, and zero-variance variables. Rows that contain only missing values after column filtering are also removed, after which the shared-row intersection is performed once more. These repeated intersections ensure that every retained layer contains the same samples in the same order for joint SOM training.

3. **Removal of low-information variables and normalization.** After row matching and missing-data filtering, zero-variance variables and any all-missing rows or columns are removed. Each retained numeric variable is then columnwise min–max normalized to a 0–1 range. Variables are normalized because SOM training is distance-based, and variables measured on larger numeric scales could otherwise dominate the distance calculations and learned map structure, even when they are not biologically more informative. Columnwise min–max normalization places each variable on the same numerical range while preserving its ordering and relative spacing among individuals, which is important because SOM training, the learned topology, and quantization error are sensitive to variable scaling (Kohonen, 2013).

4. **Inference of layer type and assignment of distance functions.** Because SOM training is fundamentally distance-based, the choice of the distance function strongly influences how similarity among samples is represented on the map and is therefore an important part of model specification, especially in multi-layer analyses where different layers may require different notions of distance (Kohonen, 1998; Wehrens & Kruisselbrink, 2018). Because each layer is assigned its own distance function, we recommend that variables grouped within each layer have broadly similar data structures. For instance, a morphological dataset combining continuous measurements with binary presence/absence characters should be split into separate continuous and binary layers rather than analyzed as a single morphological layer with mixed structure. For each retained layer, our *train.SOM* function infers whether the matrix is continuous, binary (0/1 values), or 0/1/2 dosage-like and assigns a distance function accordingly; layer inference is done before normalization. Layers containing only 0/1 values are classified as binary and assigned normalized Hamming distance (*"tanimoto"*). Non-binary, integer-like layers whose observed values are restricted to 0, 1, and 2 are classified as count/dosage-like and assigned Manhattan distance. All other numeric layers are treated as continuous and assigned squared Euclidean distance (sumofsquares). Users can override automatic distance function inference via the *layer.distance.functions* argument. In this case, we recommend the following distance functions. For continuous quantitative variables, such as most climatic, morphometric, spatial, acoustic, physiological, or chemical data, we recommend squared Euclidean distance (*"sumofsquares"*), because these variables are naturally represented in continuous multivariate space and larger differences should generally contribute more strongly to sample separation. Binary presence/absence variables, such as habitat use, host association, pollinator association, parasite presence, symbiont presence, developmental mode, or gene presence, are often

better represented using normalized Hamming distance (*"tanimoto"*), because it measures the proportion of mismatching binary states between an individual and a codebook vector (Hamming, 1950; Wehrens & Kruisselbrink, 2018). SNP- and genotype-style encodings and other dosage-like genetic matrices (e.g., SNP dosages, copy number variants, structural variant counts, microsatellite repeat numbers, or short tandem repeat lengths) are better handled by Manhattan distance (*"manhattan"*), because it sums absolute per-locus dosage differences without squaring them. For raw count data, such as meristic morphological traits, abundance data, k-mer counts, transposable element insertion counts, or read-count summaries, Manhattan distance (*"manhattan"*) is often preferable when counts are sparse, skewed, or zero-heavy, because it sums absolute differences across variables and is less influenced by a few extreme values than squared Euclidean distance. Conversely, squared Euclidean distance (*"sumofsquares"*) may be preferable when large abundance differences are biologically meaningful and should contribute much more strongly to the separation of samples, or when counts have already been transformed to behave more like continuous predictors. For example, after log, square-root, or Hellinger transformation, count-derived variables often become more appropriately represented in Euclidean space than as raw discrete counts (Legendre & Gallagher, 2001). Allowing custom *layer.distance.functions* permits users to override these defaults for specialized datasets or in cases where the automatic structure detection is not appropriate. Nonetheless, the default behavior typically provides appropriate default distance functions for common layer types. If automatic detection is used (i.e., no *layer.distance.functions* are specified), users should verify that the inferred layer types and assigned distance functions are appropriate for their data (when *verbose = TRUE*, the inferred types are printed during *train.SOM* execution).

5. **Layer weighting.** To prevent domination by layers with larger average within-layer distance in multi-layer SOMs, internal distance normalization is applied (the default of the *supersom* function in *kohonen*). This helps downweight layers with thousands of variables (e.g., SNP data) or inherently high-variance structures and pairwise distances (e.g., SNP or binary data), while upweighting layers with inherently relatively less variance and lower pairwise distances (e.g., continuous and usually normally distributed morphological or climatic datasets). For each layer in each replicate, the internal normalization factor is computed as the reciprocal of the mean pairwise distance among the randomly selected initial codebook vectors and is stored in the *distance.weights* component of the trained model (Wehrens & Kruisselbrink, 2018). User-defined layer weights can be supplied through the *manual.layer.weights* argument. These are combined multiplicatively with the internal normalization weights to determine each layer's contribution to the combined distance used during SOM training, after which the resulting layer weights are rescaled to sum to 1. Hence, user-defined layer weights operate in combination with the internal distance-based normalization. User-defined layer weights can be used to emphasize or down-weight data types according to analytical goals, prior biological knowledge or hypotheses, or measurement uncertainty. For example, layers with higher measurement uncertainty, lower information content, or stronger environmental noise may be down-weighted, whereas layers with well-characterized evolutionary signal may be emphasized. Furthermore, users may adjust weights according to the effective independence of the information represented by different layers, because internal distance normalization adjusts differences in distance scale but does not account for linkage or redundancy within or among layers. This is particularly relevant for genetic data, where the number of variables (i.e., variable sites) does not necessarily correspond to the number of independent evolutionary histories. For example, a mitochondrial-marker layer such as COI may contain many variable sites but generally represents a single mitochondrial genealogy, whereas genome-wide SNPs distributed across many unlinked regions may sample many approximately independent genomic regions. Multiple genes from the same mitochondrial or plastid genome similarly provide additional sequence information but generally share the same underlying organellar history and should not be interpreted as independent genealogical replicates. For

multilocus nuclear sequence data, loci sampled from different genomic regions may provide approximately independent evolutionary information, but sites within each locus can remain linked. When such loci are concatenated, *delimSOM* treats retained variable sites as separate variables, so loci with more retained variable sites may contribute more strongly to within-layer distances even though those sites do not constitute equivalent independent loci. Similar considerations apply to SNP datasets: multiple SNPs within the same RAD locus, linkage-disequilibrium block, inversion, or other tightly linked genomic region may provide highly redundant information, whereas one SNP per widely distributed locus or appropriately LD-pruned SNPs more closely approximate independent genomic sampling. Microsatellite datasets should likewise be evaluated according to the number and linkage of loci rather than simply the number of alleles or variables: unlinked microsatellite loci may provide several approximately independent nuclear markers, whereas physically linked loci provide less independent information. Consequently, preprocessing to reduce within-layer redundancy, such as LD pruning or retaining representative markers from strongly linked regions, is generally preferable when appropriate, because down-weighting an entire layer reduces its overall influence but does not remove redundancy among variables within that layer. Redundancy can also occur among separate data layers. For example, spatial and environmental layers may both reflect the same geographic gradient, climatic and habitat layers may describe overlapping ecological variation, or a layer derived from another dataset may duplicate information already represented in the original layer. Such layers may be down-weighted when their joint inclusion would otherwise repeatedly represent the same underlying signal. Because information content, independence, and biological relevance are distinct properties, there is no universally appropriate weighting scheme; manual weights should therefore be biologically justified and their effects evaluated when alternative reasonable weighting or layer definitions could change the inferred delimitation. Replicate-specific internal distance-normalization weights are stored in *distance_weights_matrix*, describing the distance-scale correction applied to each layer (they are not layer importance measures and are not the final combined manual and internal layer weights).

6. **SOM grid type, size, and shape.** We implement a hexagonal SOM grid using the *somgrid* function in *kohonen*. This grid shape provides more even neighborhood relationships and clearer visualization than rectangular grids (Kohonen, 1998, 2013; Vesanto & Alhoniemi, 2000). The grid size is estimated automatically by default, but the user can specify it manually with the *grid.size* argument. For automatic estimation, the total number of grid units is scaled to the sample size, and the grid aspect ratio is determined from the major structure of the data. The final grid dimensions are obtained by combining the total number of units with the aspect ratio, so that the number of units along each axis is proportional to the square root of the total grid size. In other words, once the total number of grid units has been estimated from the sample size, the x and y dimensions are chosen so that their product approximates that total while their ratio reflects the dominant elongation of the data. The total number of grid units is calculated by multiplying the *grid.multiplier* by the square root of the number of samples. The shorter grid dimension is approximated from the square root of the total number of units divided by the inferred aspect ratio, and the longer dimension by dividing the total number of units by the shorter dimension, after which both are rounded to whole numbers. Thus, with the default *grid.multiplier = 5*, a dataset with 81 samples yields 5 × sqrt(81) = 45 grid units. If the inferred aspect ratio is approximately 3:1, the automatic sizing step gives ydim ≈ sqrt(45 / 3) ≈ 4, and xdim ≈ 45 / 4 ≈ 11, resulting in a grid of 11 × 4 units (rather than an approximately square-like 6 × 7). For single-layer inputs, the aspect ratio is estimated from the covariance structure of the retained numeric matrix using eigenvalue decomposition, first using complete cases and, if necessary, using mean-imputed values for columns with limited missingness. For multi-layer inputs, it is based on multiple factor analysis performed using the *MFA* function in the *FactoMineR v.2.13* R package (Lê et al., 2008), because a single covariance matrix is not an appropriate summary of datasets composed

of multiple layers with different variable sets, scales, and dimensionalities. Multiple factor analysis is specifically designed to summarize the dominant joint structure of such grouped datasets while balancing the influence of the different layers in the ordination. If multiple factor analysis cannot be calculated because too few usable rows or columns remain, the function falls back to a square-like aspect ratio. We use this automatic grid sizing to derive a data-informed grid size and shape from sample size and the dominant structure of the data. This is important because the number and arrangement of grid units strongly affect how finely the data manifold can be represented; a grid that is too coarse merges distinct structures, whereas an overly fine grid can fragment clusters, increase empty units, and slow computation (Brentan et al., 2018; Kohonen, 1995, 1998; Vesanto, 1999). We set a default *grid.multiplier = 5*, representing a heuristic compromise between under- and overfitting (Vesanto, 1999). Estimating aspect ratio from the dominant data axes is preferable to arbitrarily choosing a square grid, because elongated data manifolds are better represented by elongated maps, which can improve topology preservation and neighborhood continuity (Kohonen, 1995, 2001, 2014; Vesanto & Alhoniemi, 2000).

7. **Initialization of codebook vectors.** SOM codebook vectors are initialized by randomly selecting retained input samples without replacement. We use stochastic initialization rather than principal-component-based linear initialization because multi-layer inputs may contain different data types and layer-specific distance functions, whereas principal-component-based initialization requires a common covariance-based representation that is not readily defined for multilayer inputs using different distance functions (Kohonen, 1995, 1998, 2014). Initialization sensitivity is addressed by training many replicate maps (see next section) and summarizing recurring structure across replicates rather than relying on any single starting configuration. Although principal-component-based initialization can accelerate convergence in classical Euclidean SOM settings, random initialization provides a more suitable option for flexible multi-layer SOM workflows (Kohonen, 1998, 2013; Wehrens & Buydens, 2007).

8. **Number of steps and replicates.** Depending on dataset size and the number of layers, we recommend approximately 80–200 SOM training iterations (*N.steps*; default: *100*) and 80–150 replicate maps (*N.replicates*; default: *110*), reflecting a balance between convergence and computational tractability. Too few training steps can yield unstable codebook vectors, whereas too many steps add run time while often providing diminishing improvement in the learned topology (Kohonen, 2013; Sun, 2000). Likewise, too few replicates make the final delimitation more sensitive to random initialization, whereas very large numbers of replicates mainly increase computation time without proportionate improvement in stability (Kohonen, 2013).

9. **Learning-rate tuning.** SOM learning uses an "online" updating scheme through the *supersom* function in *kohonen*. Although batch SOM algorithms generally converge faster and avoid explicit learning-rate specification, we use the "online" algorithm because it allows direct control over learning-rate schedules, neighborhood contraction, and replicate-wise tuning (Kohonen, 1998, 2013). If *learning.rate.tuning = FALSE*, training proceeds with the specified initial and final learning rates (*learning.rate.initial = 0.5*, *learning.rate.final = 0.1* by default). If *learning.rate.tuning = TRUE*, the function evaluates a grid of candidate initial/final learning-rate combinations and selects the combination with the lowest mean quantization error across tuning replicates. This tuning mode is computationally expensive and is therefore only recommended for small datasets or short exploratory runs. Tuning evaluates a fixed grid of five initial learning rates (0.1, 0.3, 0.5, 0.7, 0.9) and five final learning rates (0.001, 0.01, 0.1, 0.3, 0.5), retaining only combinations in which the final learning rate is smaller than the initial learning rate. Candidate combinations are evaluated with 50-step runs and 10 replicate maps per combination, and the combination minimizing mean quantization error is retained.

10. **Neighborhood scheduling.** The neighborhood function can be chosen via the *training.neighborhoods* argument and defaults to a Gaussian kernel (*"gaussian"*), with *"bubble"* available as an alternative. The neighborhood radius decreases linearly from the 2/3 quantile of pairwise grid-unit distances to zero across training. This shifts learning from global ordering in early iterations to local refinement later, facilitating stable convergence and fine-scale adaptation of codebook vectors (Kohonen, 2014; Wehrens & Kruisselbrink, 2018). Although the *kohonen* package uses the bubble neighborhood as a faster default, we set the Gaussian neighborhood as default because Gaussian neighborhoods provide smoother distance-decay across the map and often yield lower quantization error (Wehrens & Kruisselbrink 2018; Stefanovič & Kurasova 2011; Natita et al. 2016).

11. **Replicated SOM training and parallelization.** Our *train.SOM* function trains multiple SOMs (*N.replicates* argument) and stores, for each replicate, the trained SOM model, the learning trajectory for each layer, the layer-distance normalization weights, the quantization error, and the topographic error. This is consistent with standard SOM practice, because random initialization and stochastic training order can lead to different local organizations even under identical parameter settings (Wehrens & Buydens 2007; Wehrens & Kruisselbrink 2018). Replicates can be run in parallel using PSOCK clusters via the *makeCluster*, *clusterExport*, and *stopCluster* functions in the *parallel v.4.5.1 R base* package (R Core Team, 2025), the *registerDoSNOW* function in the *doSNOW v.1.0.20 R* package (Daniel et al., 2022), the *foreach* function in the *foreach v.1.5.2 R* package for parallel execution (Microsoft, 2022), and the *registerDoRNG* function in the *doRNG v.1.8.6.3 R* package for reproducible random-number generation (Gaujoux, 2026). Parallelization facilitates replicate-based SOM inference, which would otherwise be slow for larger genomic or multilayer datasets. Relative to the original SOM implementation (Pyron, 2023; Pyron et al., 2023), *delimSOM* 2.0 improves computational scalability for several reasons. First, replicate runs can be parallelized across cores rather than executed serially, which is especially useful for large datasets. Second, the earlier implementation relies on repeated explicit for loops within separate hard-coded one-, two-, three-, and four-layer workflows, whereas the current implementation replaces those repeated serial loop structures with a single training framework that uses *lapply* for non-parallel replicate execution and *foreach* for parallel replicate execution. Third, preprocessing, layer handling, error tracking, and result assembly are handled more efficiently within this unified workflow, reducing repeated bookkeeping across replicates and layer configurations. We compared runtimes using simulated datasets containing 50 samples and 10, 100, 1,000, 10,000, or 100,000 variables. All analyses used a 4 × 3 SOM grid, 150 training steps, and 110 replicate runs, with four processor cores used for parallel execution. Parallel execution improved the computational scalability of *delimSOM* 2.0 for large datasets, whereas runtimes were less efficient for small datasets and for non-parallel execution compared with the original implementation (Supplementary Table S15).

12. **Quantization and topographic error.** For each replicate, quantization error is computed from the trained SOM as the mean sample-to-best-matching-unit distance. Topographic error is computed for each replicate by taking each sample's first-best-matching unit from the trained SOM and calculating its second-best-matching unit from the combined sample-to-codebook distances across all layers, using the layer-specific distance functions and the normalized products of the user-defined and internal distance-normalization weights. Topographic error is then calculated as the fraction of samples with valid first- and second-best-matching units for which those units are not adjacent, where adjacency is defined by a grid-unit distance of one. Quantization error measures local representation accuracy (how closely the codebook vectors fit the data points) (Sun, 2000). Topographic error measures whether neighborhood relationships in the data are preserved on the map (Sun, 2000). We retain both because they measure complementary aspects of SOM quality and local representation: accuracy and topological continuity; a map can achieve relatively low quantization error while still

introducing substantial topological distortion, or conversely preserve neighborhood continuity while fitting individual observations less accurately (Kiviluoto, 1996; S.-H. Park et al., 2014; Sun, 2000).

**M5: Clustering of SOM codebook vectors and inference of K**

Our *clustering.SOM* function infers species-delimitation hypotheses by clustering the SOM codebook vectors. Compared to the prior implementation (Pyron, 2023; Pyron et al., 2023), *delimSOM* 2.0 separates SOM training from clustering, allowing users to apply different clustering analyses to the same trained SOM replicates without rerunning SOM training each time. It also allows poorly fitting SOM replicates to be removed before clustering.

1. **Input and main clustering arguments.** Our *clustering.SOM* function requires a *SOM.output* object returned by *train.SOM*. The main arguments are the maximum tested number of clusters (*max.k* argument), an optional fixed number of clusters (*set.k* argument), the clustering method (*clustering.method* argument), the BIC (Bayesian Information Criterion) threshold used in threshold-based K selection (*BIC.thresh* argument), optional quantile cutoffs for quantization-error filtering (*quantization.error.quantile* argument) and topographic-error filtering (*topographic.error.quantile* argument), whether to calculate replicate-specific soft ancestry matrices (*calculate.soft.ancestry* argument), whether to calculate variable-importance summaries (*calculate.variable.importance* argument), whether to print progress messages (*verbose* argument), the frequency of replicate-level progress messages (*message.N.replicates* argument), whether to save clustered SOM results to file (*save.SOM.results* argument), the clustered-results file name (*save.SOM.results.name* argument), whether to overwrite existing saved clustering results (*overwrite.SOM.results* argument), and the random seed (*set.seed.N* argument).

2. **Filtering of poorly fitting replicates.** Before any clustering is performed, the function can remove replicate SOM runs with unusually high quantization error and/or topographic error (Kohonen, 1995). This removes maps that fit the data poorly or do not preserve the relationships among samples well. For each requested diagnostic, a cutoff is defined from the specified quantile, and any replicate with error values above that cutoff is removed (*quantization.error.quantile* and *topographic.error.quantile* arguments; default: 95%). Filtering is applied consistently across all stored replicate outputs, including the SOM models, the learning-value matrices, and the distance-weight matrix, so that all downstream summaries remain aligned to the same retained replicate set. We recommend this filtering because results should not be driven by a subset of poorly fitted or topologically distorted maps. The 95th percentile is conservative: it removes only the worst-fitting maps while retaining most replicates for stable summarization.

3. **Extraction of codebook vectors.** For each retained SOM replicate, our *clustering.SOM* function extracts the codebook vectors using the *getCodes* function in *kohonen*. If multiple layers are present, the layer-specific codebooks are concatenated columnwise into a combined neuron-by-variable matrix. Codebook vectors are then clustered rather than individual samples. This follows the standard two-level SOM clustering strategy, in which the SOM first regularizes the data into local codebook vectors, and clustering is then applied at the second abstraction level, reducing computational burden and often sensitivity to individual-level noise because codebook vectors are local "average" representations of the original observations (Vesanto & Alhoniemi, 2000). Unlike common dimensionality-reducing statistical tools like principal component analysis, this does not primarily reduce the number of variables by projecting samples onto a small number of orthogonal axes. Instead, it regularizes the data by representing the individual observations by a smaller set of topology-preserving codebook vectors, thereby reducing sample-level noise and simplifying the set

of objects subjected to clustering, even though the codebook vectors remain in the original variable space (i.e., the codebook vectors have the same number of variables as the original data).

4. **SOM codebook clustering and selection of K.** Our *clustering.SOM* function supports six clustering methods specified via the *clustering.method* argument: *"kmeans+BICelbow"*, *"kmeans+BICthreshold"*, *"GMM+BICthreshold"*, *"hierarchical+DB"*, *"HDBSCAN"*, and *"OPTICS+Silhouette"*. For each retained SOM replicate, the SOM codebook vectors are extracted, concatenated across layers if necessary, and clustered. Generally, no single clustering algorithm performs best across all datasets, because clustering performance depends strongly on data structure, cluster geometry, dimensionality, noise, overlap, distance metric, and parameterization (de Souto et al., 2008; Omran et al., 2007; Rodriguez et al., 2019). For instance, methods that perform well for compact, approximately spherical groups may perform poorly for irregular, elongated, hierarchical, or variable-density structures (Omran et al., 2007; Vesanto & Alhoniemi, 2000). Therefore, our function provides different centroid-based, model-based, hierarchical, and density-based approaches on codebook vectors to accommodate alternative cluster geometries and delimitation patterns rather than assuming that a single clustering rule will perform best across all datasets. Classical SOM clustering studies have commonly used hierarchical clustering and U-matrix-guided interpretation, especially when structure is irregular, nonspherical, or hierarchically nested (Flexer, 2001; S.-H. Park et al., 2014; Y.-S. Park et al., 2006; Vesanto & Alhoniemi, 2000). Other SOM studies have likewise combined SOM training with secondary clustering of codebook vectors or map units using hierarchical or partitional methods, including Ward-type hierarchical clustering, k-means, and Davies–Bouldin-guided cluster selection (Licen et al., 2020; Raivio et al., 2001; Zhengtian et al., 2023). Likewise, different internal validity indices and model-selection criteria emphasize different aspects of cluster structure, and their performance depends strongly on data characteristics such as dimensionality, overlap, outliers, noisy variables, and the clustering algorithm used (Fraley & Raftery, 1998, 2002; Guerra et al., 2012; Mehar et al., 2013; Milligan & Cooper, 1985; Rodriguez et al., 2019). Consequently, no single metric consistently identifies the optimal K across all scenarios, and determining the optimal number of clusters without prior knowledge remains an inherently difficult and often contentious problem in unsupervised clustering (Hennig, 2015; Kass & Raftery, 1995; Mirkin, 2011; Omran et al., 2007; Rodriguez et al., 2019; Tibshirani et al., 2001). We recommend using *"kmeans+BICelbow"* or *"kmeans+BICthreshold"* for our SOM approach here, as these clustering methods consistently performed best for species delimitation in our testing (this study) as well as the prior SOM species delimitation implementation (Pyron, 2023; Pyron et al., 2023). Users may also apply the other clustering methods and treat the resulting partitions as complementary delimitation hypotheses rather than assuming one universally optimal method. Although partitioning around medoids has recently been used in a machine-learning species-delimitation workflow (Derkarabetian et al., 2019), we did not implement and test this method as a clustering option here because it was consistently much slower and showed poorer performance than the retained methods in trial runs, making it impractical for large sets of codebook vectors.

   a. ***"kmeans+BICelbow"* (recommended):** This method clusters SOM codebook vectors across candidate values of K with k-means, converts the resulting within-cluster dispersion profile into a BIC-like support curve, and then selects K using an elbow-based decision rule. Unlike the DB-guided k-means approaches used in some earlier SOM studies (Licen et al., 2020; Raivio et al., 2001; Vesanto & Alhoniemi, 2000), we use a BIC-elbow rule because it evaluates the one-cluster case and tests the initial improvement from K1 to K2 against a support threshold before considering further subdivision. A BIC-like criterion is used because it balances decreasing within-cluster dispersion against an explicit penalty for increasing K (Kass & Raftery, 1995). First, SOM codebook vectors are clustered with k-means (Lloyd, 1982; MacQueen, 1967a) across candidate values of K, and the within-cluster sum of squares is calculated for each candidate partition. K-means is one of the most widely used clustering algorithms and is computationally efficient because it partitions observations by minimizing within-cluster variation under Euclidean distance (Omran et al., 2007). K-means depends on the initial starting configuration and is best suited to compact, approximately spherical clusters of similar size under Euclidean distance, so it can perform poorly when clusters are non-convex, elongated, overlapping, or strongly unequal in size (E. Davies, 1997; Jain, 2010; MacQueen, 1967b; Steinley, 2006). Nevertheless, it has been applied successfully in high-dimensional datasets (Jain, 2010; Steinley, 2006; Wu et al., 2008), including gene-expression studies (Costa et al., 2004; de Souto et al., 2008) and population genetic workflows (Jombart et al., 2010). K-means is run using the *kmeans* function (*nstart = 30*, *iter.max = 1e5*) in the *base R stats v.4.5.1* package (R Core Team, 2025). The within-cluster sum of squares is converted into a BIC-like criterion by multiplying the number of SOM codebook vectors by the natural logarithm of the mean within-cluster sum of squares and adding a complexity penalty equal to the number of clusters multiplied by the natural logarithm of the number of codebook vectors. The K = 1 value is calculated as the total sum of squares around the global mean. For each candidate K ≥ 2, k-means is fitted, and the total within-cluster sum of squares is retained. Lower BIC values are treated as better support. The initial improvement from K = 1 to K = 2 is first evaluated. If this drop is smaller than the specified *BIC.thresh* value, this indicates that a two-cluster solution (K= 2) is not sufficiently supported and K = 1 is selected. Otherwise, the successive BIC improvements (i.e., the change in BIC from K minus 1 to K) are grouped into two classes via Ward clustering using Euclidean distances computed with the *dist* function, agglomerative hierarchical clustering with the *hclust* function (*method = "ward.D2"*), and final group assignment with the *cutree* function in *base stats*. The class with the larger mean BIC drop is interpreted as containing the larger improvements. Ward clustering is used here because it groups values by minimizing increases in within-group sum of squares (Ward, 1963), making it suitable for separating large BIC drops from smaller residual improvements. The candidate K values corresponding to transitions in this class are identified, and the candidate with the lowest BIC-like value is selected. This follows an elbow-style model-selection logic, in which larger BIC improvements are distinguished from smaller subsequent changes (Thorndike, 1953). This elbow rule conceptually follows the *diffNgroup* criterion used in discriminant analysis of principal components (Jombart et al., 2010), in that both methods group successive changes in BIC using Ward clustering to distinguish major early improvement from later minor gains. However, our SOM implementation additionally evaluates whether the initial improvement from K = 1 to K = 2 exceeds *BIC.thresh*, and then selects the lowest-BIC candidate associated with the class having the larger mean BIC drop rather than simply choosing the value immediately before the first group switch. The initial improvement step is evaluated first because, if a two-cluster solution is not clearly better than a single cluster, there is little evidence for meaningful partition structure in that replicate. The final K is then taken as the lowest-BIC solution within the main improvement region rather than the value immediately before the first group switch, because the grouped elbow structure can still contain large jumps associated with the upward phase of the BIC curve after the main improvement has already been achieved. Hence, selecting the lowest-BIC solution within that region avoids overestimating K simply because one boundary point happens to occur near a sharp rebound in BIC. If no clear elbow group can be identified, the method falls back to the global minimum-BIC solution. When *set.k* is supplied, the function does not evaluate all candidate values of K but fits and returns the requested K. We recommend a default of *BIC.thresh = 6* following general BIC interpretations, under which values from 6 to 10 indicate strong support, values greater than 10 indicate very strong support, and values below 6 indicate only weaker support for the more complex solution (Kass & Raftery, 1995).

b. ***"kmeans+BICthreshold"***: This method is similar to ***"kmeans+BICelbow"*** (Supplementary Methods M5.4a), but it selects K using an explicit BIC-threshold rule rather than an elbow-based detector. Although such a BIC-threshold rule has not been used for SOM clustering studies yet, there are some empirical examples in broader clustering workflows (Hamerly et al., 2005; Valente & Wellekens, 2004). The function identifies the first value of K at which the improvement relative to the previous K falls below the specified *BIC.thresh* value, and then selects the previous K. If the first improvement from K = 1 to K = 2 is already below the threshold, K = 1 is selected. This provides an explicit and conservative decision rule for whether additional clusters are sufficiently supported, because once the incremental BIC improvement drops below the chosen support threshold, further subdivision is no longer treated as meaningfully justified. If none of the evaluated improvements falls below *BIC.thresh*, the K with the lowest BIC value is selected.

c. ***"GMM+BICthreshold"***: This method fits Gaussian mixture models (Fraley & Raftery, 1998) to the codebook vectors and selects K with the same threshold-based rule used for *"kmeans+BICthreshold"* (Supplementary Methods M5.4b). SOM studies have not used this specific combination for SOM codebook-vector clustering, but Gaussian finite mixture models with BIC-based model selection are well established in model-based clustering more broadly (Fraley & Raftery 2002, 2003; Scrucca et al. 2016). Furthermore, a similar clustering approach has recently been used in a machine-learning species delimitation study (Derkarabetian et al., 2019). The function fits Gaussian mixture models using the *mclustBIC* function in the *mclust v. 6.1.2 R* package (Scrucca et al., 2023). For each K, it evaluates candidate covariance parameterizations in a tiered manner (7 model groups), first testing a stable subset of models and then broader subsets if needed, retaining the best finite BIC value for each K. When *set.k* is supplied, only that fixed value of K is evaluated. This staged evaluation is used to reduce unnecessary fitting of more complex covariance structures when simpler models already provide stable solutions. Because the Gaussian mixture implementation treats larger BIC values as better, the sign of the BIC vector is inverted before applying the threshold rule (following our other *BICthreshold* methods). This overall model-based clustering approach is useful when clusters are better approximated by ellipsoidal distributions with unequal covariance structure than by centroid-based partitions, because Gaussian mixture models can represent differences in cluster shape, volume, and orientation that k-means cannot, but it relies on Gaussian mixture assumptions and requires sufficient codebook vectors to estimate the selected covariance structure (Fraley & Raftery, 2007, 1998; Scrucca et al., 2016).

d. ***"hierarchical+DB"***: This method performs agglomerative hierarchical clustering of the codebook vectors and selects K using the Davies–Bouldin (DB) criterion (D. L. Davies & Bouldin, 1979). Hierarchical clustering is commonly used as a downstream clustering approach in SOM studies (S.-H. Park et al., 2014; Y.-S. Park et al., 2006; Raivio et al., 2001; Vesanto & Alhoniemi, 2000; Zhengtian et al., 2023). Several SOM studies have also evaluated hierarchical or partitional clustering solutions with the DB index, supporting its use as a heuristic criterion for selecting compact and well-separated partitions of SOM-derived representations (Licen et al., 2020; Raivio et al., 2001; Vesanto & Alhoniemi, 2000). First, pairwise Euclidean distances among SOM codebook vectors are computed with the *dist* function in *base stats*, and agglomerative hierarchical clustering is then performed with the *hclust* function (*method = "ward.D2"*) using Ward's D2 linkage method (Murtagh & Legendre, 2014). Final cluster assignments for a given K are obtained with the *cutree* function in *stats*, which cuts the hierarchical dendrogram at the specified number of groups and assigns each codebook vector to one of those resulting clusters. Ward's D2 method is used because, unlike linkage methods such as single, complete, or average linkage, it merges clusters to minimize the increase in total within-cluster variance at each step, which tends to favor compact and relatively well-separated groups (Murtagh & Legendre, 2014). This makes it especially suitable when the goal is to evaluate partitions with the DB index, which likewise favors low within-cluster dispersion and strong between-cluster separation (D. L. Davies & Bouldin, 1979). The DB is calculated with the *intCriteria* function (*crit = "Davies_Bouldin"*) in the *clusterCrit v.1.3 R* package (Desgraupes, 2023). Lower DB values indicate more compact and better-separated clusters. If *set.k* is supplied, the dendrogram is simply cut at that user-specified k. If *set.k* is not supplied, the function first evaluates whether a two-cluster partition is supported beyond random expectation, because a replicate that is not clearly better represented by two clusters than by randomized structure should not be forced into a multi-cluster solution. This is done by comparing the observed DB value for K = 2 with a null distribution generated from 500 column-wise permutations of the codebook matrix. For each permutation, values are independently shuffled within columns, Euclidean distances are recalculated, hierarchical clustering is repeated, the permuted dendrogram is cut at K = 2, and the DB index is recalculated. The permutation p-value is calculated as the proportion of the remaining null values that are less than or equal to the observed DB value. If the observed K = 2 value is not significantly smaller than the null expectation ($p \geq 0.05$), K = 1 is selected. If the permutation p-value is below 0.05, DB values are calculated for each candidate K from 2 through *max.k* by cutting the same dendrogram into K clusters. K is then selected as the candidate with the lowest finite DB value. The final K is then chosen as the partition with the minimum DB value. If all DB values for $K \geq 2$ are invalid or non-finite, K = 1 is selected because no multi-cluster solution can be ranked reliably by the DB criterion. This method is useful because hierarchical clustering does not require K to be fixed before the dendrogram is built and can better reflect nested or multilevel structure in the codebook vector space, but it can also be sensitive to early agglomeration decisions and to distortions introduced high in the dendrogram.

e. ***"HDBSCAN"*:** This method applies hierarchical density-based clustering (HDBSCAN) (Campello et al., 2013) to the codebook vectors and selects K using membership probability and silhouette support after noise handling. HDBSCAN is a density-based clustering method that builds a hierarchy of dense regions and identifies clusters as groups of points separated by density valleys, while allowing points in sparse regions to remain unassigned as noise. Prior SOM clustering studies have not used this clustering method yet, and it is sensitive to data structure (Rodriguez et al., 2019). Nonetheless, HDBSCAN is a well-established clustering method more broadly, including in biological clustering workflows, such as metagenomic binning and single-cell analyses (Campello et al., 2013, 2015; Duan & Hon, 2021; Hickl et al., 2022; McInnes et al., 2017). HDBSCAN is performed with the *hdbscan* function in the *dbscan v.1.2.4 R* package (Hahsler & Piekenbrock, 2025) using Euclidean distances among codebook vectors, and candidate parameterizations are evaluated across a range of *minPts* values. The *minPts* parameter specifies the minimum neighborhood size used to define local density, so smaller values make the method more sensitive to fine-scale dense structure, whereas larger values require broader and more stable density-supported groups. The maximum candidate *minPts* value is the larger of 2 and the smallest of 15, one less than the number of codebook vectors, and half the number of codebook vectors rounded down. The minimum candidate is the larger of 3 and 5% of the number of codebook vectors rounded down, capped at the maximum candidate. Up to 15 unique values are evaluated at equal intervals between these bounds. For each candidate *minPts* value, the method records the number of non-noise clusters and the mean membership probability of the non-noise points. This quantity summarizes how confidently retained codebook vectors belong to their assigned density clusters, so higher values indicate more stable density-supported assignments. If *set.k* is supplied, only parameterizations yielding exactly that number of clusters are considered; if no tested *minPts* value produces the requested number of clusters, the replicate stops. When *set.k* is not supplied, valid *minPts* parameterizations are first ranked by the mean HDBSCAN membership probability of their non-noise points, and the parameterization with the highest mean membership probability is selected for post-processing. For each candidate *minPts* value, the initial candidate K is defined as the number of non-noise clusters before noise reassignment. After noise handling, the final replicate-specific K is defined as the number of clusters in the selected post-processed solution. Because downstream ancestry estimation requires every SOM unit to have a cluster label, noise points are then handled explicitly. When *set.k* is specified, noise points are reassigned to the nearest core cluster using nearest-neighbor matching through the internal *get.knnx.custom* function. This custom function identifies the nearest non-noise codebook vector based on Euclidean distance and assigns the noise point to that vector's cluster, so that the requested number of clusters is preserved while avoiding unassigned SOM units. When *set.k* is not specified, two post-processing strategies are compared. In the first strategy, all noise points are reassigned to the nearest core cluster using the same nearest-neighbor procedure, which favors a more compact interpretation of the density structure. In the second strategy, the 95th percentile of the distances from noise vectors to their nearest non-noise vectors is calculated. Each noise vector with a distance above this threshold is assigned to a separate singleton cluster, whereas the remaining noise vectors are assigned to their nearest non-noise clusters. The better of these two post-processed solutions is then selected by silhouette support when possible. The silhouette criterion measures how well each point matches its assigned cluster relative to neighboring clusters, so higher average silhouette values indicate more cohesive and better-separated clusters after noise handling (Rousseeuw, 1987). If no valid parameterization is found and *set.k* is not supplied, K = 1 is selected because no density-based multi-cluster solution can then be supported reliably for that replicate. If *set.k* is supplied and greater than 1, failure to recover the requested number of clusters is treated as an error because HDBSCAN uses *set.k* as a filter over density-based solutions rather than as a forced partitioning step. This method is particularly useful when putative taxon boundaries are better represented by density valleys, variable-density structure, or irregular cluster shapes than by centroid separation, because HDBSCAN does not require clusters to be approximately spherical and does not require K to be specified before the density hierarchy is built. However, it is also sensitive to density parameterization and to how noise points are treated after clustering, and should therefore be interpreted alongside the other clustering methods.

f. "***OPTICS+Silhouette***": This method applies OPTICS (Ankerst et al., 1999) to the SOM codebook vectors and selects K by mean silhouette width (Rousseeuw, 1987). OPTICS is a density-based ordering procedure that does not directly return a single partition, but produces an ordering of points and their reachability structure, from which cluster partitions can subsequently be extracted under different parameter settings. Prior SOM clustering studies have not used this clustering method yet, and it is sensitive to data structure (Rodriguez et al., 2019). However, OPTICS has been applied by a few studies in broad clustering workflows (Rodriguez et al., 2019; Rodríguez-Echeverría et al., 2020), sometimes together with silhouette-based validation (Rodríguez-Echeverría et al., 2020). Silhouette remains common in both benchmarking and applied clustering studies (Batool & Hennig, 2021; Guerra et al., 2012). OPTICS is performed on the codebook vectors with the *optics* function in *dbscan*, and candidate partitions are then extracted with the *extractXi* function in *dbscan* across combinations of *minPts* and *xi*. The *minPts* parameter controls the minimum neighborhood size used to define local density. Smaller *minPts* values make the method more sensitive to fine-scale dense structure, whereas larger values require broader and more stable density-supported groups. The parameter *xi* controls the minimum relative steepness required to identify density drops and rises as cluster boundaries. Smaller *xi* values permit finer subdivision of the reachability structure, whereas larger values require more pronounced density changes before clusters are separated. When *set.k* is supplied, an adaptive *minPts* grid is generated by defining the maximum candidate value as the larger of 2 or the smaller of 15, half the number of codebook units, or one less than the number of codebook units. The starting value is defined as the smaller of this maximum and the larger of 3 or 10% of the codebook units, ensuring that the *minPts* range remains valid even for small SOM grids. At least one and at most eight evenly spaced *minPts* values are evaluated, together with *xi* values of 0.05, 0.10, 0.20, 0.30, and 0.40. When *set.k* is not specified, the same adaptive procedure is used, but the starting value is based on the larger of 5 or 10% of the codebook units before being capped by the maximum valid *minPts* value. For each candidate parameter combination, the *extractXi* function is used to derive a cluster assignment from the OPTICS ordering; solutions that fail, contain only noise, exceed *max.k*, or do not match the user-specified K are discarded. Because downstream ancestry estimation requires every SOM unit to have a cluster label, each codebook vector labelled as noise is matched to its nearest non-noise codebook vector using Euclidean distance and k = 1 via the custom *get.knnx.custom* function. The noise vector is then assigned to the cluster of that nearest non-noise vector, and all cluster labels are subsequently converted to consecutive integers. Candidate solutions are ranked by mean silhouette width (Rousseeuw, 1987), and the valid solution with the highest silhouette support is retained. If *set.k* is supplied, only solutions with exactly that number of clusters are considered; otherwise, all valid solutions with at least two clusters are evaluated. When *set.k* is supplied, failure to find an OPTICS *minPts*/*xi* combination yielding exactly the requested number of clusters is treated as an error. When *set.k* is not supplied, K = 1 is selected for that replicate if no valid parameter combination with at least two clusters is found or if the highest mean silhouette width is missing or less than or equal to 0.15, indicating that no multi-cluster OPTICS solution is sufficiently well separated. This method is useful when putative taxon boundaries are better represented by irregularly shaped or variable-density clusters than by compact centroid-based groups, because OPTICS does not impose a single global density threshold before constructing the density ordering. However, it remains sensitive to parameterization and to the treatment of noise points after clustering.

5. **Assignment of samples and ancestry coefficients.** After clustering the SOM codebook vectors, our *clustering.SOM* function derives hard sample assignments and replicate-specific soft cluster-membership probabilities for each retained SOM replicate. Hard sample-level assignments are obtained by mapping each individual to the cluster label of its assigned best-matching SOM unit. For each replicate, cluster_gridcell_assignments stores one row per sample containing its best-matching SOM-unit number and the corresponding replicate-specific cluster label. When *calculate.soft.ancestry = TRUE*, the function calculates sample-to-unit distances separately for each SOM layer using the corresponding distance function and combines them using the products of the user-defined and internal distance-normalization weights. The default temperature is the median of the finite integrated distances, with a value of 1 used if this median is non-positive or non-finite. Distances are converted to exponentially decreasing unit weights, normalized across units for each sample, summed across the units belonging to each cluster, and renormalized to obtain *replicate_ancestry_matrix*. After Hungarian label harmonization, these replicate-specific soft matrices are relabeled to the same cluster-label space as the hard assignments and are returned collectively as *replicate_ancestry_matrices*. Because cluster labels are arbitrary across replicate clustering runs, direct comparison of raw replicate labels is not meaningful unless labels are first aligned to a common reference. To achieve this, the function generates a reference labeling for each value of K from 1 to *max_reference_k*, where *max_reference_k* is the smallest of *max.k*, the largest

replicate-specific selected K, and the number of unique rows in the processed input data. These reference clusterings are stored (*cluster_labels*) and are used only for cross-replicate label harmonization. The function then forms *assignment_matrix* by retaining replicate assignments with selected K less than or equal to *max_reference_k*. For each retained replicate with K ≥ 2, the corresponding replicate-specific sample labels are compared with the reference labels for the same K by constructing a cluster-overlap count table in which rows correspond to reference clusters and columns correspond to the replicate's current cluster labels. The overlap matrix is converted to a cost matrix by subtracting overlap counts from the maximum overlap value, and the optimal one-to-one label correspondence is obtained with the *solve_LSAP* function in the *clue v.0.3.67 R* package (Hornik, 2005), which solves the linear sum assignment problem with the Hungarian method (Kuhn, 1955; Munkres, 1957). The resulting mapping is inverted so that each replicate-specific cluster label is reassigned to the matched reference label, and the relabeled assignments are written back into *assignment_matrix*. We use this relabeling step because replicate SOM clustering runs can recover the same partition with permuted numeric labels, making label harmonization necessary before replicate summaries can be interpreted (Hornik, 2005; Jakobsson & Rosenberg, 2007; Kopelman et al., 2015). Label harmonization is performed separately against a different reference partition for each value of K. Cluster labels are therefore harmonized among replicates sharing the same K, but a label such as Cluster 1 is not guaranteed to represent the same group at other K. Consequently, the replicate-consensus *ancestry_matrix* is most straightforwardly interpreted when the retained replicates share the same K. After relabeling, the function constructs *ancestry_matrix* by tabulating, for each individual, the frequency with which that individual is assigned to each relabeled cluster across retained SOM replicates and converting those counts to proportions with the *prop.table* function in *base R*. Thus, the final *ancestry_matrix* stored in the results object is a replicate-consensus summary derived from relabeled hard sample assignments rather than a direct average of the replicate-specific soft ancestry matrices. The resulting matrix has one row per individual, and one column for each cluster label from 1 through the largest K represented in *assignment_matrix*, with entries summing to 1 across rows. The replicate-consensus proportions in *ancestry_matrix* are somewhat equivalent to individual ancestry coefficients (Beugin et al., 2018; Frichot et al., 2014; Pritchard et al., 2000). These "species coefficients" indicate how consistently each individual is assigned to each of the K inferred clusters (Pyron, 2023; Pyron et al., 2023). We retain this output because species delimitation is not always strictly discrete, and replicate-consensus assignment profiles are useful for identifying strongly assigned, weakly assigned, or intermediate individuals, especially in cases of recent divergence, incomplete lineage sorting, gene flow, hybridization, or discordance among genetic, phenotypic, spatial, and environmental data layers. When replicate-specific soft ancestry matrices are calculated, the function summarizes them with two assignment-sharpness diagnostics: *mean_assignment_margin*, which measures the average difference between the highest and second-highest cluster probabilities across individuals, and *mean_normalized_assignment_entropy*, which measures the average diffuseness of assignment probabilities across clusters. The *ancestry_matrix* is the main downstream delimitation output used in our Structure-like barplots and geographic assignment maps. Accordingly, *clustering.SOM* returns two complementary assignment summaries: replicate-specific soft assignment matrices (*replicate_ancestry_matrices*) that capture within-replicate assignment uncertainty, and a final hard-assignment consensus ancestry matrix (*ancestry_matrix*) that summarizes how consistently each individual is assigned to each cluster across retained replicates.

6. **Variable-importance summaries.** Our *clustering.SOM* function computes two complementary variable-importance summaries for each SOM layer (when *calculate.variable.importance = TRUE*): *median_etasquared_variable_importance* and *median_map_variance_variable_importance*. These are calculated after the final replicate-specific SOM-unit clusters and sample assignments have been

obtained, and summarize how strongly each variable is associated with separation among the inferred clusters or varies across the trained SOM map. The function first extracts the layer-specific codebook matrices from each retained SOM replicate using the *getCodes* function in *kohonen* and coerces single-layer outputs to list format if needed. For each layer, it then iterates across retained replicates, extracts the corresponding codebook matrix, and calculates both a map-variance importance vector and a cluster-separation importance vector for all variables in that layer. These replicate-specific vectors are combined across replicates and summarized by their median values to produce the final layer-specific importance summaries. The first variable-importance measure, map variance, quantifies how strongly the codebook values of each variable vary among neurons. It is calculated by first counting how many samples map to each neuron and then defining a neuron's weight as the number of mapped samples plus a baseline value of 1. This weighting gives greater influence to neurons that represent more samples and lower influence to empty or sparsely occupied neurons, thereby emphasizing regions of the map that are more strongly supported by the observed data. Adding the baseline value ensures that empty neurons still contribute to the calculation. For each variable, the function then computes a weighted mean across neurons and calculates the corresponding weighted variance of the neuron codebook values around that mean. Therefore, variables with high weighted map variance are those whose codebook values vary strongly across the SOM surface. We retain this measure because a variable may vary strongly among SOM neurons even when it is not the strongest discriminator of the final delimitation clusters. This is especially relevant in multilayer delimitation, where some predictors may structure geographic clines, environmental gradients, or within-cluster heterogeneity that are biologically meaningful but not perfectly aligned with the final cluster cut. The second summary, $\eta^2$ (eta squared), quantifies how strongly each variable separates the inferred neuron clusters within a retained SOM replicate. For each variable, the function aligns the neuron-level cluster labels to the rows of the corresponding codebook matrix, removes invalid rows, and derives neuron weights as the number of mapped samples per neuron plus a baseline value of 1 (similar to the map variance calculation). It then calculates a weighted grand mean for the variable across all valid neurons, a weighted total sum of squares around that grand mean, weighted cluster means, and a weighted between-cluster sum of squares. $\eta^2$ is then computed as the ratio of weighted between-cluster sum of squares to weighted total sum of squares, yielding an ANOVA-like effect size measure (Cohen, 1973; Richardson, 2011). Larger $\eta^2$ values indicate that a greater fraction of the total weighted variation in neuron codebook values is explained by the inferred neuron clusters (i.e., by the selected delimitation partition for that replicate). Because each neuron's weight equals its number of mapped samples plus 1, the total weight of a cluster equals the number of samples mapped to its neurons plus the number of neurons assigned to that cluster. Consequently, $\eta^2$ is weighted by both sample support and the number of neurons occupied by each cluster. Their relative influence depends on the distribution of samples and neurons and cannot generally be assumed to be dominated by sample counts. Therefore, map variance asks which variables vary most strongly among SOM neurons, whereas $\eta^2$ asks which variables most strongly distinguish the inferred delimitation groups. This distinction is useful because a variable can be highly structured across the map while contributing mainly to within-cluster gradients or smooth transitions rather than to the selected delimitation boundary itself. One limitation of $\eta^2$ is that it provides an overall estimate of cluster separation without distinguishing among different clusters within a given K. For example, a variable may strongly separate cluster 1 from cluster 2 while contributing much less to separation involving cluster 3. For each layer, replicate-specific map-variance vectors are stored in *mapvar_across_replicates*, and replicate-specific $\eta^2$ vectors are stored in *etasquared_across_replicates*. These are combined across retained SOM replicates into replicate-by-variable matrices, after which the median value for each variable is computed across replicates. The resulting median vectors are stored as the layer-specific outputs: *median_map_variance_variable_importance* and *median_etasquared_variable_importance*. If a replicate yields only K = 1, $\eta^2$ cannot be computed because no between-cluster partition exists. Overall, we interpret $\eta^2$ importance primarily as a summary of cluster-separation signal and map-variance importance primarily as a summary of variation among codebook vectors across the trained SOM layers. The latter might be an especially useful variable importance measurement if $\eta^2$ cannot be calculated (as in the case of K = 1).

7. **Output.** Our *clustering.SOM* function extends the object returned by *train.SOM* by adding the clustering, assignment, replicate-filtering, and variable-importance results used for downstream interpretation. After optional filtering of poorly fitting SOM replicates, the function stores the retained replicate indices in *retained_replicates*, the removed replicate indices in *removed_replicates*, the number of retained replicates in *N_replicates_retained*, and the retained replicate-wise error diagnostics in *quantization_error_retained* and *topographic_error_retained* when available. It also stores the quantile thresholds used for replicate filtering (*quantization.error.quantile* and *topographic.error.quantile*). It adds the replicate-specific clustering results, including the codebook-vector cluster assignments, the sample-level assignment matrix (*cluster_assignment*), the replicate-specific sample-level best-matching-unit and cluster assignments (*cluster_gridcell_assignments*), the replicate-specific soft ancestry matrices (*replicate_ancestry_matrices*), the replicate-wise inferred cluster numbers across retained replicates (*optim_k_vals*), and the corresponding support summaries for alternative values of k, including BIC-based, Davies–Bouldin index, or Silhouette-based results where relevant. The function stores the counts and proportions of replicates supporting each selected value of K (*optim_k_summary*), the relabeled individual-level replicate-consensus ancestry summary (*ancestry_matrix*), the assignment diagnostics *mean_assignment_margin* and *mean_normalized_assignment_entropy*, the replicate-specific soft ancestry matrices when calculated, and the variable-importance metrics *median_etasquared_variable_importance* and *median_map_variance_variable_importance* for each layer.

#### M6: Visualization and interpretation of *delimSOM* output

Our *delimSOM* 2.0 *R* package includes plotting functions corresponding to the main inferential stages of the analysis: SOM learning, layer weights, layer importance, K evaluation, assignment structure, spatial projection, and variable importance. These visualizations serve as diagnostic tools for examining SOM learning and map structure, evaluating support for candidate delimitations, identifying variables and layers associated with cluster separation or variation among SOM neurons, and assessing the biological and geographic patterns of the resulting assignments.

1. **General plotting implementation.** All plotting functions in *delimSOM* 2.0 operate on the *SOM.output* object and extract the relevant stored components for visualization. They support optional file output through the arguments *save*, *overwrite*, *plot.type*, *file.name*, *width*, *height*, and *resolution*. When saving is requested, the functions open the requested graphics device (svg, png, or jpg), draw the plot, and close the device with the *dev.off* function in the *base R grDevices v.4.5.1* package (R Core Team, 2025). We use perceptually uniform and colorblind-friendly color palettes (*col.pal* argument) across plotting functions via the *viridis v.0.6.5 R* package (Garnier et al., 2023). General margin controls of the plots can be specified via the *bottom.margin, left.margin, top.margin, and right.margin* arguments. The functions manage or restore plotting parameters so that graphical settings are controlled consistently during plotting.

2. **Structure-like barplots.** Our *plot.structure.SOM* function displays the individual-level *ancestry_matrix* as a Structure-like stacked barplot, providing an admixture-style visualization similar to those commonly used in *STRUCTURE*-like population genetic analyses (Pritchard et al. 2000). Main arguments include the size of sample labels (*Individual.labels.font.size* argument), the

y-axis title (*Y.axis.title*), the cluster column used to determine the starting point of the individual ordering (*sort.by.col* argument; default: *1*), and optional borders between bars (*bar.border.col* and *bar.border.lwd* arguments). When *sort.by.col* is supplied, each individual is grouped according to the cluster for which it has the highest assignment coefficient. Individuals are then grouped by dominant cluster, beginning with the cluster specified by sort.by.col and continuing through the remaining clusters in column order. Within each dominant-cluster group, individuals are ordered from highest to lowest assignment coefficient for that cluster. For example, with three clusters, *sort.by.col = 1* produces the order Cluster 1, Cluster 2, and Cluster 3, whereas *sort.by.col = 2* produces Cluster 2, Cluster 3, and Cluster 1. When *sort.by.col = NULL*, Euclidean distances among complete assignment profiles are calculated with the *dist* function in *base R* stats, and individuals are ordered by single-linkage hierarchical clustering with the *hclust* function in *base R* stats. The function generates a Structure-like plot with the *plot.Structure* custom function that first converts the assignment coefficients to cumulative proportions and then draws each stacked bar segment as a polygon. Internal segment borders are suppressed (*border = NA*), and optional vertical separator lines between individuals are added with segments when *bar.border.col* is specified, thereby emphasizing relative assignment coefficients rather than within-bar outlines.

3. **Learning-progress plots.** Our *plot.learning.SOM* function visualizes the training trajectories stored in *learning_values_list*. This shows whether SOM learning stabilizes smoothly or whether some layers retain erratic changes late into training. A rapid initial decline followed by a plateau in the training change values suggests the map has converged toward a coherent representation, whereas erratic trajectories or the absence of a clear plateau indicate insufficient training steps, problematic scaling, or weakly structured input data. Function-specific arguments include the line transparency (*lines.alpha* argument), line thickness (*lines.thickness* argument), a user-specified title (*plot.title* argument), legend position (*legend.position* argument), legend line thickness (*legend.lines.thickness* argument), and axis labels (*x.axis.label* and *y.axis.label* arguments). The function determines a global y-axis range across all layers, initializes a single base plot, assigns one color per layer, and overlays the replicate-specific change curves for each layer across the training iterations. Within each layer, all replicate trajectories are plotted with the same layer color, and transparency is applied with the *adjustcolor* function in *base R grDevices* so that repeated overlaid trajectories remain visible without obscuring one another. The legend is added using the stored layer names (*input_data_names*).

4. **Layer-distance-scale plots.** Our *plot.layer.distance.scale.SOM* function visualizes the average pairwise distance scale of each input layer. Such a summary is useful because input layers often differ strongly in dimensionality, scale, variance structure, and overall distance magnitude, so that some layers could otherwise dominate map formation simply because they generate larger raw inter-object distances rather than because they contain more biologically informative structure (Kohonen, 1998, 2001; Wehrens & Kruisselbrink, 2018). This is done across multilayer SOM replicates by transforming the replicate-specific entries in *distance_weights_matrix* back to average pairwise distances, calculating their mean across replicates, and plotting these values as a barplot ordered from highest to lowest layer-distance scale. Function-specific arguments include the plot title (*plot.title* argument) and y-axis label (*y.axis.label* argument). Relatively large values typically indicate layers with many variables, large variances, or data structures that inherently generate large aggregate distances (e.g., SNP or binary layers). During SOM training, layers with large mean pairwise distances receive smaller internal distance-normalization weights, whereas layers with smaller distances receive larger weights, reducing the influence of raw distance magnitude on the aggregated multilayer SOM distance. The plot is a diagnostic of whole-layer distance structure rather than biological importance because high values indicate large raw distances before normalization, whereas low values do not imply weak delimitation signal. The plot should be treated as a diagnostic visualization of whole-layer distance structure and relative scale differences among layers. In the

original implementation (Pyron, 2023; Pyron et al., 2023), layer weighting was visualized from the matrix of replicate-specific internal layer weights (*d_mat*) by plotting transformed mean entries as relative layer weights. *delimSOM* 2.0 instead takes the reciprocal of each replicate-specific weight to recover the layer-distance scale and then averages these values across replicates, avoiding the additional square-root transformation and presenting the result directly as average pairwise distance.

5. **Layer importance plots.** Our *plot.layer.importance.varimp.SOM* and *plot.layer.importance.leaveoneout.SOM* functions visualize layer importance. The first summarizes the variable-importance values within each layer, whereas the second measures how strongly the inferred clustering changes when one layer is omitted and the analysis is rerun. Together, these functions assess whether variables within a layer are associated with cluster separation or variation across the trained SOM map and whether omitting that layer changes the inferred clustering.

    a. ***plot.layer.importance.varimp.SOM.*** This function provides an overview of the two layer-specific variable-importance outputs returned by our *clustering.SOM* function: *median_etasquared_variable_importance* and *median_map_variance_variable_importance* (see Supplementary Methods M5.6). The first measure summarizes the median $\eta^2$ values of variables within each layer across retained SOM replicates and therefore reflects how strongly variables in that layer are associated with separation among the inferred neuron clusters. The second summarizes the median weighted map-variance values of variables within each layer across retained replicates, reflecting how strongly variables in that layer are associated with broader variation across the trained map. These two summaries represent complementary views of post-clustering layer importance: $\eta^2$ emphasizes cluster-separation signal whereas map variance emphasizes variation across the trained map. The function extracts these stored layer-specific outputs and summarizes the distributions of stored variable-importance values across layers. The function plots η2, map variance, or both according to which stored importance summaries contain valid values. Function-specific arguments include separate titles for the $\eta^2$ and map-variance panels (*etasquared.title* and *mapvariance.title* arguments), separate y-axis labels (*etasquared.y.axis.label* and *mapvariance.y.axis.label* arguments), title and axis font sizes (*plot.title.font.size*, *axis.labels.font.size*, and *axis.ticks.font.size* arguments), optional boxplot whiskers (*add.boxplot.whiskers* argument), and ordering of layers by median importance (*sort.by.median* argument). The function returns a layer-summary table of post-clustering importance (*layer_importance_summary_table*) containing the mean and standard deviation of the stored per-variable median importance values within each layer.

    b. ***plot.layer.importance.leaveoneout.SOM***. This function estimates layer importance via replicate-matched leave-one-layer-out analyses, thereby assessing how strongly the inferred delimitation changes when one layer is removed. The function takes a clustered multilayer *SOM_output* object, extracts the stored training and clustering settings, and then reruns matched single-replicate SOM and clustering analyses while omitting one layer at a time. To ensure comparability to the baseline retained replicates, the function overrides training to use one replicate without parallelization, uses matched training and clustering seeds, and performs clustering with replicate filtering disabled. It then compares the baseline (i.e., original) retained replicate to the corresponding leave-one-layer-out analysis for each omitted layer and summarizes the effect of dropping that layer on inferred structure. At the replicate level, the current function records three main change diagnostics. *baseline.modal.k* and *leave.one.layer.out.modal.k* contain the selected K for the single matched baseline and leave-one-layer-out replicate, respectively. *absolute.k.deviation* is the absolute difference between these two K values and measures how much the inferred number of clusters changes after removing the layer. A value of 0 indicates that the inferred number of clusters is unchanged, whereas larger values indicate a greater change in the inferred number of clusters. *pairwise.coassignment.change* measures changes in cluster composition and is calculated as the proportion of shared sample pairs whose same-cluster versus different-cluster relationship changes after removing the layer. A value of 0 indicates that the grouping of individuals is unchanged, whereas larger values indicate that more pairwise grouping relationships change. *baseline.mean.assignment.margin* and *leave.one.layer.out.mean.assignment.margin* measure the sharpness of individual cluster assignments before and after layer removal. For each individual, the assignment margin is the difference between its highest and second-highest soft cluster-assignment probabilities, such that a larger margin indicates stronger support for the best-supported cluster relative to the next-best cluster. The mean assignment margin is then calculated across individuals for each analysis. *delta.mean.assignment.margin* is calculated as the baseline mean assignment margin minus the leave-one-layer-out mean assignment margin. Positive values therefore indicate that removing the layer makes individual cluster assignments less distinct, whereas values near 0 indicate little change in assignment sharpness. Larger absolute K deviations, larger pairwise co-assignment changes, and larger positive assignment-margin changes indicate that the inferred delimitation depends more strongly on the omitted layer. The *layer.summary* table reports the number of attempted and successful comparisons and the mean and median absolute K deviation, pairwise co-assignment change, and assignment-margin change for each omitted layer. The function plots the replicate-level distributions of absolute K deviation (*absolute.k.deviation*; left panel), pairwise co-assignment change (*pairwise.coassignment.change*; middle panel), and assignment-margin change (*delta.mean.assignment.margin*; right panel) for each omitted layer, and returns *layer.summary* and *replicate.matched.results* in a *leave.one.layer.out.results* object. Function-specific arguments include optional replicate-level points (*add.points*, *point.cex*, and *point.alpha* arguments), plot title (*title* argument), panel-specific y-axis labels (*absolute.k.deviation.y.axis.label*, *pairwise.coassignment.change.y.axis.label*, and *assignment.margin.change.y.axis.label* arguments), title and axis font sizes (*title.font.size*, *axis.labels.font.size*, and *axis.ticks.font.size* arguments), optional boxplot whiskers (*add.boxplot.whiskers* argument), whether progress messages are printed (*verbose* argument), progress-message frequency (*message.N.replicates* argument), and controls for saving or reloading the leave-one-layer-out analysis results (*save.leave.one.layer.out.results*, *save.leave.one.layer.out.results.name*, and *overwrite.leave.one.layer.out.results* arguments).

6. **K-evaluation plots.** Our *plot.K.SOM* function visualizes replicate support for alternative cluster numbers. These plots summarize how support changes across alternative values of K and help distinguish between strong support for a single delimitation level and a more gradual structure in which multiple K values remain plausible. We include this visualization because uncertainty in K is one of the main sources of uncertainty in species delimitation and should be assessed directly rather than reduced to a single chosen value. For clustering methods based on BIC (*kmeans+BICelbow*, *kmeans+BICthreshold*, and *GMM+BICthreshold*), the function plots BIC-based support profiles, a ΔBIC panel summarizing replicate-specific BIC improvements between successive K values when max.k > 2, and the frequency of selected K values. For *hierarchical+DB* and *OPTICS+Silhouette*, it plots the Davies–Bouldin index or mean silhouette support together with the frequency of selected K values. For *HDBSCAN*, it plots only the frequency of selected K values across retained replicates. Function-specific arguments include the plot title (*plot.title* argument) and title and axis font sizes (*plot.title.font.size*, *axis.labels.font.size*, and *axis.ticks.font.size* arguments).

7. **SOM-grid visualizations.** Our *plot.model.SOM* function visualizes SOM grid results in map space, allowing the inspection of sample assignment to SOM units, mean codebook distances between

adjacent SOM units, and boundaries between SOM-unit clusters. Function-specific arguments include the color palette for the neighbor-distance panel (*col.pal.neighbor.dist* argument) and cluster panel (*col.pal.clusters* argument), the replicate-selection mode (replicate.mode argument; options: *"first"* or *"representative"*), an optional restriction to replicates with a specified number of clusters (*set.k* argument), the color and line width of cluster boundaries (*boundary.col.clusters* and *boundary.lwd.clusters* arguments), the color, size, and plotting symbol of sample points (*point.col.clusters*, *point.size.clusters*, and *point.shape.clusters* arguments), the cell-shape settings for the cluster and neighbor-distance panels (*cluster.shape.clusters* and *cluster.shape.neighbor.dist* arguments), a horizontal shift applied to the lower cluster plot to align with the upper plot (*shift.plot.clusters* argument), and optional titles for the neighbor-distance and cluster panels (*title.neighbor.dist* and *title.clusters* arguments). If *set.k* is provided, the retained replicate set is restricted to SOM replicates whose inferred number of clusters matches the requested value. Regardless of selection mode, the function displays the selected replicate in a two-panel SOM layout using the *plot* function in *kohonen*. The upper panel shows each SOM unit's mean codebook distance to its adjacent units as a SOM property surface (*type* = *"property"*), highlighting local discontinuities or sharp transitions across the map. This U-matrix visualization is commonly used in SOM studies (Ultsch, 1993; Vesanto & Alhoniemi, 2000; Wehrens & Buydens, 2007). Low neighbor distances indicate smoother local continuity across adjacent map units and are often seen as valleys corresponding to more cohesive regions of the map. High neighbor distances indicate sharp local changes in the codebook vector surface, marking transitions between distinct regions of multivariate structure and potential cluster boundaries (ridges). These features should be interpreted jointly with mapped sample occupancy and the cluster panel, because low-occupancy or empty regions can also arise from topological smoothing and interpolation rather than from strongly supported biological clusters (Vesanto & Alhoniemi, 2000; Wehrens & Buydens, 2007). If low-distance regions (valleys) are not separated by clear boundaries (ridges), this may indicate weaker separation, uncertainty, gradual transitions, intermediate assignments, or discordance among data layers. The lower panel shows sample assignment to SOM cells together with SOM-unit cluster membership (type = *"mapping"*). Boundaries between clustered regions are added with the *add.cluster.boundaries* function in *kohonen*. This visualization shows whether inferred clusters occupy compact contiguous regions of SOM space or whether they are fragmented across the map. The latter indicates weak separation, gradual transitions, or overly fine partitioning. It also allows the topological arrangement of samples on the trained map to be evaluated. Empty or sparsely occupied SOM units are not in themselves evidence of model failure but are expected in many applications, often reflecting regions of map space that are weakly supported by observations or that arise through topological smoothing between more strongly occupied regions (Vesanto & Alhoniemi, 2000; Wehrens & Buydens, 2007). Two *replicate.mode* options are available:

a. ***"first"*.** This mode uses the first retained replicate in the eligible set. This is useful when the goal is simply to inspect one retained SOM realization without additional summarization across replicates. Neighbor distances for the upper panel are calculated for that replicate with the *calc.unit.neighbor.dist* custom function, which calculates codebook distances with the *object.distances* function in *kohonen* using the trained layer-distance functions and weights, identifies immediately adjacent SOM units with the *unit.distances* function in *kohonen*, and computes each unit's mean codebook distance to those adjacent units.

b. ***"representative"* (default).** This mode selects the retained replicate whose SOM-unit clustering produces the most representative sample-level partition among replicates. The *choose.representative.replicate* custom function first identifies the most frequently inferred K among eligible replicates, resolving ties by selecting the smallest K, and then compares the replicates supporting that K. If *set.k* is specified, the eligible replicate set is first restricted to replicates supporting that K. For each comparison, agreement between the sample partitions is quantified with the Adjusted Rand Index. This index is a chance-corrected measure of agreement between two partitions: values near one indicate nearly identical partitions, whereas values near zero indicate agreement close to that expected by chance (Hubert & Arabie, 1985). The mean pairwise Adjusted Rand Index is calculated for each candidate replicate from pairwise comparisons using the *adjustedRandIndex* function in *mclust*, and the replicate with the highest mean agreement is selected as the representative replicate. If no valid pairwise comparisons are available, the function defaults to the first candidate replicate. Neighbor distances for the upper panel are then calculated for that representative replicate with *calc.unit.neighbor.dist*. We use this representative-selection step because SOM clustering is stochastic, so a single retained replicate can still reflect idiosyncratic local variation even after filtering. Selecting the replicate with the greatest average agreement to the other candidate replicates favors a map that best reflects the recurrent clustering structure across runs rather than an arbitrary retained realization. We use this mode as the default because it provides a single retained SOM realization for visualization while reducing the chance that the displayed map is an idiosyncratic stochastic replicate.

8. **Variable-importance plots.** Our *plot.variable.importance.SOM* function visualizes variable importance either as cluster-separation or as map-variance importance, depending on the specified *mode* argument (*"Cluster.separation"* or *"Map.variance"*). Both metrics are described in detail in Supplementary Methods M5.6. The function recalculates these metrics for each retained replicate to generate layer-specific visualizations showing which predictors are most strongly associated either with separation among the inferred SOM-unit clusters or the broader variation of the trained SOM map. Function-specific arguments include outer and inner margin settings (*bottom.margin.total*, *left.margin.total*, *top.margin.total*, *right.margin.total*, *bottom.margin*, *left.margin*, *top.margin*, and *right.margin* arguments), the shared x-axis label (*x.axis.label* argument), the threshold for suppressing variable labels when many bars are present (*bars.threshold.N* argument), font-size controls (*title.font.size, layer.labels.font.size, bar.label.font.size, axis.labels.font.size, and axis.ticks.font.size* arguments), whether boxplot whiskers are added (*add.boxplot.whiskers* argument), the minimum importance shown (*importance.threshold* argument; default: 0.001), and an optional restriction to a specific K (*set.k* argument) in cluster-separation mode. If *mode = "Cluster.separation"* (default), the function determines the inferred number of clusters for each retained replicate and excludes replicates with K = 1 (because no between-cluster separation can then be calculated). If *set.k* is specified, only replicates matching that value are retained. If *mode = "Map.variance"*, all retained SOM replicates are included. For each retained replicate, the function extracts the layer-specific codebook matrices with the *getCodes* function in *kohonen* and calculates the requested metric for each layer (Supplementary Methods M5.6). These per-replicate vectors are assembled into replicate-by-variable matrices for each layer. The function then computes the median importance per variable across retained replicates, removes variables whose median importance does not exceed *importance.threshold*, and sorts the remaining variables by median importance. Plotting is performed as a set of layer-specific horizontal boxplots, with one panel per SOM layer. Whiskers can optionally be suppressed either when *add.boxplot.whiskers = FALSE* or when the number of retained variables exceeds *bars.threshold.N*. Restricting the cluster-separation plot to replicates with a specific K can be useful when one wishes to interpret variable importance conditional on a particular delimitation level.

9. **Assignment maps.** Our *plot.map.SOM* function displays the replicate-consensus assignment coefficients stored in *ancestry_matrix* as pie charts at the geographic coordinates of each sample. The function requires a data frame or matrix supplied through Coordinates containing Latitude and

Longitude columns and matches samples to *ancestry_matrix* using row names. Samples occurring in only one object are omitted, the two objects are reordered to the same shared-sample order, and samples with missing coordinates are excluded. The map extent is determined from the retained coordinates, with the *lat.buffer.range* and *lon.buffer.range* arguments extending the observed latitude and longitude ranges. Using the *map* function in the *maps v.3.4.3 R* package (Becker et al., 2025), a world-map background is drawn, and optional US state and county boundaries are added with the same function. When *preserve.map.aspect = TRUE*, the plotting region and pie radii are adjusted so that the geographic aspect ratio is preserved and pies remain visually circular. Each pie is drawn with the custom *add.admixture.pie.SOM* function, with slice colors following the column order of *ancestry_matrix*. Pies dominated by one color indicate strong replicate-consensus assignment to one cluster, whereas multicolored pies indicate assignment coefficients distributed among multiple clusters. A legend is added with the *add.map.legend.SOM* argument using sequential cluster names (unless names are supplied manually; *legend.cluster.names* argument). A geographic scale bar is added with the *map.scale* function in maps, and a north arrow and its "N" label are drawn with the *arrows* and *text* functions in *base R*. Additional function-specific arguments include the plot title and title font size (plot.title and plot.title.font.size arguments), pie-chart size and color palette (pie.size and pie.col.pal arguments), US state and county boundaries and their line widths (USA.add.states, USA.add.counties, USA.state.lwd, and USA.county.lwd arguments), north-arrow position, length, line width, label offset, and label size (*north.arrow.position*, *north.arrow.length*, *north.arrow.lwd*, *north.arrow.N.position*, and *north.arrow.N.size* arguments), scale-bar position, size, and font size (*scale.position*, *scale.size*, and *scale.font.size* arguments), axis-label font size (*axis.labels.font.size* argument), and legend position, title, text and title font sizes, text italics, enclosing box, and symbol size (*legend.position*, *legend.title*, *legend.text.font.size*, *legend.title.font.size*, *legend.text.italics*, *legend.box*, and *legend.symbol.size* arguments).

##### M7: Simulation set 1: multilayer benchmark simulations of *delimSOM* 2.0 settings

Simulation set 1 was designed to benchmark *delimSOM* 2.0 on synthetic multilayer datasets with known group membership and to evaluate sensitivity to several analytical and model settings. We tested two scenarios representing a relatively low (K = 3) and high number of species (K = 7). Benchmarking was organized around the custom *run.SOM.benchmark* function, which trained replicate SOMs, clustered the resulting codebook vectors, and returned replicate-level performance statistics for each simulated dataset. Unless otherwise specified below, simulation parameter ranges were selected based on preliminary analyses and pilot simulations to ensure that simulated clusters were sufficiently differentiated while retaining variation in clustering difficulty.

1. **Summary of simulation design and main parameters.** Each standard benchmark dataset contained 20 individuals per cluster and four simulated data layers: 1,000 SNP loci, fifteen morphological traits, 30 climate variables, and twelve host categories. The main simulation custom function *simulate.data* accepted the true number of clusters (*N_clusters* argument) and a requested missing-data proportion (*missing_data_prop* argument, default: *0.2*). The function assigned individuals to clusters as evenly as possible and generated individual IDs. Cluster positions for continuous differentiated variables were constructed with the simplex-based *generate.simplex.coordinates* custom function to place cluster means at equal pairwise distances in multivariate space.

2. **Simulation of SNP layer.** SNP data were simulated as diploid 0/1/2 genotype dosages under three locus classes: differentiated loci (D), neutral loci (N), and random-noise loci (R). This mixture of locus classes was used to simulate somewhat realistic genomic datasets containing both structured and unstructured variation, together with stochastic noise. For each simulation replicate, the proportion of differentiated loci was drawn uniformly from 0.15–0.40, and the proportion of random

loci from 0.01–0.06. The remaining loci were treated as neutral. Neutral loci were simulated from a shared allele frequency drawn from a U-shaped beta distribution with the shape parameters alpha = 0.5 and beta = 0.5 using the *rbeta* function in *base R stats*, and then sampled with the *rbinom* function (*size = 2*) in *base R stats*. Random loci were generated similarly, but a randomly selected 1–6% of individuals at each locus were reassigned genotypes sampled uniformly from 0, 1, and 2 to mimic drift-like noise and sequencing or genotyping error (Laurie et al., 2010; Mastretta-Yanes et al., 2015; Nielsen et al., 2011; Pompanon et al., 2005). For each differentiated locus, a target $F_{ST}$ was drawn uniformly from 0.85–1.00, and a global allele frequency was drawn uniformly from 0.25–0.75. Cluster-specific allele frequencies were then drawn from beta distributions parameterized from those values, and cluster-specific diploid genotypes were sampled accordingly. The differentiated loci were iteratively regenerated until the observed mean pairwise $F_{ST}$ across all cluster pairs and all loci fell within a target range of 0.10–0.20 (with an added tolerance of ±0.005). This target $F_{ST}$ range was intended to approximate low to moderate genomic divergence observed among conspecific populations (e.g., Hall, 2022; Hasbún et al., 2016). For each pair of clusters, observed $F_{ST}$ was calculated at each locus as the variance of the two cluster allele frequencies relative to the expected heterozygosity at their mean allele frequency, and values were then averaged across all loci and cluster pairs.

3. **Simulation of morphology layer.** Morphology data were simulated as continuous traits comprising a mixture of differentiated (D) and neutral (N) variables. This design was used to represent realistic phenotypic datasets in which some traits vary primarily within clusters, whereas others contribute to among-cluster separation. For each simulation replicate, the number of differentiated morphology traits was set to K – 1, where K is the true number of simulated clusters, and the remaining morphology traits were treated as neutral. Trait classes were assigned randomly among the morphology columns. Neutral traits were simulated from normal distributions shared across all clusters, with a trait-specific integer mean drawn uniformly from 1 to 100 and a shared simulation-specific standard deviation drawn uniformly from 0.6 to 1.4. Differentiated morphology traits were simulated by first defining cluster-specific trait means in multivariate space, with the separation distance drawn uniformly from 0.8 to 1.8. Specifically, cluster means were arranged using the *generate.simplex.coordinates* custom function as a regular simplex in K − 1 dimensions, so that the pairwise distance between every pair of cluster means equaled the sampled separation distance regardless of K. (e.g., the distance between the means of K1 and K2 was the same as that between K1 and K3 or between K2 and K3). Individual trait values for differentiated morphology traits were then sampled from normal distributions centered on the corresponding cluster-specific means, using the same shared simulation-specific standard deviation applied to the neutral traits. Thus, morphology datasets contained a controlled combination of undifferentiated and cluster-informative traits, with the degree of among-cluster separation determined by the spacing of cluster means and the degree of within-cluster dispersion determined by the shared standard deviation. The morphology distance and standard-deviation settings were chosen to approximate realistic cluster separation relative to multivariate within-cluster trait dispersion (De Kort et al., 2013; Leinonen et al., 2013; Opedal et al., 2023; Siefert et al., 2015; Westerband et al., 2021).

4. **Simulation of climate layer.** Like the morphology layer (see section above), climate data were simulated as continuous environmental variables comprising a mixture of differentiated (D) and neutral (N) predictors. However, we added a skewness parameter so that the simulated distributions could be asymmetric and either left- or right-tailed, introducing realistic non-normality into the simulated environmental data. Neutral climate variables were simulated from skew-normal distributions shared across all clusters using the *rsn* function in the *sn v.2.1.3 R* package, with a variable-specific integer location parameter drawn uniformly from 1 to 100, a shared simulation-specific scale parameter drawn uniformly from 0.6 to 1.4, and a variable-specific skew parameter

drawn uniformly from −6 to 6. The direction of skewness was determined by the sign of the skewness parameter (*alpha*), such that negative values produced left-skewed distributions, positive values produced right-skewed distributions, and values near zero produced approximately symmetric distributions. $K - 1$ differentiated climate variables were simulated by defining cluster-specific location vectors in multivariate environmental space using the same regular-simplex procedure applied to the morphology layer, with the pairwise distance between all location vectors drawn uniformly from 0.8 to 1.8. Individual values for differentiated climate variables were then sampled from skew-normal distributions using the same shared simulation-specific scale parameter but drawing cluster- and variable-specific skew parameters uniformly from −6 to 6. The climate distance and scale settings were intended to approximate realistic among-cluster environmental separation relative to within-cluster multivariate environmental dispersion (Bates & Bertelsmeier, 2021; Broennimann et al., 2012; Carscadden et al., 2020; Liu et al., 2020).

5. **Simulation of host layer.** Host data were simulated as a binary individual-by-host matrix comprising 12 host categories. We used host association as an example of a categorical ecological layer, based on its importance in many herbivorous insect systems; however, this layer could represent any analogous categorical ecological variable, such as habitat type, substrate use, symbiont association, or developmental mode. This design was used to represent realistic ecological datasets in which most individuals within a cluster are associated with one dominant host, whereas the remaining individuals are distributed among one or more secondary hosts. For each simulation replicate, the host column with the same index as the cluster was treated as that cluster's dominant host. The proportion of individuals assigned to the dominant host was drawn uniformly from 0.70 to 0.96. Individuals assigned to the dominant host were selected randomly within each cluster, and their corresponding dominant-host entry was coded as 1 (presence) in the host matrix. The remaining individuals within each cluster were each assigned to one of the secondary host categories selected for that cluster. The number of secondary hosts was generated as one plus a Poisson-distributed random value with $\lambda = 1$ using the *rpois* function in *base R stats*, ensuring a minimum of one. This number was capped at the number of available secondary-host categories, and the actual secondary hosts were sampled without replacement from the set of host categories excluding the dominant host. If only one secondary host was selected, all remaining individuals in that cluster were assigned to that host. If more than one secondary host was selected, uneven secondary-host use was generated by drawing gamma-distributed weights with $shape = 0.8$ and $rate = 1$ using the *rgamma* function in *base R stats*, normalizing those weights to probabilities, and then sampling host assignments for the remaining individuals with replacement according to those probabilities. If those probability weights were invalid, the selected secondary hosts were assigned in a simple recycled sequence across the remaining individuals. Thus, host datasets contained a controlled combination of dominant host association and heterogeneous secondary-host use, with the degree of dominance determined by the dominant-host proportion and the distribution of secondary-host use determined by the number of selected secondary hosts and their gamma-distributed assignment weights. The host simulation settings were chosen to approximate realistic dominant and secondary host-use structure in empirical host-associated systems (Feder et al., 1994; Ramírez-Martínez & Tlapaya-Romero, 2023; Wehmeyer et al., 2024).

6. **Missing data and stored simulation summaries**. After generation of all four layers, missing values were introduced separately within each layer using the *add.NA* custom function, which converted a randomly selected set of cells to *NA*, with the number of selected cells equal to the integer obtained by rounding down the product of the missing-data proportion and the total number of cells. Missing data were therefore imposed separately on the SNP, morphology, climate, and host matrices after simulation of the complete datasets. The simulation function then returned the missing-data versions of all four layers together with the true cluster labels and a simulation summary table (*sim_stats*)

containing selected simulation settings and outcomes. Thus, each simulation replicate retained the four input layers containing missing values, the true cluster labels, and the corresponding simulation summary for downstream benchmarking.

7. **Custom functions to score SOM performance.** Several custom functions were used for benchmark analyses. *extract.QE.TE* calculated quantization error (QE) and topographic error (TE) separately for each layer of a trained SOM replicate and then averaged each metric across layers (Supplementary Methods M4.12). *get.accuracy* calculated the proportion of correctly assigned individuals after matching inferred cluster labels to true labels using the Hungarian algorithm (Supplementary Methods M5.5), provided that the inferred and true numbers of clusters were equal. *get.cluster.labels.for.replicate* extracted the clustering solution for a given SOM replicate, *add.sim.stats* appended simulation metadata to result tables, *make.failed.stats.df* returned placeholder rows when training or clustering failed, and *bind.results.with.sim* combined per-simulation result tables while preserving original simulation indices. *simulate.standard.datasets* repeatedly generated datasets with *simulate.data* using a sequence of random seeds and retried failed simulations with successive seeds until the requested number of successful datasets had been obtained.

8. **Replicate-level SOM benchmarking procedure**. The main inputs to the custom *run.SOM.benchmark* function were an aligned list of simulated data matrices, the true cluster labels, the name and value of a grouping variable describing the benchmark condition being tested, the clustering method, the SOM neighborhood function, the maximum tested K, and whether learning-rate tuning was enabled. It then trained and clustered SOMs with our *train.SOM* and *clustering.SOM* functions using the benchmark-wide settings. Combined SOM training and clustering time was measured with the *system.time* function in *base R* and retained as the total runtime for each simulated dataset. For each replicate, the function stored the inferred number of clusters (*K_inferred*), adjusted Rand index (*ARI*), assignment accuracy (*Acc*; only when the inferred K matched the true k), a logical indicator of whether the inferred K was correct (*K_correct*), a logical indicator of whether inferred K was off by at least two (*K_far_off*), and the layer-averaged QE and TE.

9. **Common benchmark settings for all simulation experiments**. The comparative simulations were conducted separately for datasets with true K = 3 and true K = 7. Within each benchmark experiment, all SOM settings were held at the benchmark defaults described above except for the focal setting being tested. The *set.seed* function in *base R* was used to ensure reproducibility, and a common simulation-seed vector was used to generate the base datasets for each K scenario. These base datasets were then reused across benchmark experiments so that the same simulated datasets were analyzed under the tested settings.

10. **Effect of clustering method.** To test the effect of the clustering and K-selection approaches used after SOM training, we compared the six clustering and K-selection methods implemented in clustering.SOM (Supplementary Methods M5.4). Each method was evaluated using the same base simulated datasets while all training settings were held constant, thereby evaluating the effect of the post-SOM clustering and K-selection approach.

11. **Effect of SOM iterations.** To test the effect of the number of SOM training iterations, we varied *N.steps* across seven values: 20, 50, 100, 200, 500, 1,000, and 2,000.

12. **Effect of missing-data proportion.** To test robustness to structured missing data, we varied the target missing-data proportion within selected high-missingness individuals and variables across nine values: 0.0, 0.1, 0.2, 0.3, 0.4, 0.5, 0.6, 0.7, and 0.8. For each simulated dataset, the proportions of high-missingness individuals and variables were independently drawn from normal distributions with a mean of 0.20 and a 95% interval of 0.05–0.35, truncated to values between 0 and 1. Within each data layer, the specified target proportion of entries was then converted to NA for every selected high-

missingness individual and variable. Fixed simulation-specific seeds were reused across missing-data proportions so that the same high-missingness individuals and, within each layer, the same high-missingness variables were selected across the tested conditions.

13. **Effect of SOM neighborhood function.** To test the effect of the neighborhood function used during SOM training, we compared the *"gaussian"* and *"bubble"* options while holding the simulated datasets and all other SOM settings constant.
14. **Effect of codebook learning-rate tuning.** To test whether automatic codebook learning-rate tuning affected SOM performance, we compared *learning.rate.tuning = TRUE* and *FALSE* using the same simulated datasets while holding all other SOM-training and clustering settings constant.
15. **Output summaries and visualization across experiments.** For each benchmark, results were grouped by the focal experimental factor and summarized using mean ARI, mean inferred K, mean assignment accuracy, mean runtime, failed rows, and total result rows; mean QE and TE were also reported where relevant. ARI, assignment accuracy, QE, TE, and runtime were first averaged across SOM replicates within each simulated dataset and condition. Each continuous response was then analyzed separately using a Gaussian mixed-effects model fitted with the *lme* function in the *nlme v.3.1.168 R* package (Pinheiro et al., 2025; Pinheiro & Bates, 2006), with the focal experimental factor as a categorical fixed-effect predictor and simulation identity as a random intercept. The binary responses indicating exact recovery of K and deviation from the true K by at least two were analyzed separately using binomial mixed-effects models fitted with the *glmmTMB* function in the *glmmTMB v.1.1.14 R* package (Brooks et al., 2017), with the focal experimental factor as the fixed-effect predictor and random intercepts for simulation identity and the simulation-by-condition combination. Estimated marginal means and their one-standard-error intervals were calculated with the emmeans function in the *emmeans v.2.0.1 R* package (Lenth & Piaskowski, 2025) and displayed together with simulation-level values.

#### M8: Simulation set 2: *fastsimcoal2* demographic simulations and comparison with alternative population-genetic clustering methods

Simulation set 2 was designed to evaluate *delimSOM* 2.0 performance on genomic data under two-population/species (K = 2) demographic scenarios spanning a range of divergence times, migration rates, sample sizes, and uneven sampling. *delimSOM* 2.0 performance was then compared with commonly used alternative population-genetic clustering methods, namely *DAPC*, *sNMF*, and *STRUCTURE* (see below). Although this SNP-only comparison does not take advantage of the integrative, multilayer nature of *delimSOM* 2.0, it was included to place its performance in the context of commonly used population-genetic clustering methods. SNP data were simulated in *fastsimcoal2*, a continuous-time coalescent-based simulator that models genomic data under complex demographic histories from the site frequency spectrum under a maximum-likelihood framework (Excoffier et al., 2021). Simulation settings broadly followed previous studies that have used *fastsimcoal2* for simulation-based analyses (DeRaad et al., 2022; Smith & Carstens, 2020; Smith & Hahn, 2024). However, rather than drawing parameters from broad uniform priors, we used fixed values or parameter grids to isolate the effects of divergence time and migration rate within each sampling treatment while separately evaluating the effects of sample size and uneven sampling.

1. **Demographic simulation design.** Unlinked SNP data were simulated to represent genetic divergence between two populations. Divergence time (*TDIV*) was varied across 251 evenly spaced values from no divergence (*TDIV = 0*) to 200,000 generations. This range was selected based on preliminary analyses to ensure that the simulations covered the full range from no detectable

population structure to strong genetic differentiation. The two demes were assigned equal effective sizes of 500,000 haploid gene copies each (corresponding to 250,000 diploid effective individuals per deme). Growth rates were set to zero in both demes. For all simulations, we specified 100,000 independent chromosomes, one contiguous DNA block per chromosome, no recombination, and a mutation rate of $10^{-8}$. In addition to the no-gene-flow scenario (m = 0), we evaluated *delimSOM* 2.0 under gene flow by simulating secondary-contact scenarios using three symmetric migration rates (fastsimcoal2 migration parameter $m = 1 \times 10^{-6}$, $4 \times 10^{-6}$, and $7 \times 10^{-6}$). Together with the no-gene-flow treatment, the four migration rates corresponded to approximately 0, 0.25, 1, and 1.75 diploid immigrants per generation per direction, given the simulated deme size. Migration was represented by two migration matrices: a matrix containing the symmetric migration rate between demes and a second matrix containing no migration. Secondary contact occurred halfway between divergence and the present, after which gene flow continued to the present. The combination of 251 divergence times and four migration treatments yielded 1,004 simulated datasets. All demographic, chromosome, recombination, and mutation settings were specified in the *fastsimcoal2* template file.

2. **Sample size and uneven sampling.** First, we tested the effect of different sample sizes by analyzing high (24), medium (16), and low (8) numbers of diploid individuals per population (even sampling). Second, we evaluated the effect of uneven sampling by producing a 3:1 sampling ratio while maintaining the same total sample sizes. Together, this led to three even ("symmetric") and three uneven ("asymmetric") subsets. The same individual subsets were used across *delimSOM* 2.0 and the alternative clustering methods within each sampling treatment to permit direct comparisons among methods.

3. **Generation of simulated SNP datasets.** Simulations were run in a Singularity container on the Morgan Compute Cluster (University of Kentucky) using the *fsc28* command. The *fastsimcoal2* template specified 100,000 independent chromosomes for each simulation. We deliberately oversimulated loci and then downsampled them so that every final dataset contained the same number of SNPs, thereby standardizing marker number across demographic scenarios and preventing differences in locus count from confounding downstream clustering performance. Each *fastsimcoal2* output *.gen* file was converted into a VCF file using a custom *awk* parser that wrote a minimal VCF header, converted genotype codes (0, 1, and 2) to diploid VCF genotypes (0/0, 0/1, and 1/1), skipped malformed rows, and randomly downsampled variants to exactly 1,000 SNPs using reproducible seeds. Downsampled variants were re-sorted by chromosome and position. This procedure produced 1,004 final VCF files, with filenames encoding the migration-treatment index, divergence time, and migration value.

4. **Import of VCFs.** VCF files were imported into *R* and converted to *genind* objects using the *read.vcfR* and *vcfR2genind* functions in *vcfR*. Individuals were assigned to the two simulated populations based on the *G_1_* and *G_2_* prefixes in their sample names. For each sampling treatment, individuals were selected through reproducible random subsampling using a base seed of 1. Sampling seeds were defined by divergence-time group and population so that the same individuals were retained across migration treatments for a given divergence time and population. The same sampled individuals were then used across clustering methods within each sampling treatment

5. ***delimSOM* 2.0 analysis.** Each processed SNP matrix was analyzed by training 110 replicate SOMs with 100 training iterations per replicate using *train.SOM*. A grid multiplier of four was used for all six sampling treatments. The resulting codebook vectors were analyzed with *clustering.SOM* using the recommended k-means clustering and BIC elbow method (*clustering.method = "kmeans+BICelbow"*), with the candidate range restricted to K = 1–2 and the default BIC threshold of 6. Poorly fitting SOM replicates were removed using the default quantization- and topographic-error filters before support was summarized. For each dataset, the

*optim_k_summary* was retained, and recovery of the expected two-population solution was quantified as the proportion of retained replicates selecting K = 2.

6. **Comparison with alternative methods.** We compared *delimSOM* 2.0 with three alternative population-genetic clustering methods: *DAPC*, *sNMF*, and *STRUCTURE*. Each method was applied to the same simulated VCF datasets and individual subsets within each sampling treatment, using method-specific input preparation and model-selection criteria. For all three alternative methods, K = 1 and K = 2 were evaluated, and performance was quantified according to whether the expected two-population solution (K = 2) was recovered.

    a. ***DAPC.*** Discriminant analysis of principal components (*DAPC*) is a multivariate method commonly used to identify and describe genetic clusters (Jombart et al., 2010; Miller et al., 2020). When groups are unknown, the *adegenet DAPC* workflow infers genetic clusters by applying k-means clustering to principal-component scores and comparing alternative values of K using BIC. We followed this de novo clustering stage of the *DAPC* workflow following recommendations (Jombart, 2022) using the *adegenet v.2.1.11 R* package (Jombart & Ahmed, 2011) to infer whether each simulated dataset supported one or two genetic groups. The *adegenet* workflow generally recommends selecting K by inspecting the BIC curve and retaining additional clusters only while they provide an appreciable improvement in BIC (Jombart, 2022). Because our comparison was restricted to K = 1 and K = 2, such inspection of the BIC curve was not possible. We therefore defined an appreciable improvement as a decrease in BIC of at least four units, a pragmatic threshold that falls within the range conventionally interpreted as positive evidence for the lower-BIC model (Kass & Raftery, 1995). Accordingly, K = 2 was selected when its BIC was at least four units lower than that of K = 1, while otherwise K = 1 was selected.

    b. Because our comparison focused solely on the inference of K, the subsequent supervised discriminant-analysis step of *DAPC* was not performed.

    c. ***sNMF***. Sparse nonnegative matrix factorization (*sNMF*) estimates individual ancestry coefficients from multilocus genotype data and is implemented in the *LEA v.3.22 R* package (Frichot et al., 2014; Frichot & François, 2015). Model performance was evaluated using cross-entropy, which measures how accurately masked genotypes are predicted under a given value of K, with lower cross-entropy values indicating greater predictive accuracy (Frichot et al., 2014). Following application of each sampling treatment, the simulated VCF genotypes were converted to the *LEA .geno* format, with diploid genotypes represented as 0, 1, or 2 copies of the alternative allele and missing genotypes represented as 9. We evaluated K = 1 and K = 2 because the objective was to distinguish the absence of detectable population structure from recovery of the two simulated populations. For each value of K, we performed 50 independent runs with a ploidy of two and cross-entropy calculation. Multiple runs were performed because the stochastic optimization can produce different solutions depending on its initialization. For each value of K, the run with the lowest cross-entropy was retained. The K = 2 solution was selected only when its minimum cross-entropy was lower than that obtained for K = 1, thereby requiring the more complex two-population solution to improve predictive performance. If this criterion was not met, K = 1 was selected.

    d. ***STRUCTURE.*** *STRUCTURE* is a Bayesian model-based clustering method used to infer population structure and individual ancestry from multilocus genotype data (Pritchard et al., 2000). First, the simulated VCF files were converted to *STRUCTURE* input format using *VCFtools* v.0.1.16 (Danecek et al., 2011) and a custom *awk* script. Diploid genotype dosages of 0, 1, and 2 were converted to the allele pairs 1/1, 1/2, and 2/2, respectively. Missing genotypes were represented as 0/0. Population identifiers were included in the input files to retain the known simulated origins of individuals but were not used as prior information during clustering (*USEPOPINFO = 0*). Analyses were performed using *STRUCTURE v.2.3.4* under the admixture model (*NOADMIX = 0*), with correlated allele frequencies among populations (*FREQSCORR = 1*) and no linkage model (*LINKAGE = 0*) (Falush et al., 2003). We evaluated K = 1 and K = 2 using ten independent runs per value of K. Each run consisted of a burn-in period of 200,000 iterations followed by 800,000 Markov chain Monte Carlo iterations. Runs for K = 1 and K = 2 were matched by replicate number. For each dataset and value of K, the estimated log probability of the data *ln P(D)* was averaged across the ten independent runs to reduce variation among MCMC runs and obtain a more stable measure of model support. The commonly used ΔK method (Evanno et al., 2005) was not applicable in our comparison because it cannot evaluate K = 1 and requires at least three successive values of K. We therefore compared the mean *ln P(D)* values directly. Selecting K = 2 whenever its mean *ln P(D)* exceeded that of K = 1 resulted in frequent inference of two clusters in simulations with zero divergence, where no population structure was present. Because no established threshold exists for distinguishing K = 1 from K = 2 based directly on differences in mean *ln P(D)*, we adopted 50 log-probability units as a pragmatic, conservative cutoff. This threshold was chosen based on preliminary test runs, in which it effectively reduced spurious inference of K = 2 under the no-divergence condition while retaining sensitivity to population structure in divergent simulations. Accordingly, K = 2 was selected when its mean *ln P(D)* exceeded that of K = 1 by more than 50 units; otherwise, K = 1 was selected. *STRUCTURE* analyses were run on the Morgan Compute Cluster.

7. **Evaluation of K = 2 recovery.** Recovery of the expected two-population solution was evaluated separately for each method. For *delimSOM*, support for K = 2 was quantified as the number and proportion of retained SOM replicates selecting K = 2. *DAPC*, *sNMF*, and *STRUCTURE* each produced one binary K = 1 or K = 2 decision per dataset.

8. **Genetic differentiation.** To evaluate the degree of genetic differentiation between the simulated populations, overall Weir and Cockerham $F_{ST}$ (Weir & Cockerham, 1984) was calculated for each simulated dataset and sampling treatment. $F_{ST}$ was calculated from the subsampled *genind* object before preprocessing, using the same sampled individuals as the clustering analyses while retaining all SNPs. Variance components were calculated separately for each biallelic locus, and the multilocus $F_{ST}$ estimate was obtained by dividing the sum of the among-population variance component across loci by the sum of the among-population, among-individual within-population, and within-individual variance components across loci.

9. **Result summaries and statistical comparison.** Results were summarized separately for each of the three even and three uneven sampling treatments. Per-dataset output tables retained the simulated demographic and sampling treatments, sample sizes, numbers of retained loci, $F_{ST}$ estimates, analysis-status information, *delimSOM* support proportions for K, *DAPC* clustering results, *sNMF* cross-entropy values, and *STRUCTURE* log-probability estimates. To estimate the divergence time at which the probability of recovering the expected two-population solution reached 0.5, binomial logistic regressions were fitted separately for each combination of clustering method and migration rate within each sampling treatment using the *glm* function in *base R stats*. For *delimSOM*, the response consisted of the numbers of retained SOM replicates selecting and not selecting K = 2. For *DAPC*, *sNMF*, and *STRUCTURE*, each simulated dataset contributed a single binary outcome indicating whether K = 2 or K = 1 was selected. The numbers of outcomes selecting and not selecting K = 2 were modeled as a binomial response, with divergence time (*TDIV*) as the predictor and a logit link function. The fitted intercept and slope were extracted using the *coef* function in *base R stats*. For each fitted model, the divergence time corresponding to a predicted probability of 0.5 was calculated by solving the fitted logistic-regression equation for divergence time.

**M9: Empirical evaluation using multi-layer case studies**

We evaluated *delimSOM* 2.0 using eight empirical multi-layer case studies spanning diverse taxonomic systems and different numbers of individuals and candidate taxa.

1. **Common preparation.** Data were obtained from the source studies and reformatted as individual-by-variable matrices, with consistent individual identifiers used across all layers. When geographic coordinates were available, we used *NicheDiv* (Schönberger et al., 2026) to extract a broad set of environmental variables for inclusion as an environmental data layer. Default settings were used for preprocessing. Genetic inputs were imported and filtered using *process.SNP.data.SOM*. Categorical variables were converted to binary indicator variables using *make.cols.binary.SOM*. Continuous morphological and environmental variables were processed using *remove.lowCV.multicollinearity.SOM* to remove variables with low variation and highly correlated predictors. Spatial layers included latitude, longitude and elevation. To detect finer taxonomic substructure, we subsequently performed hierarchical reanalyses within each of the main recovered clusters (Janes et al. 2017). The number of samples and variables before and after processing are summarized in Supplementary Table S2.
2. **SOM training and clustering.** Unless otherwise stated, each aligned empirical dataset was analyzed using *train.SOM* with default settings. Then, the codebook vectors were clustered using each of the six implemented clustering and K-selection methods with default settings. Unless otherwise stated, values from K = 1 to K = 10 were evaluated.
    a. ***Polygonia* butterflies.** The first case study (focus study 1 in main text) examined species boundaries among four broadly sympatric anglewing butterflies from western Canada: *Polygonia faunus*, *P. gracilis*, *P. progne*, and *P. satyrus*. Dupuis et al. (2018) found that mitochondrial COI sequences and genome-wide SNP data supported the four species. Phylogenetic analyses recovered each species as reciprocally monophyletic, and *STRUCTURE* supported four genetic clusters corresponding to the four species. Hierarchical analyses detected additional substructure within *P. faunus* and *P. satyrus*, although this structure lacked clear geographic or habitat associations. Their study found no evidence of admixture among the four species, but the mitochondrial and nuclear results showed some cytonuclear discordance. Visually scored wing characters broadly separated the four species, although some overlap remained, particularly between *P. faunus* and *P. gracilis*. In contrast, wing colors did not separate the species in a PCA and showed only partial separation when species assignments from the genetic analyses were used to guide linear discriminant analysis. Overall, Dupuis et al. (2018) evaluated the system using mitochondrial COI sequences, genome-wide SNPs from genotyping-by-sequencing, and two morphological datasets. The first morphological dataset comprised visually scored discrete and ordinal characters describing wing spots and patterns commonly used for field identification. The second comprised continuous wing RGB color measurements obtained from six standardized regions of the dorsal and ventral wing surfaces. Female *P. faunus* and *P. satyrus* were additionally classified into smeared and contrasted morphotypes. In our reanalysis, we included six layers: genome-wide SNPs, mitochondrial COI characters, continuous morphology, categorical morphology, environmental, and spatial data. The continuous morphological layer contained the RGB measurements, whereas the categorical morphological layer contained the visually scored wing characters together with binary indicators of the morphotypes. We separated the continuous and categorical morphological

variables because of their different data structures. We used *NicheDiv* to extract a comprehensive set of environmental variables from the specimen coordinates.

b. ***Desmognathus aeneus*.** The second case study (focus study 2 in main text) examined the seepage salamander *Desmognathus aeneus* from the southeastern United States. Pyron et al. (2024) found deep geographic genetic structure but concluded that the identified lineages represented intraspecific population structure rather than independently evolving species. *DAPC* and *sNMF* recovered three major genetic clusters, whereas the spatially explicit *conStruct* analysis supported two broader clusters after accounting for isolation by distance. The lineages exhibited widespread admixture across contact zones and an approximately continuous relationship between geographic and genetic distance. They also showed little overall phenotypic differentiation and limited adaptive genomic differentiation associated with environmental and morphological variables. These results suggested that recurrent gene flow and stabilizing selection maintained cohesion among deeply divergent phylogeographic lineages. Overall, Pyron et al. (2024) analyzed this system using genomic, spatial, environmental, and phenotypic data. The genomic data consisted of SNPs generated using genotype-by-sequencing. The phenotypic data comprised allometrically size-corrected linear measurements of body, limb, tail, and craniofacial morphology. The environmental data included climatic variables from *WorldClim* and *ENVIREM* together with Level IV ecoregion identity. In our reanalysis, we included five layers: genomic, phenotypic, spatial, environmental, and watershed. The watershed layer contained binary indicators representing Level IV ecoregion identity and watershed classifications. We also replaced the original *WorldClim* and *ENVIREM* variables with a more comprehensive environmental dataset extracted using *NicheDiv*.

c. ***Desmognathus monticola* and *D. cheaha*.** The third case study examined the eastern North American seal salamanders *Desmognathus monticola* and *D. cheaha*. Specifically, this dataset had previously been analyzed using a multilayer SOM implemented in the original version of *delimSOM* (Pyron, 2023). His multi-layer SOM recovered two clusters with strong support, corresponding to the two species. *DAPC* recovered admixed individuals as a third cluster. Random forest recovered additional geographically structured clusters within both species, whereas t-SNE recovered the admixed individuals and additional genetic clusters within *D. monticola*. In contrast, the integrative SOM produced nearly discrete assignments and did not delimit the admixed or geographically structured groups as additional species. (Pyron, 2023) analyzed this system using four data layers representing genomic, spatial, climatic, and phenotypic data. The genomic layer consisted of SNP data generated using genotype-by-sequencing. The phenotypic layer included size-corrected measurements of body size and shape together with larval spot counts. In our reanalysis, we included five separate layers: genomic, phenotypic, spatial, environmental, and watershed. We treated watershed and ecoregion variables as a separate categorical layer (because of its different data structure compared to the continuous environmental data) and replaced the original climate variables with a more comprehensive set of environmental variables extracted using *NicheDiv*.

d. ***Desmognathus valentinei* and *D. pascagoula*.** The fourth case study examined two swamp-dwelling dusky salamanders from the Gulf Coastal Plain of Mississippi and Alabama: *Desmognathus valentinei* and *D. pascagoula*. Pyron et al. (2022) described *D. pascagoula* as a narrowly distributed species that was genetically, morphologically, and geographically distinct from *D. valentinei*. Spatially explicit genomic clustering supported two genetic clusters with little mixed ancestry. Mitochondrial analyses also recovered *D. pascagoula* as a strongly supported lineage sister to *D. valentinei*, while morphological analyses identified differentiation primarily in body proportions and craniofacial traits. Overall, Pyron et al. (2022) evaluated the system using nuclear genomic, mitochondrial, morphological, and geographic evidence. The nuclear genomic data consisted of SNPs generated using genotype-by-sequencing, whereas the mitochondrial data comprised the ND2–COI region. The phenotypic data included size-corrected linear measurements of body, limb, and craniofacial morphology together with qualitative assessments of coloration and pattern. In our reanalysis, we included five layers: genomic, phenotypic, spatial, environmental, and watershed. The phenotypic layer contained the retained continuous morphological measurements. Watershed identity was encoded as binary indicators and included as a separate categorical layer because its structure differed from that of the continuous environmental variables. We also added a comprehensive environmental layer extracted using *NicheDiv*. Individual-level mitochondrial data and qualitative coloration and pattern characters were not available.

e. ***Viburnum* shrubs.** The fifth case study examined the *Viburnum nudum* species complex, a morphologically variable group of woody shrubs distributed across eastern North America (Spriggs et al., 2019). Genetic clustering (*STRUCTURE* and *DAPC*), phylogenetic analyses (*RAxML*, *IQ-TREE*, *tetrad*, and *SplitsTree*), and coalescent species delimitation (*BPP*) consistently supported three major lineages: *V. cassinoides*, *V. nitidum*, and *V. nudum*. The latter was highly divergent from the other two species, whereas *V. cassinoides* and *V. nitidum* were recently diverged. *Viburnum nitidum* and *V. nudum* occurred sympatrically across much of the southeastern United States but remained genetically distinct. Hierarchical *STRUCTURE* analyses detected geographic substructure within each species, but there was limited evidence of admixture. Tests of introgression likewise found little evidence of gene flow. Morphological analyses provided weaker support for the three-species delimitation: unsupervised clustering (normal mixture models) supported either one or two groups and misclassified one quarter of individuals under the two-group model. *Viburnum nudum* differed from the other species in several leaf and inflorescence measurements, whereas *V. cassinoides* and *V. nitidum* overlapped extensively in most measured traits and differed only partially in leaf aspect ratio, peduncle length, and leaf-margin morphology. Overall, Spriggs et al. (2019) evaluated the complex using RADseq, morphology, geography, and species distribution modelling. The morphological data comprised quantitative leaf and inflorescence measurements together with qualitative leaf characters. In our reanalysis, we included genomic and morphological layers. Spatial and environmental data were unavailable.

f. ***Elysia* sea slugs.** The sixth case study examined a cryptic radiation of photosynthetic sea slugs in the western Atlantic *Elysia papillosa* species complex. Krug et al. (2026) evaluated six described species and one previously unrecognized candidate lineage using mitochondrial and nuclear sequence data together with host use, larval development, reproductive anatomy, and geography. Maximum-likelihood (*RAxML*) and Bayesian phylogenetic analyses (*BayesPhylogenies*) of mitochondrial COI sequences recovered the recognized lineages, whereas species-delimitation analyses (*ABGD*, *ASAP*, *GMYC*, and *bPTP*) broadly supported the six described species together with a divergent specimen provisionally identified as *E. cf. zuleicae* from the U.S. Virgin Islands. The barcode-gap methods recovered exactly seven lineages, whereas the coalescent methods frequently oversplit geographically structured populations, particularly in species with less dispersive larvae. Nuclear H3 alleles largely reinforced the COI-based species boundaries, although three specimens from Jamaica exhibited cytonuclear discordance. In our reanalysis, we included four layers: mitochondrial COI DNA, host use and development, environmental variation, and spatial data. Host identity and developmental mode were converted to binary indicators and combined into a single layer. The spatial layer contained latitude, longitude, and sea depth, whereas the environmental layer contained marine environmental variables obtained from *Bio-ORACLE v.2.0* (Assis et al., 2018) using the *bio_oracle* function in the *geodata v.0.6.9 R* package (Hijmans et al., 2024). Nuclear H3 and reproductive-morphology data were not available.

g. ***Pocillopora* corals.** The seventh case study examined species boundaries in Indo-Pacific *Pocillopora* corals, a morphologically variable genus with longstanding taxonomic uncertainty. Oury et al. (2023) identified four major clades and 21 genomic species hypotheses using phylogenomic inference (*RAxML-NG* and *BEAST*), clustering and network analyses (*STRUCTURE*, *sNMF*, *DAPC*, a minimum-spanning tree, and a split network), and coalescent species delimitation (*BFD**). Thirteen of these hypotheses were strongly supported by the combined evidence, whereas six represented undescribed or nominal species that had been incorrectly synonymized. Genomic assignment analyses recovered nearly all genomic species hypotheses and detected introgression among several groups, including one lineage consistent with a hybrid origin. Colony macromorphology was generally unreliable for species identification because morphotypes overlapped extensively among genomic lineages, whereas corallite micromorphology provided more informative diagnostic variation. Overall, Oury et al. (2023) evaluated the system using genomic SNPs, microsatellites, mitochondrial and nuclear marker haplotypes, macro- and micromorphology, biogeography, and algal symbiont community composition. The genomic SNPs were generated through target enrichment of ultraconserved elements and exon loci. The marker data included mitochondrial open reading frame (mtORF) and nuclear PocHistone haplotypes, whereas communities of Symbiodiniaceae (the symbiotic dinoflagellate algae associated with corals) were characterized through internal transcribed spacer 2 (ITS2) metabarcoding. Macromorphology was represented by colony morphotypes based on branch form, verrucae, and growth form. Micromorphology comprised seven continuous measurements and two categorical characters describing septum and columella shape. In our reanalysis, we included eight layers: genomic SNPs, microsatellites, host marker haplotypes, continuous morphology, binary morphology, spatial data, environmental variation, and algal symbiont community composition. The host marker-haplotype layer was constructed from mtORF and PocHistone haplotypes, which were treated as categorical variables and converted to binary indicators. Continuous morphology comprised the quantitative micromorphometric measurements, whereas binary morphology comprised septum and columella shape and colony morphotype. We separated the continuous and binary morphological variables because of their different data structures. The spatial layer contained latitude and longitude of the sampling localities. The environmental layer contained marine environmental variables obtained from surface *Bio-ORACLE v.2.0* layers (Assis et al., 2018) using the *bio_oracle* function in the *geodata v.0.6.9 R* package (Hijmans et al., 2024). Symbiodiniaceae community composition was represented by relative abundances of ITS2 operational taxonomic units (OTUs). Relative abundances were Hellinger transformed to reduce the influence of highly abundant OTUs and make distances among samples more appropriate for the Euclidean-distance-based SOM analysis. Because of the large number of candidate lineages, we evaluated values up to K30 instead of K10. After preprocessing and sample matching across layers, 64 colonies were retained, representing all four major clades and 19 of the 21 genomic species hypotheses.

h. ***Microcebus* lemurs.** The eighth case study examined the cryptic radiation of Malagasy *Microcebus* mouse lemurs. van Elst et al. (2024) evaluated 27 candidate lineages derived from all 25 described species and one putative species, with *M. murinus* divided into northern and central candidates. Their integrative framework combined phylogenetic inference (*IQ-TREE* and *SVDquartets*), ancestry estimation (*NGSadmix*), an isolation-by-distance test based on genome-wide genetic distances, the genealogical divergence index, morphometric differentiation, climatic-niche overlap, reproductive activity, and acoustic communication. The study concluded that geographic population structure had frequently been interpreted as species-level divergence and proposed synonymizing seven candidates—*M. bongolavensis*, *M. boraha*, *M. ganzhorni*, *M. manitatra*, *M. marohita*, *M. mittermeieri*, and *M.* sp. 1—with their closest relatives, reducing the classification from 26 to 19 species. These decisions reflected geographic structure in genomic differentiation, evidence of gene flow, and limited differentiation in morphology, climatic niche, reproductive activity, or acoustic communication. Overall, van Elst et al. (2024) assembled RADseq-derived SNP, morphometric, geographic, bioclimatic, reproductive, and acoustic data. The morphometric data described body, head, ear, limb, foot, and tail dimensions and body mass, whereas the reproductive data recorded female reproductive condition and male testis enlargement. In our reanalysis, we included four layers: genomic, phenotypic, environmental, and spatial. The phenotypic layer combined body size and size-corrected somatic measurements with testis measurements and the inferred month of reproductive activity. The original climatic variables were replaced with a more comprehensive set of environmental variables extracted using *NicheDiv*. To expand genomic coverage (van Elst, personal communication), we reprocessed publicly available RADseq reads for the 113 individuals shared across non-genomic data. This followed the workflow of van Elst et al. (2024), including mapping to their *Microcebus murinus* Mmur 3.0 reference genome and joint SNP calling. Individual-level acoustic data were not available. After matching and filtering the four layers, 79 individuals were retained for SOM training, representing fifteen of the 27 candidate lineages. Because of the large number of candidate species, we evaluated values up to K25 instead of K10.

3. **Summary and comparison of empirical performance.** For each study, we recorded the number and proportion of retained SOM replicates supporting each value of K and the correspondence between inferred assignments and the published candidate lineages. We also examined representative SOM maps, geographic assignment maps and variable-and layer- importance summaries.

### Supplementary Figures

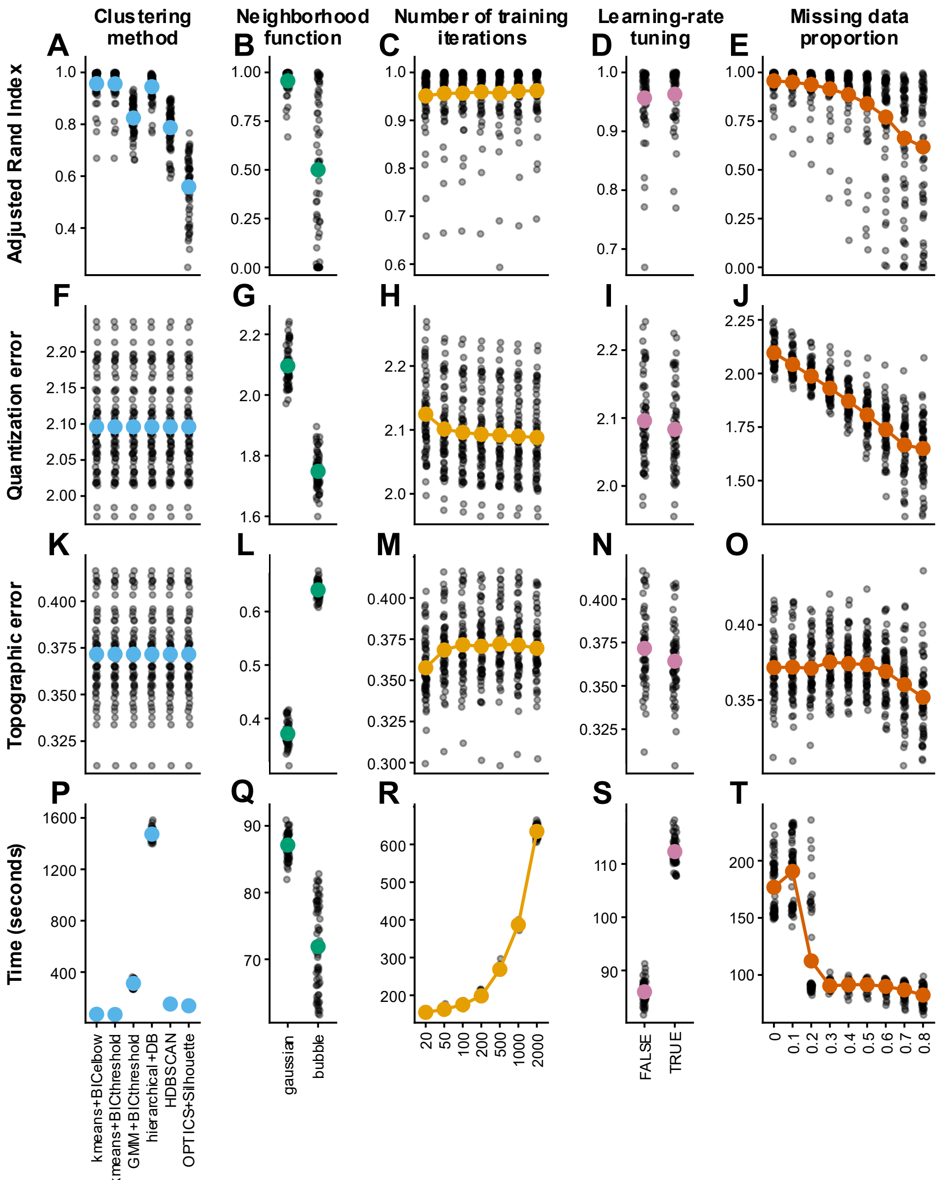


**Supplementary Figure S1: Effects of SOM parameter settings on *delimSOM* 2.0 performance in simulation set 1 with three simulated groups (K = 3).** Each simulated dataset comprised 20 individuals per group and four data layers representing SNPs, morphology, climate, and host association. Rows show Adjusted Rand Index **(A–E)**, quantization error **(F–J)**, topographic error **(K–O)**, and computation time **(P–T)**. Results are grouped by clustering and K-selection method, neighborhood function, number of training steps, learning-rate tuning, and proportion of missing data. Gray points represent simulation-level results, whereas colored points and vertical bars represent estimated marginal means ± 1 standard error from Gaussian linear mixed models with simulation as a random effect. Higher Adjusted Rand Index indicates greater agreement with the simulated group assignments, whereas lower quantization and topographic errors indicate better SOM map fit and topology preservation, respectively. Note that the decrease in quantization error with increasing missingness should not be interpreted as improved SOM performance, because distances are calculated only from available variables, reducing the information constraining sample-to-unit matching. Abbreviations: BIC = Bayesian information criterion, DB = Davies–Bouldin index, GMM = Gaussian mixture model, HDBSCAN = hierarchical density-based spatial clustering of applications with noise, OPTICS = ordering points to identify the clustering structure.

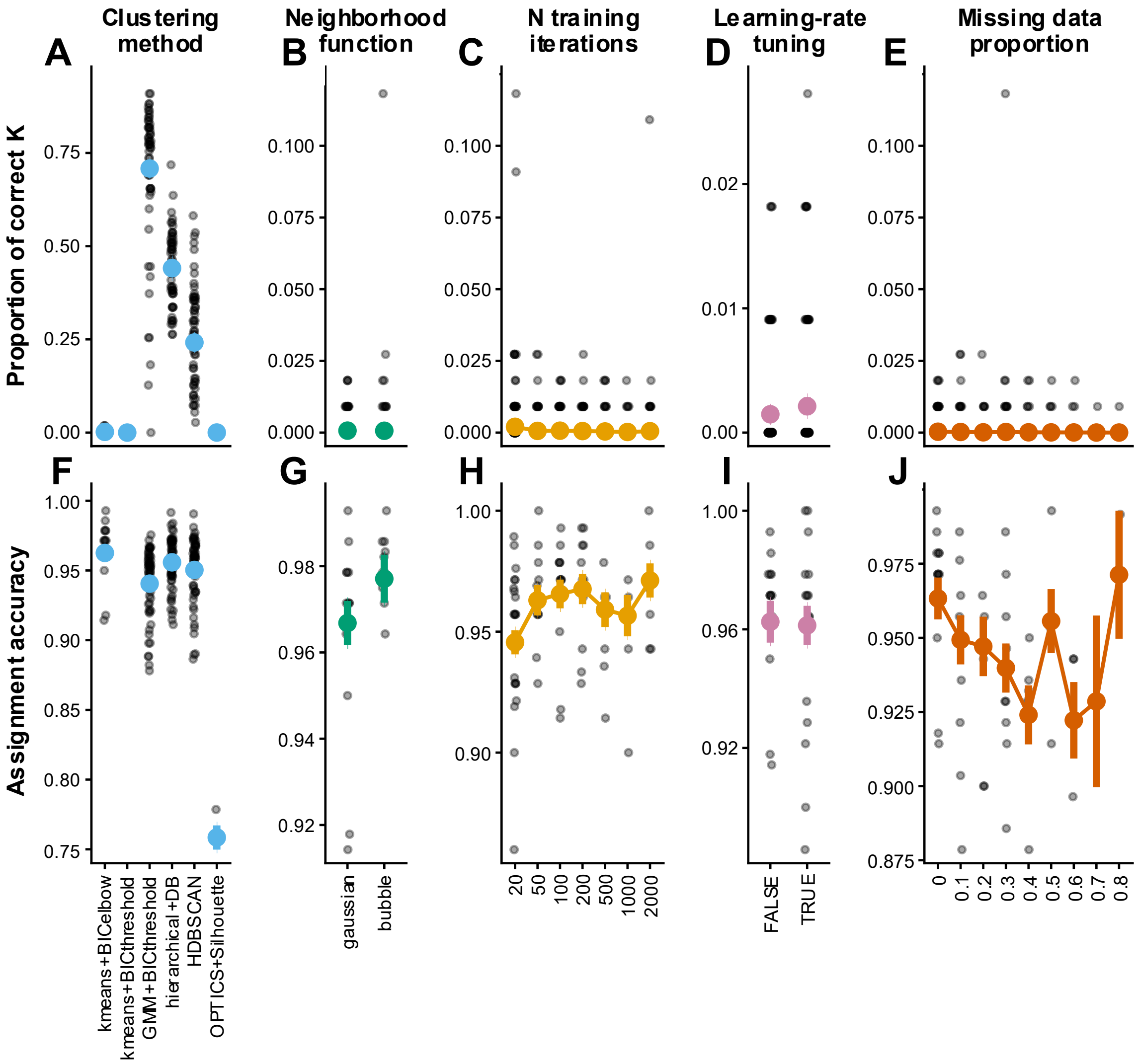


**Supplementary Figure S2: Effects of SOM parameter settings on *delimSOM* 2.0 performance in simulation set 1 with seven simulated groups (K = 7).** Each simulated dataset comprised 20 individuals per group and four data layers representing SNPs, morphology, climate, and host association. Upper panels show the proportion of simulations in which the correct number of K clusters was recovered. Lower panels show assignment accuracy (proportion of correctly assigned individuals among simulations that recovered K = 7). Results are grouped by clustering and K-selection method **(A, F)**, neighborhood function **(B, G)**, number of training steps **(C, H)**, learning-rate tuning **(D, I)**, and proportion of missing data **(E, J)**. Small gray points represent simulation-level results, whereas larger colored points and vertical bars represent estimated marginal means ± 1 standard error from binomial generalized (upper panels) and Gaussian linear mixed models (lower panels). Abbreviations: BIC = Bayesian information criterion, DB = Davies–Bouldin index, GMM = Gaussian mixture model, HDBSCAN = hierarchical density-based spatial clustering of applications with noise, OPTICS = ordering points to identify the clustering structure.

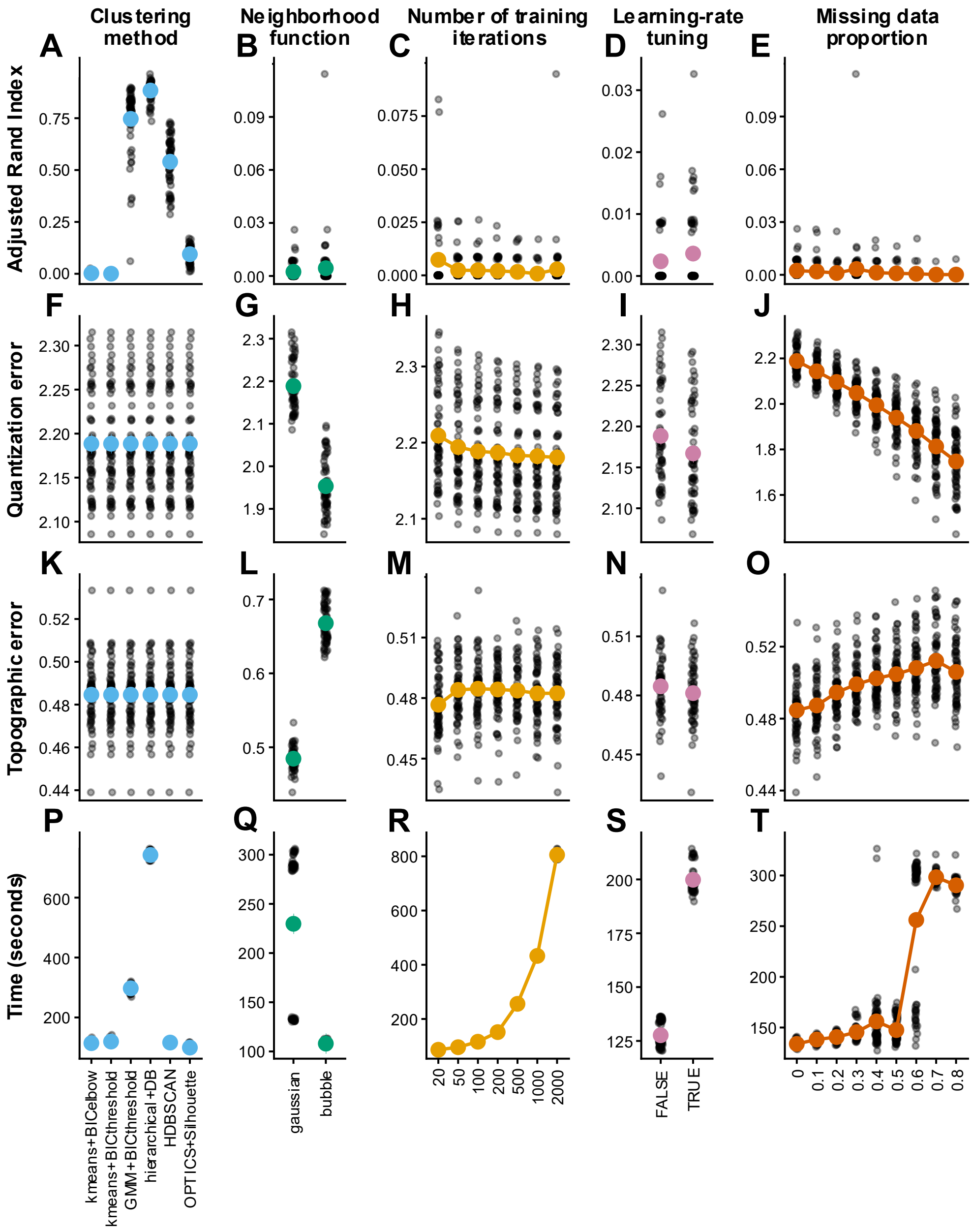


**Supplementary Figure S3: Effects of SOM parameter settings on *delimSOM* 2.0 performance in simulation set 1 with seven simulated groups (K = 7).** Each simulated dataset comprised 20 individuals per group and four data layers representing SNPs, morphology, climate, and host association. Rows show Adjusted Rand Index **(A–E)**, quantization error **(F–J)**, topographic error **(K–O)**, and computation time **(P–T)**. Results are grouped by clustering and K-selection method, neighborhood function, number of training steps, learning-rate tuning, and proportion of missing data. Gray points represent simulation-level results, whereas colored points and vertical bars represent estimated marginal means ± 1 standard error from Gaussian linear mixed models with simulation as a random effect. Higher Adjusted Rand Index indicates greater agreement with the simulated group assignments, whereas lower quantization and topographic errors indicate better SOM map fit and topology preservation, respectively. Note that the decrease in quantization error with increasing missingness should not be interpreted as improved SOM performance, because distances are calculated only from available variables, reducing the information constraining sample-to-unit matching. Abbreviations: BIC = Bayesian information criterion, DB = Davies–Bouldin index, GMM = Gaussian mixture model, HDBSCAN = hierarchical density-based spatial clustering of applications with noise, OPTICS = ordering points to identify the clustering structure.

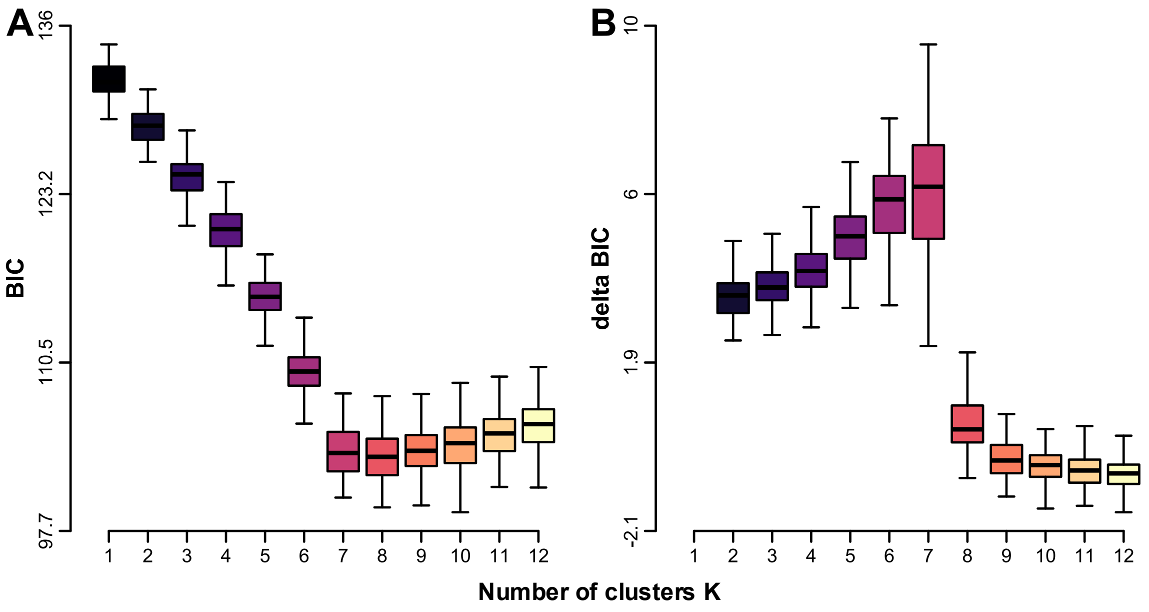


**Supplementary Figure S4: BIC (Bayesian information criterion) support for the number of clusters using k-means clustering with BIC-elbow K selection in a randomly selected simulation from simulation set 1 with seven simulated groups (K = 7).** Note the relatively small initial improvement between K = 1 and K = 2 below the BIC support threshold (*BIC.thresh = 6*), causing our conservative BIC-elbow approach to select K = 1 before evaluating subsequent improvements and thereby preventing automatic selection of the distinct elbow at K = 7. **A)** BIC values for K = 1–12 across SOM replicates. **B)** Change in BIC between successive values of K, with positive values indicating improved BIC support after adding a cluster.

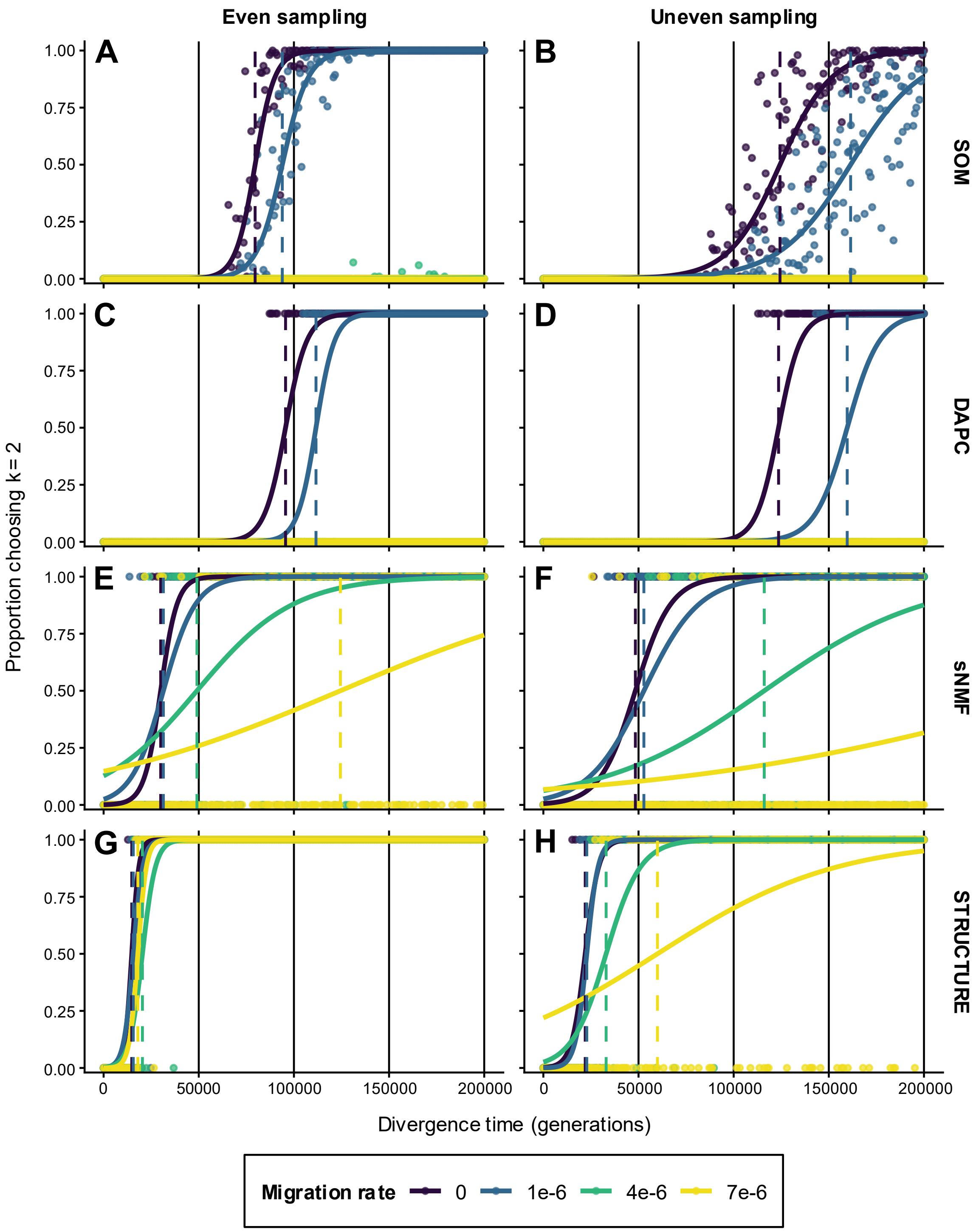


**Supplementary Figure S5: Effects of divergence time, migration rate, and uneven sampling on recovery of two simulated populations (K = 2) in simulation set 2.** Datasets comprising 1,000 SNPs were simulated with gradually increasing divergence between two populations and analyzed using *delimSOM* 2.0 **(A, B)**, *DAPC* **(C, D)**, *sNMF* **(E, F)**, and *STRUCTURE* **(G, H)**. Left and right columns show even (16 individuals per population; 1:1 ratio) and uneven sampling (24 and 8 individuals in the two populations; 3:1 ratio), respectively. Points show observed proportions selecting K = 2, solid colored lines show fitted probabilities from binomial logistic regression models, and dashed vertical lines mark the divergence time at which the fitted probability reaches 0.5. Abbreviations: *DAPC* = discriminant analysis of principal components, *sNMF* = sparse non-negative matrix factorization, SOM = self-organizing map.

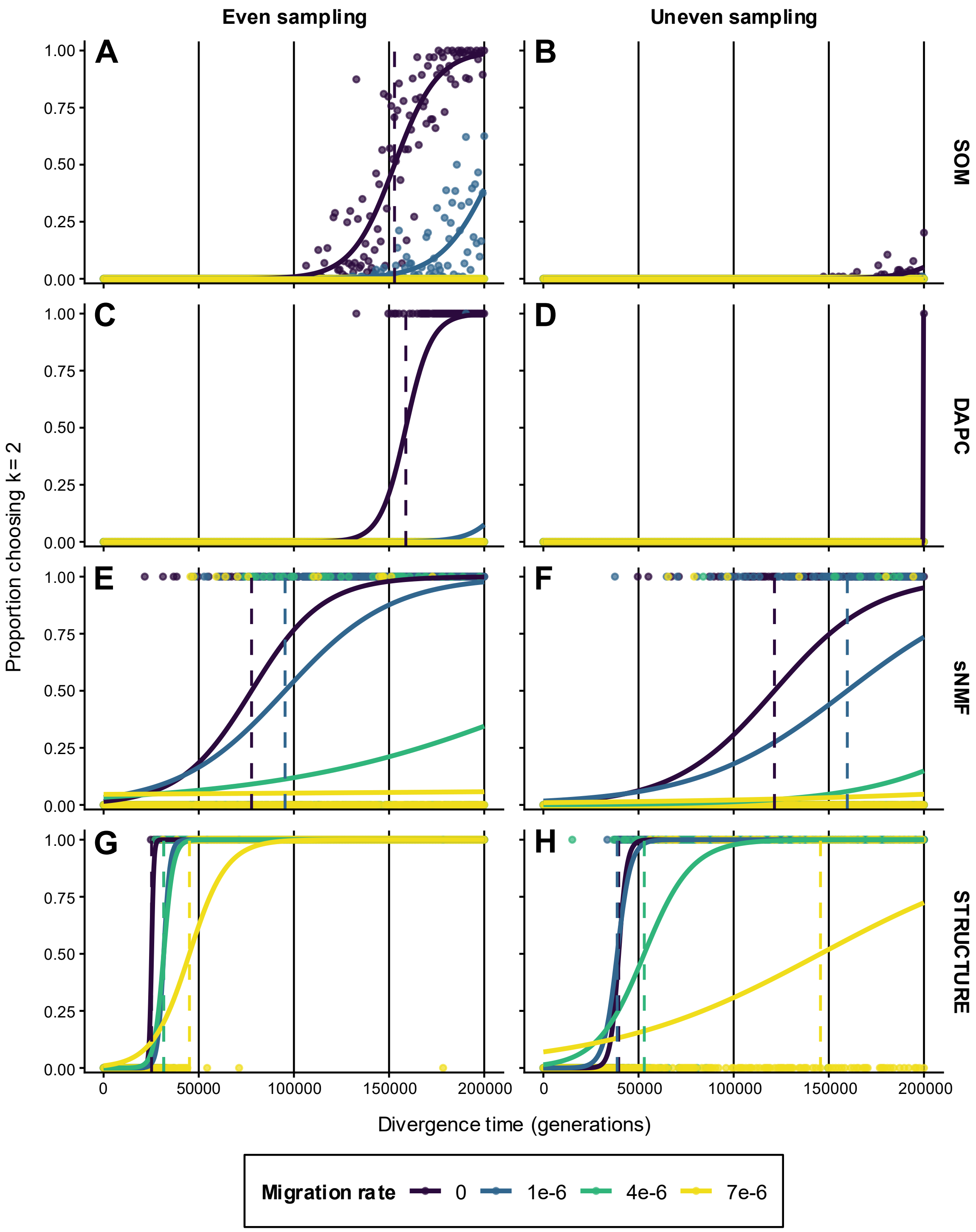


**Supplementary Figure S6: Effects of divergence time, migration rate, and uneven sampling on recovery of two simulated populations (K = 2) in simulation set 2.** Datasets comprising 1,000 SNPs were simulated with gradually increasing divergence between two populations and analyzed using *delimSOM* 2.0 **(A, B)**, *DAPC* **(C, D)**, *sNMF* **(E, F)**, and *STRUCTURE* **(G, H)**. Left and right columns show even sampling (8 individuals per population; 1:1 ratio) and uneven sampling (12 and 4 individuals in the two populations; 3:1 ratio), respectively. Points show observed proportions selecting K = 2, solid colored lines show fitted probabilities from binomial logistic regression models, and dashed vertical lines mark the divergence time at which the fitted probability reaches 0.5. Abbreviations: *DAPC* = discriminant analysis of principal components, *sNMF* = sparse non-negative matrix factorization, SOM = self-organizing map.

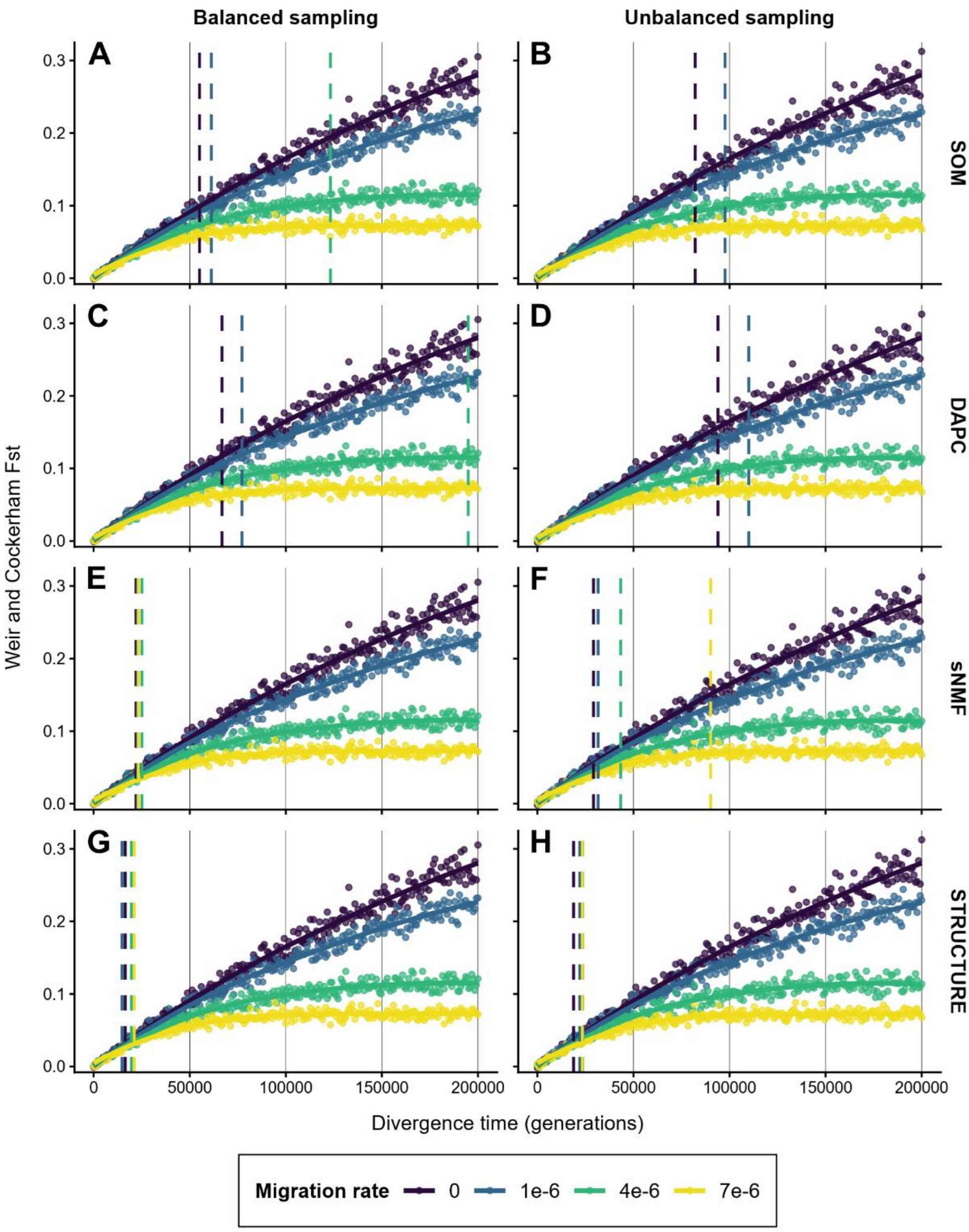


**Supplementary Figure S7**: **Genetic differentiation across divergence time and migration rates for the same simulation datasets and sampling treatments shown in Figure 2.** Datasets comprising 1,000 SNPs were simulated with gradually increasing divergence between two populations. Panels correspond to *delimSOM* 2.0 **(A, B)**, *DAPC* **(C, D)**, *sNMF* **(E, F)**, and *STRUCTURE* **(G, H)**. Left and right columns show even (24 individuals per population) and uneven sampling (36 and 12 individuals in the two populations), respectively. Points show Weir and Cockerham $F_{ST}$ values, and solid colored lines show smoothed LOESS fits generated with *ggplot2*. For direct comparison, dashed vertical lines reproduce the K = 2 recovery thresholds from Figure 2, marking the divergence time at which the fitted probability of selecting K = 2 reaches 0.5 for each method and migration rate. Colors represent migration rates. $F_{ST}$ values and LOESS curves are identical across methods within each sampling treatment, so panels differ only in the method-specific dashed threshold lines. No dashed line is shown when the fitted probability does not reach 0.5 within the simulated divergence-time range. Abbreviations: *DAPC* = discriminant analysis of principal components, *sNMF* = sparse non-negative matrix factorization, SOM = self-organizing map.

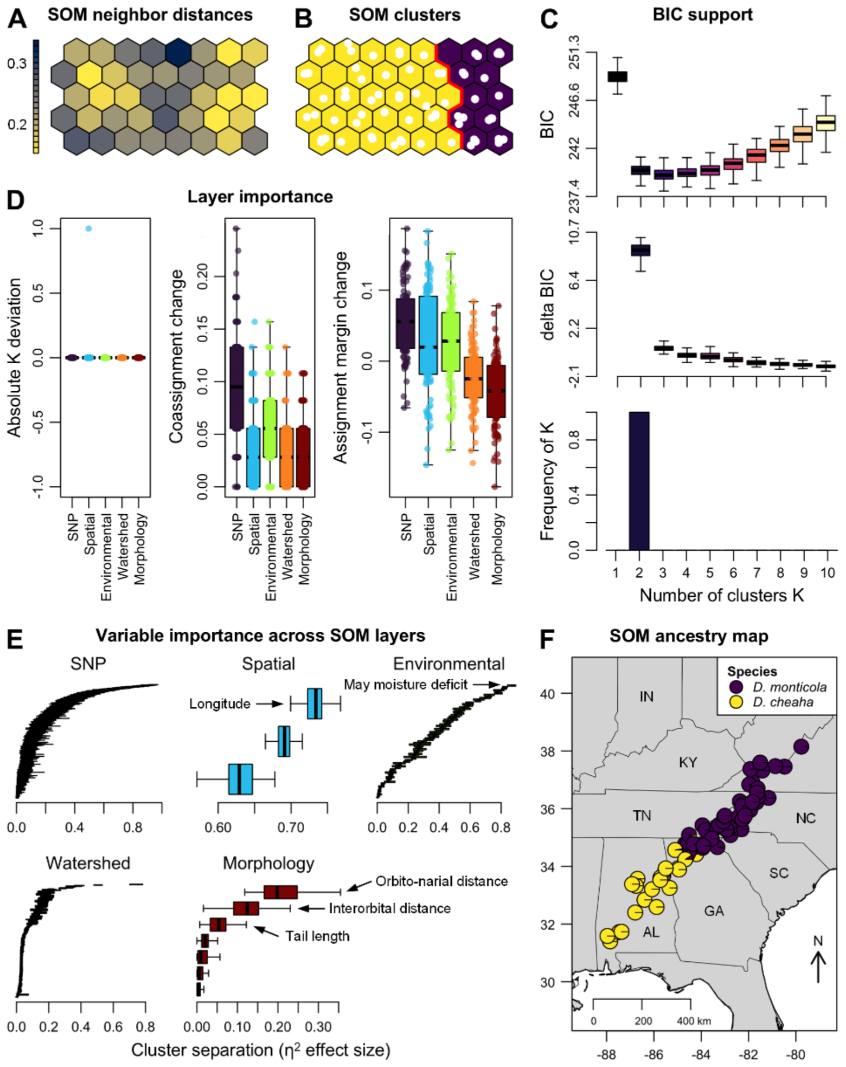


**Supplementary Figure S8**: ***delimSOM* 2.0 results for the empirical case study of 71 *Desmognathus monticola* and *D. cheaha* seal salamanders and five data layers (Pyron, 2023).** SOM (self-organizing map) analyses recovered two candidate lineages (K = 2), corresponding to the two recognized species. **A)** Neighbor distances among adjacent SOM units for a representative replicate, with dark areas showing larger distances between neighboring codebook vectors that indicate group boundaries ("ridges") and light areas showing lower distances that form "valleys," which indicate candidate lineages when separated by ridges. **B)** Recovered SOM clustering for the same replicate, with white points representing mapped individuals and red lines indicating lineage boundaries. **C)** Support for alternative numbers of K clusters based on BIC values, changes in BIC between successive K values, and the frequency with which each K was selected across SOM replicates. **D)** Layer importance from leave-one-layer-out analyses measuring changes in the inferred number of clusters (left), cluster composition (which individuals are grouped together; middle), and confidence of individual cluster assignments (how much support for the best cluster decreases relative to the second-best cluster; right) after omitting each layer, with points representing replicate-matched comparisons and boxplots summarizing variation across SOM replicates. **E)** Variable importance across SOM layers based on an ANOVA-like $\eta^2$ effect size representing the proportion of variation in each variable associated with the inferred clusters, with boxplots showing replicate-level $\eta^2$ distributions for individual variables, where higher values indicate stronger associations with cluster separation (see Supplementary Table S8 for the most important variables). **F)** Species distribution in the eastern USA, with replicate-consensus assignment coefficients showing the two inferred lineages.

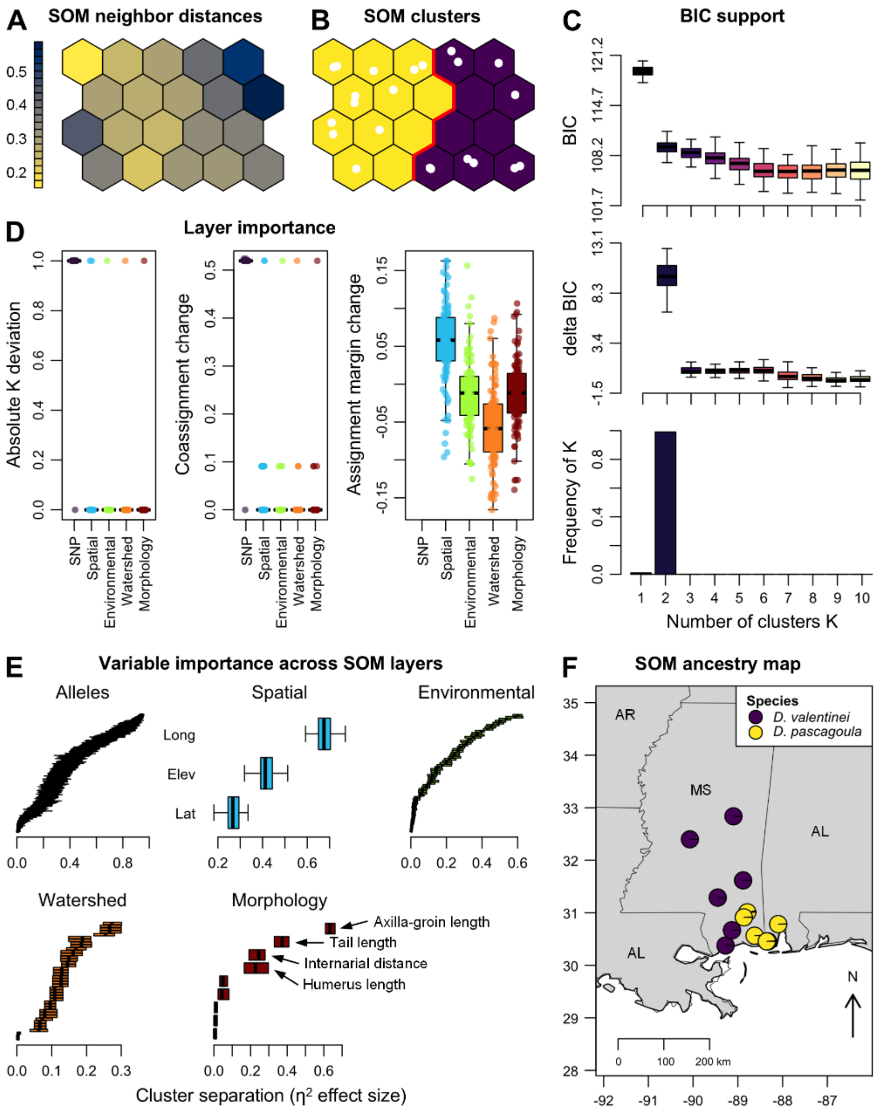


**Supplementary Figure S9: *delimSOM* 2.0 results for the empirical case study of 22 *Desmognathus valentinei* and *D. pascagoula* dusky salamanders and five data layers (Pyron et al., 2022).** SOM (self-organizing map) analyses recovered two candidate lineages (K = 2), corresponding to the two recognized species. **A)** Neighbor distances among adjacent SOM units for a representative replicate, with dark areas showing larger distances between neighboring codebook vectors that indicate group boundaries ("ridges") and light areas showing lower distances that form "valleys," which indicate candidate lineages when separated by ridges. **B)** Recovered SOM clustering for the same replicate, with white points representing mapped individuals and red lines indicating lineage boundaries**.** **C)** Support for alternative numbers of K clusters based on BIC values, changes in BIC between successive K values, and the frequency with which each K was selected across SOM replicates. **D)** Layer importance from leave-one-layer-out analyses measuring changes in the inferred number of clusters (left), cluster composition (which individuals are grouped together; middle), and confidence of individual cluster assignments (how much support for the best cluster decreases relative to the second-best cluster; right) after omitting each layer, with points representing replicate-matched comparisons and boxplots summarizing variation across SOM replicates. **E)** Variable importance across SOM layers based on an ANOVA-like $\eta^2$ effect size representing the proportion of variation in each variable associated with the inferred clusters, with boxplots showing replicate-level $\eta^2$ distributions for individual variables, where higher values indicate stronger associations with cluster separation (see Supplementary Table S9 for the most important variables). **F)** Species distributions in Mississippi (MS) and Alabama (AL), with replicate-consensus assignment coefficients showing the two inferred lineages.

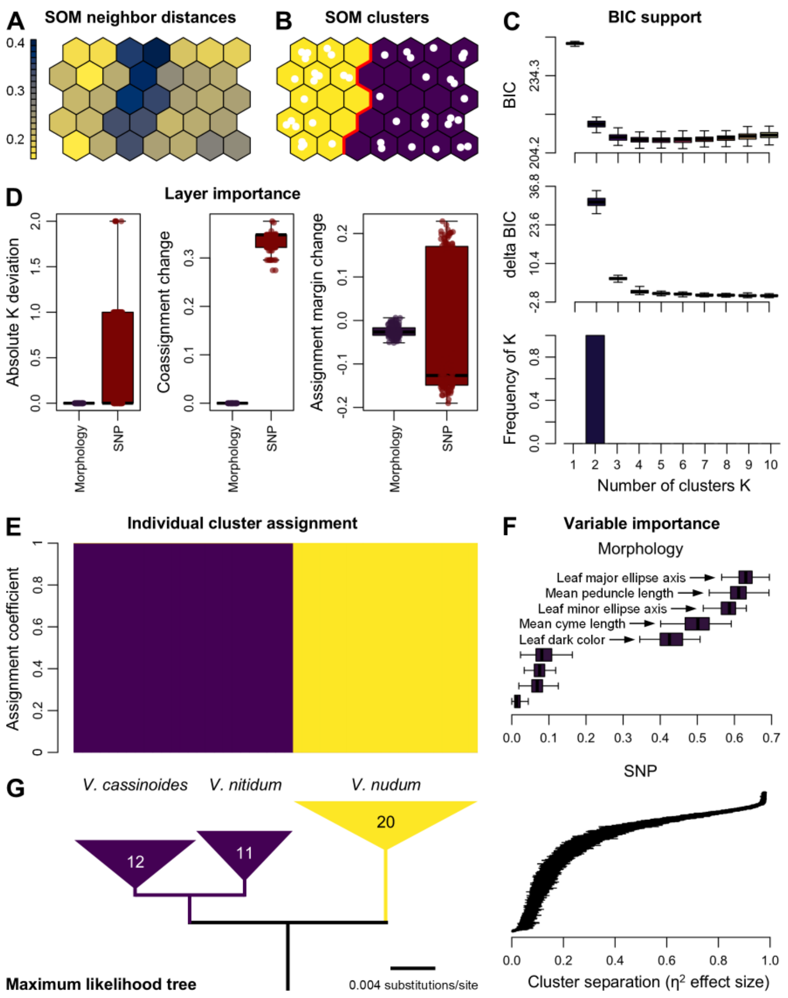


**Supplementary Figure S10: *delimSOM* 2.0 results for the empirical case study of 43 shrubs of the *Viburnum nudum* complex and two data layers (Spriggs et al., 2019).** SOM (self-organizing map) analyses recovered two candidate lineages (K = 2), with *V. cassinoides* and *V. nitidum* grouped into a single lineage. **A)** Neighbor distances among adjacent SOM units for a representative replicate, with dark areas showing larger distances between neighboring codebook vectors that indicate group boundaries ("ridges") and light areas showing lower distances that form "valleys," which indicate candidate lineages when separated by ridges. **B)** Recovered SOM clustering for the same replicate, with white points representing mapped individuals and red lines indicating lineage boundaries. **C)** Support for alternative numbers of K clusters based on BIC values, changes in BIC between successive K values, and the frequency with which each K was selected across SOM replicates. **D)** Layer importance from leave-one-layer-out analyses measuring changes in the inferred number of clusters (left), cluster composition (which individuals are grouped together; middle), and confidence of individual cluster assignments (how much support for the best cluster decreases relative to the second-best cluster; right) after omitting each layer, with points representing replicate-matched comparisons and boxplots summarizing variation across SOM replicates. **E)** *STRUCTURE*-like barplot of replicate-consensus assignment coefficients, with each bar representing one individual. **F)** Variable importance across SOM layers based on an ANOVA-like $\eta^2$ effect size representing the proportion of variation in each variable associated with the inferred clusters, with boxplots showing replicate-level $\eta^2$ distributions for individual variables, where higher values indicate stronger associations with cluster separation (see Supplementary Table S10 for the most important variables). **G)** Maximum-likelihood phylogeny of the three recognized species modified from Spriggs et al. (2019) and aligned with the STRUCTURE-like barplot in E), with colored triangles representing collapsed intraspecific branches and numbers indicating number of individuals included in our analyses (outgroup not shown).

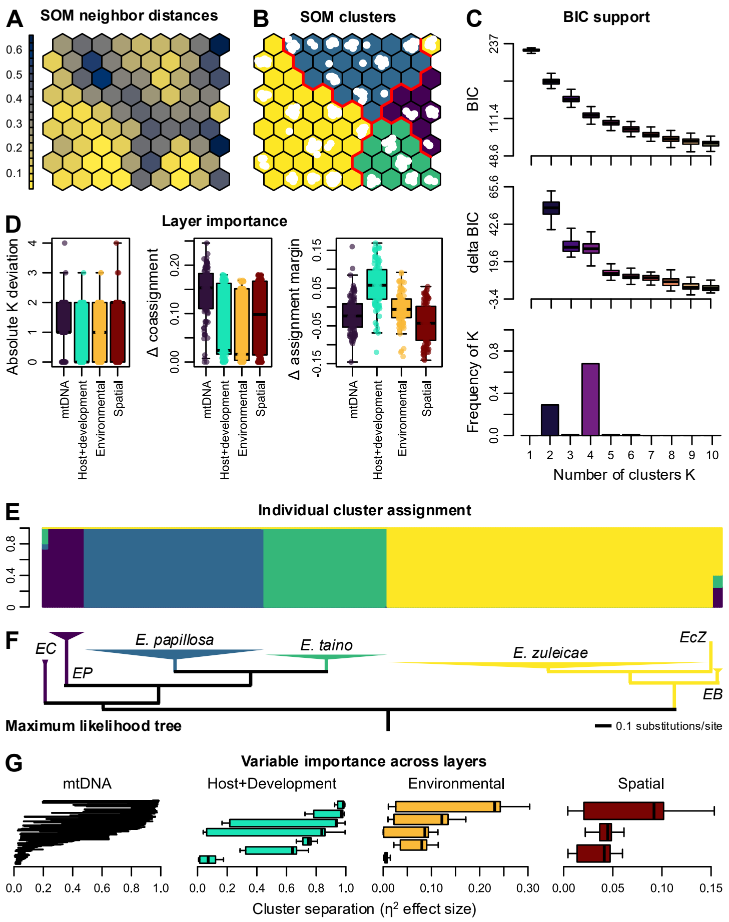


**Supplementary Figure S11: *delimSOM* 2.0 results for the empirical case study of 276 western Atlantic *Elysia* sea slugs and four data layers (Krug et al., 2026).** "mtDNA" and "Host+Development" layers represent mitochondrial COI (cytochrome c oxidase subunit I) and host use and developmental mode, respectively. SOM (self-organizing map) analyses most frequently recovered four candidate lineages (K = 4; 68% of replicates). **A)** Neighbor distances among adjacent SOM units for a representative replicate, with dark areas showing larger distances between neighboring codebook vectors that indicate group boundaries ("ridges") and light areas showing lower distances that form "valleys," which indicate candidate lineages when separated by ridges. **B)** Recovered SOM clustering for the same replicate, with white points representing mapped individuals and red lines indicating lineage boundaries. **C)** Support for alternative numbers of K clusters based on BIC values, changes in BIC between successive K values, and the frequency with which each K was selected across SOM replicates. **D)** Layer importance from leave-one-layer-out analyses measuring changes in the inferred number of clusters (left), cluster composition (which individuals are grouped together; middle), and confidence of individual cluster assignments (how much support for the best cluster decreases relative to the second-best cluster; right) after omitting each layer, with points representing replicate-matched comparisons and boxplots summarizing variation across SOM replicates. **E)** *STRUCTURE*-like barplot of replicate-consensus assignment coefficients, with each bar representing one individual. **F)** Maximum-likelihood COI phylogeny of the six described species and one candidate lineage modified from Krug et al. (2026) and aligned with the *STRUCTURE*-like barplot in E), with colored triangles representing collapsed intraspecific branches (*EC* = *E. christinae*, *EP* = *E. patina*, *EcZ* = *E.* cf. *zuleicae*, *EB* = *E. buonoi*). **G)** Variable importance across SOM layers based on an ANOVA-like $\eta^2$ effect size representing the proportion of variation in each variable associated with the inferred clusters, with boxplots showing replicate-level $\eta^2$ distributions for individual variables, where higher values indicate stronger associations with cluster separation (see Supplementary Table S11 for the most important variables).

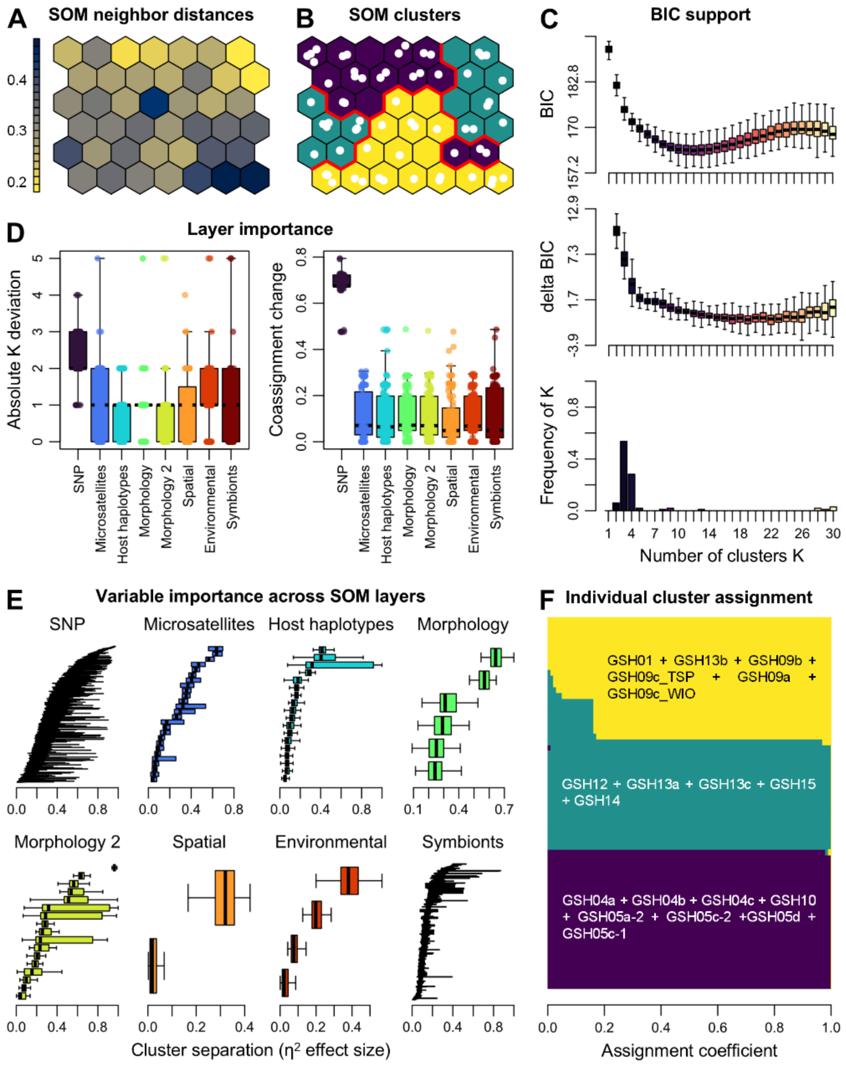


**Supplementary Figure S12: *delimSOM* 2.0 results for the empirical case study of 64 Indo-Pacific *Pocillopora* coral colonies and eight data layers (Oury et al., 2023).** The retained colonies represented 19 of their 21 genomic species hypotheses (GSHs). SOM (self-organizing map) analyses most frequently recovered three and four candidate lineages (K = 3: 54% of replicates; K = 4: 28% of replicates). **A)** Neighbor distances among adjacent SOM units for a representative replicate, with dark areas showing larger distances between neighboring codebook vectors that indicate group boundaries ("ridges") and light areas showing lower distances that form "valleys," which indicate candidate lineages when separated by ridges. **B)** Recovered SOM clustering for the same replicate, with white points representing mapped individuals and red lines indicating lineage boundaries. **C)** Support for alternative numbers of K clusters based on BIC values, changes in BIC between successive K values, and the frequency with which each K was selected across SOM replicates. **D)** Layer importance from leave-one-layer-out analyses measuring changes in the inferred number of clusters (left) and cluster composition (which individuals are grouped together; right) after omitting each layer, with points representing replicate-matched comparisons and boxplots summarizing variation across SOM replicates (the assignment-margin-change panel is not shown because differences among layers were minimal). **E)** Variable importance across SOM layers based on an ANOVA-like $\eta^2$ effect size representing the proportion of variation in each variable associated with the inferred clusters, with boxplots showing replicate-level $\eta^2$ distributions for individual variables, where higher values indicate stronger associations with cluster separation (see Supplementary Table S12 for the most important variables). **F)** *STRUCTURE*-like barplot of replicate-consensus assignment coefficients for K = 3, with each horizontal bar representing one individual and labels indicating the GSH grouped within each inferred cluster.

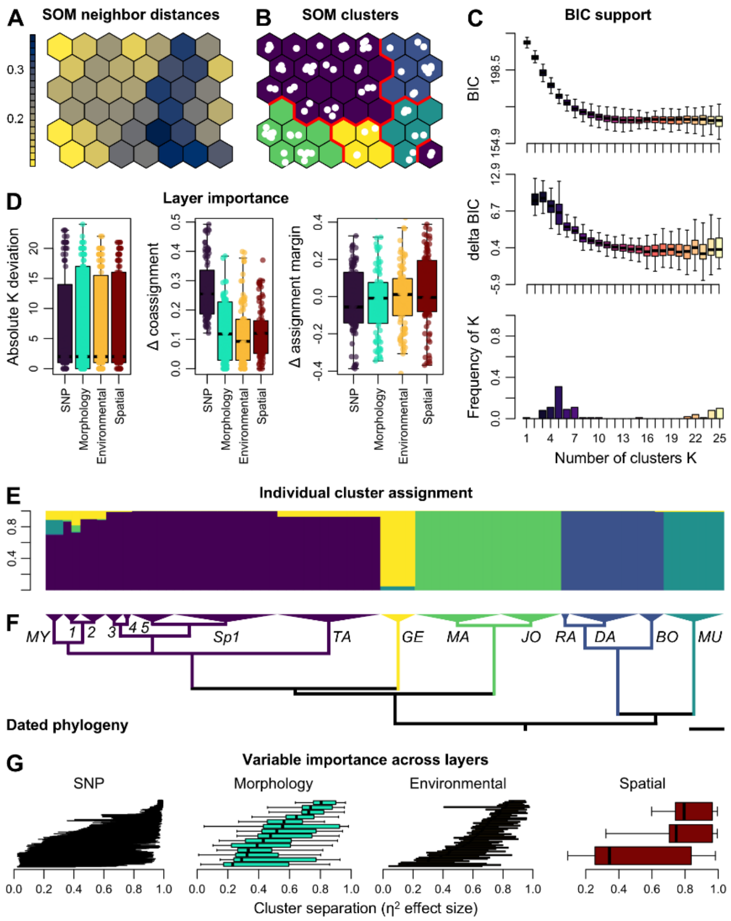


**Supplementary Figure S13: *delimSOM* 2.0 results for the empirical case study of 79 Malagasy *Microcebus* mouse lemurs and four data layers (van Elst et al., 2024).** SOM (self-organizing map) analyses most frequently recovered five candidate lineages (K = 5; 31% of replicates). **A)** Neighbor distances among adjacent SOM units for a representative replicate, with dark areas showing larger distances between neighboring codebook vectors that indicate group boundaries ("ridges") and light areas showing lower distances that form "valleys," which indicate candidate lineages when separated by ridges. **B)** Recovered SOM clustering for the same replicate, with white points representing mapped individuals and red lines indicating lineage boundaries. **C)** Support for alternative numbers of K clusters based on BIC values, changes in BIC between successive K values, and the frequency with which each K was selected across SOM replicates. **D)** Layer importance from leave-one-layer-out analyses measuring changes in the inferred number of clusters (left), cluster composition (which individuals are grouped together; middle), and confidence of individual cluster assignments (how much support for the best cluster decreases relative to the second-best cluster; right) after omitting each layer, with points representing replicate-matched comparisons and boxplots summarizing variation across SOM replicates. **E)** *STRUCTURE*-like barplot of replicate-consensus assignment coefficients for K = 5, with each bar representing one individual. **F)** Time-calibrated phylogeny modified from van Elst et al. (2024), with topology inferred by maximum likelihood from genome-wide RAD-seq SNPs and divergence times estimated under a multispecies coalescent model in BPP, aligned with the *STRUCTURE*-like barplot in E), with the scale bar representing 250 thousand years. (other *Microcebus* not shown; *MY* = *M. myoxinus*, *1* = *M. lehilahytsara*, *2* = *M. mittermeieri*, *3* = *M. mamiratra*, *4* = *M. margotmarshae*, *5* = *M. arnholdi*, Sp1 = *M.* sp. 1, *TA* = *M. tavaratra*, *GE* = *M. gerpi*, *MA* = *M. macarthurii*, *JO* = *M. jonahi*, *RA* = *M. ravelobensis*, *DA* = *M. danfossi*, *BO* = *M. bongolavensis*, *MU* = *M. murinus* north). **G)** Variable importance across SOM layers based on an ANOVA-like η² effect size representing the proportion of variation in each variable associated with the inferred clusters, with boxplots showing replicate-level η² distributions for individual variables (see Supplementary Table S13 for the most important variables).

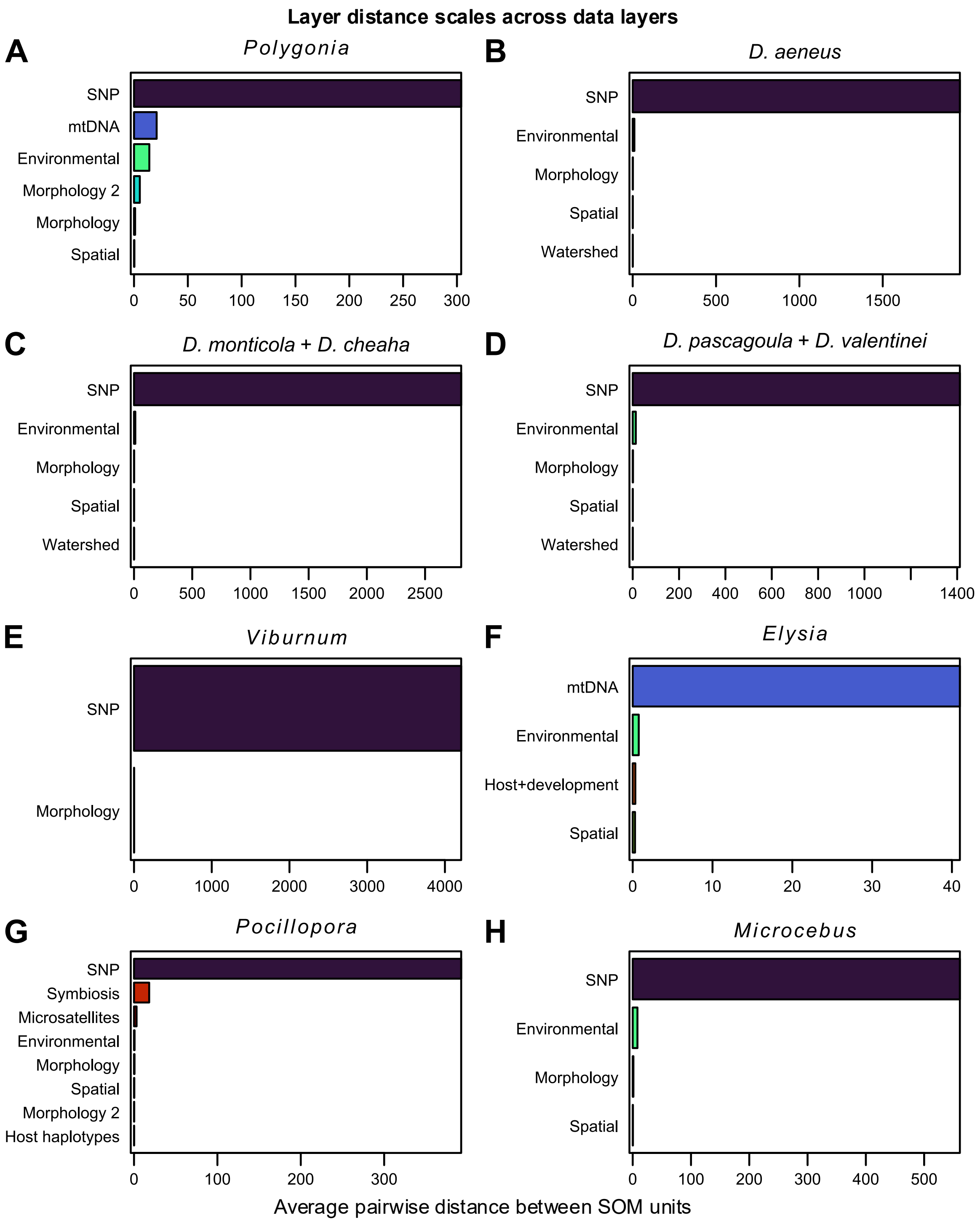


**Supplementary Figure S14: Average pairwise distance scales across data layers for the eight empirical case studies prior to SOM training, before internal SOM layer normalization is applied.** Bars show mean pairwise distances for each layer, calculated as the inverse of replicate-specific distance-normalization weights and averaged across SOM replicates. Larger values indicate larger raw within-layer distance scales (reflecting greater data dispersion, more variables, and/or differences in the layer-specific distance function), which result in greater down-weighting during internal distance normalization. Colors identify data-layer types consistently across case studies. Note that distance scales are not measures of layer importance.

### Supplementary Tables

**Supplementary Table S1:** Summary and package dependencies of *delimSOM* 2.0 functions

| Function name | Description | Package dependencies |
|---|---|---|
| *clustering.SOM* | Clusters SOM codebook vectors across replicates, filters replicates based on quantization/topographic error, selects optimal K, and builds ancestry and variable-importance summaries | base R, base R stats, base R tools, clue, cluster, clusterCrit, dbscan, kohonen, mclust |
| *make.cols.binary.SOM* | Converts specified categorical columns in dataframe into binary (0/1) indicator columns | base R |
| *plot.K.SOM* | Plots clustering-support profiles, successive ΔBIC values, and selected-K frequencies across retained SOM replicates | base R graphics, base R grDevices, viridis |
| *plot.layer.importance.leaveoneout.SOM* | Estimates and plots layer importance by rerunning replicate-matched leave-one-layer-out SOM analyses and comparing each reduced model to the full multilayer SOM | base R, base R graphics, base R grDevices, base R stats, base R tools, clue, cluster, clusterCrit, dbscan, FactoMineR, kohonen, mclust, viridis |
| *plot.layer.importance.varimp.SOM* | Summarizes and plots within-layer distributions of replicate-median eta-squared and map-variance variable-importance values | base R graphics, base R grDevices, base R stats, viridis |
| *plot.layer.distance.scale.SOM* | Plots average pairwise distance scales of input layers across SOM replicates | base R graphics, base R grDevices, viridis |
| *plot.learning.SOM* | Plots SOM learning trajectories across training steps | base R graphics, base R grDevices, viridis |
| *plot.map.SOM* | Plots replicate-consensus sample assignment coefficients as pie charts at geographic coordinates | base R graphics, base R grDevices, maps, viridis |
| *plot.model.SOM* | Plots sample mappings to SOM units, each unit's mean codebook distance to adjacent units, and boundaries between SOM-unit clusters | base R graphics, base R grDevices, kohonen, mclust, viridis |
| *plot.structure.SOM* | Plots replicate-consensus sample assignment coefficients as a Structure-like stacked barplot | base R graphics, base R grDevices, base R stats, viridis |
| *plot.variable.importance.SOM* | Plots replicate-level variable importance based on cluster-separation or map-variance metrics across SOM layers | base R graphics, base R grDevices, kohonen, matrixStats, viridis |
| *process.SNP.data.SOM* | Reads and filters genetic/genomic data from genind, genlight, VCF, FASTA, NEXUS, PHYLIP, or tabular inputs, and returns a processed SNP matrix for SOM analysis | adegenet, ape, base R utils, vcfR |
| *remove.lowCV.multicollinearity.SOM* | Removes low-variation and highly collinear variables from input data before SOM analysis | base R stats |
| *train.SOM* | Preprocesses input matrices and trains single- or multi-layer Super-SOMs | base R, base R parallel, base R stats, base R tools, doSNOW, doRNG, FactoMineR, foreach, kohonen |

*R* package versions: *adegenet v.2.1.11* (Jombart, 2008; Jombart & Ahmed, 2011), *ape v.5.8.1* (Paradis & Schliep, 2019), *base R v.4.5.1* (R Core Team, 2025), *base R graphics v.4.5.1* (R Core Team, 2025), *base R grDevices v.4.5.1* (R Core Team, 2025), *base R parallel v.4.5.1* (R Core Team, 2025), *base R stats v.4.5.1* (R Core Team, 2025), *base R tools v.4.5.1* (R Core Team, 2025), *base R utils v.4.5.1* (R Core Team, 2025), *clue v.0.6.7* (Hornik, 2005), *cluster v.2.1.8.2* (Maechler et al., 2026), *clusterCrit v.1.3* (Desgraupes, 2023), *dbscan v.1.2.4* (Hahsler & Piekenbrock, 2025), *doSNOW v.1.0.20* (Daniel et al., 2022), *doRNG v.1.8.6.3* (Gaujoux, 2026), *FactoMineR v.2.13* (Lê et al., 2008), *foreach v.1.5.2* (Microsoft, 2022), *kohonen v.3.0.13* (Wehrens & Buydens, 2007; Wehrens & Kruisselbrink, 2018), *matrixStats v.1.5* (Bengtsson, 2017), *maps v.3.4.3* (Becker et al., 2025), *mclust v.6.1.2* (Scrucca et al., 2023), *vcfR v.1.16* (Knaus & Grünwald, 2017), *viridis v.0.6.5* (Garnier et al., 2023). Abbreviation: SOM = self-organizing maps.

**Supplementary Table S2:** Summary of data layers used for analyses of eight empirical case studies

| Empirical study | Datasets (layers) | N variables before processing | N variables after processing | N individuals after processing |
|---|---|---|---|---|
| *Polygonia* anglewing butterflies from western Canada (Dupuis et al., 2018) | SNP (GBS) | 961 | 961 | 237 |
| | COI | 1,348 | 213 | 255 |
| | Morphology (continuous) | 18 | 15 | 237 |
| | Morphology (categorical) | 11 | 15 | 217 |
| | Environment | 309 | 125 | 265 |
| | Spatial | 3 | 3 | 265 |
| Seepage salamander *Desmognathus aeneus* from the southeastern United States (Pyron et al., 2024) | SNP (GBS) | 67,249 | 7,667 | 51 |
| | Morphology | 17 | 13 | 45 |
| | Environment | 310 | 53 | 56 |
| | Spatial | 3 | 3 | 56 |
| | Watershed | 7 | 98 | 56 |
| Eastern North American *Desmognathus* seal salamanders (*D. monticola* & *D. cheaha*) (Pyron, 2023) | SNP (GBS) | 56,191 | 13,031 | 71 |
| | Morphology | 16 | 7 | 71 |
| | Environment | 309 | 74 | 71 |
| | Spatial | 3 | 3 | 71 |
| | Watershed | 7 | 197 | 71 |
| *Desmognathus* dusky salamanders from Mississippi and Alabama (*D. valentinei* & *D. pascagoula*) (Pyron et al., 2022) | SNP (GBS) | 20,182 | 3,728 | 22 |
| | Morphology | 17 | 9 | 22 |
| | Environment | 309 | 59 | 22 |
| | Spatial | 3 | 3 | 22 |
| | Watershed | 7 | 42 | 22 |
| Eastern North American woody shrubs of the *Viburnum nudum* complex (Spriggs et al., 2019) | SNP (RADseq) | 58,310 | 42,159 | 65 |
| | Morphology | 10 | 9 | 145 |
| Western Atlantic sea slugs of the *Elysia papillosa* complex (Krug et al., 2026) | COI | 658 | 160 | 282 |
| | Environment | 12 | 5 | 282 |
| | Spatial | 3 | 3 | 282 |
| | Host & Development | 5 | 7 | 282 |
| Indo-Pacific *Pocillopora* corals (Oury et al., 2023) | SNP (UCE & exon loci) | 1,559 | 1,559 | 350 |
| | Microsatellites Host | 26 | 26 | 367 |
| | marker haplotypes | 2 | 31 | 367 |
| | Morphology (continuous) | 7 | 6 | 175 |
| | Morphology (binary) | 8 | 18 | 175 |
| | Spatial Environment | 2 | 2 | 367 |
| | Algal symbiont | 9 | 4 | 367 |
| | community | 552 | 500 | 213 |
| Malagasy *Microcebus* mouse lemurs (van Elst et al., 2024) | SNP (RADseq) | 4,928 | 4,664 | 102 |
| | Morphology | 20 | 18 | 102 |
| | Environment | 80 | 56 | 102 |
| | Spatial | 3 | 3 | 102 |

Abbreviations: COI = cytochrome c oxidase subunit I, GBS = genotyping-by-sequencing, N = number of, RAD-seq = restriction site-associated DNA sequencing, SNP = single-nucleotide polymorphism, UCE = ultraconserved element.

**Supplementary Table S3:** Recognized species/lineages and their sample sizes in empirical case studies

| Empirical study | Recognized species | Sample size |
|---|---|---|
| *Polygonia* anglewing butterflies from western Canada (Dupuis et al., 2018) | *Polygonia faunus* | 79 |
| | *Polygonia gracilis* | 17 |
| | *Polygonia progne* | 58 |
| | *Polygonia satyrus* | 46 |
| Seepage salamander *Desmognathus aeneus* from the southeastern United States (Pyron et al., 2024) | *Desmognathus aeneus* | 40 |
| Eastern North American *Desmognathus* seal salamanders (*D. monticola* & *D. cheaha*) (Pyron, 2023) | *Desmognathus monticola* | 49 |
| | *Desmognathus cheaha* | 22 |
| *Desmognathus* dusky salamanders from Mississippi and Alabama (*D. valentinei* & *D. pascagoula*) (Pyron et al., 2022) | *Desmognathus valentinei* | 10 |
| | *Desmognathus pascagoula* | 12 |
| Eastern North American woody shrubs of the *Viburnum nudum* complex (Spriggs et al., 2019) | *Viburnum cassinoides* | 12 |
| | *Viburnum nitidum* | 11 |
| | *Viburnum nudum* | 20 |
| Western Atlantic sea slugs of the *Elysia papillosa* complex (Krug et al., 2026) | *Elysia buonoi* | 4 |
| | *Elysia cf. zuleicae* | 1 |
| | *Elysia christinae* | 2 |
| | *Elysia papillosa* | 73 |
| | *Elysia patina* | 15 |
| | *Elysia taino* | 50 |
| | *Elysia zuleicae* | 131 |
| Indo-Pacific *Pocillopora* corals (Oury et al., 2023) | GSH01 | 7 |
| | GSH04a | 4 |
| | GSH04b | 1 |
| | GSH04c | 2 |
| | GSH05a-2 | 4 |
| | GSH05c-1 | 2 |
| | GSH05c-2 | 3 |
| | GSH05d | 3 |
| | GSH09a | 2 |
| | GSH09b | 5 |
| | GSH09c-TSP | 1 |
| | GSH09c-WIO | 3 |
| | GSH10 | 5 |
| | GSH12 | 7 |
| | GSH13a | 3 |
| | GSH13b | 3 |
| | GSH13c | 1 |
| | GSH14 | 4 |
| | GSH15 | 4 |
| Malagasy *Microcebus* mouse lemurs (van Elst et al., 2024) | *Microcebus arnholdi* | 5 |
| | *Microcebus bongolavensis* | 8 |
| | *Microcebus danfossi* | 3 |
| | *Microcebus gerpi* | 4 |
| | *Microcebus jonahi* | 7 |
| | *Microcebus lehilahytsara* | 2 |
| | *Microcebus macarthurii* | 10 |
| | *Microcebus mamiratra* | 2 |
| | *Microcebus margotmarshae* | 1 |
| | *Microcebus mittermeieri* | 3 |
| | *Microcebus murinus* (north) | 7 |
| | *Microcebus myoxinus* | 2 |
| | *Microcebus ravelobensis* | 1 |
| | *Microcebus* sp. 1 | 12 |
| | *Microcebus tavaratra* | 12 |

Sample size represents the number of individuals assigned to each recognized species or lineage among the samples included in SOM training and clustering. GSH = genomic species hypothesis.

**Supplementary Table S4:** *delimSOM* 2.0 results for eight empirical case studies

| Empirical study | N individuals for SOM training & clustering | N lineages | Clustering & K-selection method | Supported K |
|---|---|---|---|---|
| *Polygonia* anglewing butterflies from western Canada (Dupuis et al., 2018) | 200 | 4 | **kmeans + BIC elbow** | **K3** 88%, K4 12% |
| | | | kmeans + BIC threshold | K6 46%, K5 40%, K4 11% |
| | | | HDBSCAN | **K3** 84%, K4 13% |
| | | | hierarchical clustering + DB | **K3** 87%, K4 9% |
| | | | GMM + BIC threshold | K4 53%, K5 21%, K2 13%, K6 6% |
| | | | OPTICS + Silhouette | **K3** 98% |
| Seepage salamander *Desmognathus aeneus* from the southeastern United States (Pyron et al., 2024) | 40 | 1 | **kmeans + BIC elbow** | **K1** 100% |
| | | | kmeans + BIC threshold | **K1** 100% |
| | | | HDBSCAN | K2 64%, K1 23%, K3 13% |
| | | | hierarchical clustering + DB | K10 91%, K9 9% |
| | | | GMM + BIC threshold | K2 47%, K3 42%, K4 8% |
| | | | OPTICS + Silhouette | **K1** 93%, K2 7% |
| Eastern North American *Desmognathus* seal salamanders (*D. monticola* & *D. cheaha*) (Pyron, 2023) | 71 | 2 | **kmeans + BIC elbow** | **K2** 100% |
| | | | kmeans + BIC threshold | **K2** 100% |
| | | | HDBSCAN | **K2** 83%, K3 13% |
| | | | hierarchical clustering + DB | K2 74%, K10 26% |
| | | | GMM + BIC threshold | K3 53%, K2 46% |
| | | | OPTICS + Silhouette | **K1** 80%, K2 20% |
| *Desmognathus* dusky salamanders from Mississippi and Alabama (*D. valentinei* & *D. pascagoula*) (Pyron et al., 2022) | 22 | 2 | **kmeans + BIC elbow** | **K2** 99% |
| | | | kmeans + BIC threshold | **K2** 99% |
| | | | HDBSCAN | **K2** 90%, K1 5%, K3 5% |
| | | | hierarchical clustering + DB | **K10** 94%, K9 6% |
| | | | GMM + BIC threshold | K4 35%, K3 24%, K1 21%, K5 16% |
| | | | OPTICS + Silhouette | K1 52%, K2 46% |
| Eastern North American woody shrubs of the *Viburnum nudum* complex (Spriggs et al., 2019) | 46 | 3 | **kmeans + BIC elbow** | **K2** 100% |
| | | | kmeans + BIC threshold | **K2** 91%, K3 9% |
| | | | HDBSCAN | **K2** 98% |
| | | | hierarchical clustering + DB | **K2** 100% |
| | | | GMM + BIC threshold | **K3** 88%, K4 7% |
| | | | OPTICS + Silhouette | **K2** 95% |
| Western Atlantic sea slugs of the *Elysia papillosa* complex (Krug et al., 2026) | 276 | 7 | **kmeans + BIC elbow** | K4 68%, K2 29% |
| | | | kmeans + BIC threshold | K8 48%, K7 39%, K9 7% |
| | | | HDBSCAN | K4 51%, K3 23%, K5 16%, K2 5% |
| | | | hierarchical clustering + DB | K4 35%, K8 20%, K9 13%, K10 11%, K3 7% |
| | | | GMM + BIC threshold | K10 37%, K9 23%, K2 20%, K8 10% |
| | | | OPTICS + Silhouette | K4 70%, K3 25% |
| Indo-Pacific *Pocillopora* corals (Oury et al., 2023) | 64 | 19 | **kmeans + BIC elbow** | K3 54%, K4 28%, K2 26% |
| | | | kmeans + BIC threshold | K3 72%, K2 28% |
| | | | HDBSCAN | K2 34%, K3 23%, K5 14%, K4 13%, K6 11% |
| | | | hierarchical clustering + DB | **K30** 95% |
| | | | GMM + BIC threshold | K2 33%, K3 17%, K9 10%, K10 8%, K11 5% |
| | | | OPTICS + Silhouette | K2 46%, K1 46%, K3 10% |
| Malagasy *Microcebus* mouse lemurs (van Elst et al., 2024) | 79 | 15 | **kmeans + BIC elbow** | K5 31%, K4 11%, K7 11%, K25 10%, K6 9% |
| | | | kmeans + BIC threshold | K5 45%, K4 34%, K3 19% |
| | | | HDBSCAN | K2 39%, K3 23%, K4 16%, K5 16% |
| | | | hierarchical clustering + DB | K25 72%, K24 16% |
| | | | GMM + BIC threshold | K10 22%, K5 15%, K11 12%, K8 9%, K12 8% |
| | | | OPTICS + Silhouette | K2 54%, K3 17%, K1 12%, K4 9%, K5 7% |

N lineages represents the number of recognized lineages included in our analyses. Only K values with ≥5% support or the five most-supported K values are shown. Strongly supported K values (≥80% of SOM replicates) are shown in bold. Abbreviations: BIC = Bayesian information criterion, DB = Davies–Bouldin index, GMM = Gaussian mixture model, HDBSCAN = hierarchical density-based spatial clustering of applications with noise, K = number of clusters, N = number of, OPTICS = ordering points to identify the clustering structure, SOM = self-organizing map.

**Supplementary Table S5:** Summary of *delimSOM* 2.0 results for eight empirical case studies using k-means with BIC-elbow clustering and K selection, including hierarchical reanalyses

| Empirical study | N lineages | Main results | Hierarchical analyses |
|---|---|---|---|
| *Polygonia* anglewing butterflies from western Canada (Dupuis et al., 2018) | 4 | K3 90% (K4 10%): *P. faunus* and *P. satyrus* were recovered as separate lineages, while *P. gracilis* and *P. progne* were combined. The latter two species were separated in 10% of SOM replicates. No individuals showed highly intermediate cluster assignment coefficients | *P. faunus*: K1 100%<br>*P. gracilis* + *P. progne*: K2 100%[1]<br>*P. satyrus*: K1 100% |
| Seepage salamander *Desmognathus aeneus* from the southeastern United States (Pyron et al., 2024) | 1 | K1 100%: A single lineage was recovered corresponding to *D. aeneus* | NA |
| Eastern North American *Desmognathus* seal salamanders (*D. monticola* & *D. cheaha*) (Pyron, 2023) | 2 | K2 100%: *D. monticola* and *D. cheaha* were recovered as separate lineages. 1–2 individuals showed intermediate cluster assignment coefficients, consistent with potential admixture | *D. monticola*: K1 100%<br>*D. cheaha*: K1 100% |
| *Desmognathus* dusky salamanders from Mississippi and Alabama (*D. valentinei* & *D. pascagoula*) (Pyron et al., 2022) | 2 | K2 99%: *D. valentinei* and *D. pascagoula* were recovered as separate lineages. No individuals showed highly intermediate cluster assignment coefficients | *D. valentinei*: K1 100%<br>*D. pascagoula*: K1 100% |
| Eastern North American woody shrubs of the *Viburnum nudum* complex (Spriggs et al., 2019) | 3 | K2 100%: *V. nudum* was recovered as a separate lineage, while *V. cassinoides* and *V. nitidum* were combined. No individuals showed highly intermediate cluster assignment coefficients | *V. nudum*: K1 100%<br>*V. cassinoides* + *V. nitidum*: K1 100% |
| Western Atlantic sea slugs of the *Elysia papillosa* complex (Krug et al., 2026) | 7 | K4 68% (K2 29%): *E. papillosa* and *E. taino* were recovered as separate lineages, while *E. zuleicae*, *E. cf. zuleicae*, and *E. buonoi* were combined, and *E. patina* and *E. christinae* were combined. *E. christinae* showed intermediate assignment primarily between the *E. patina* and *E. taino* lineages. *E. buonoi* showed intermediate assignment but was most strongly associated with the lineage containing *E. zuleicae* and *E.* cf. *zuleicae* | *E. papillosa*: K2 88%<br>*E. zuleicae* + *E.* cf. *zuleicae* + *E. buon.*: K4 27%<br>*E. taino*: K1 91%<br>*E. patina* + *E. christinae*: K14 81% |
| Indo-Pacific *Pocillopora* corals (Oury et al., 2023) | 19 | K3 54% (K4 28%): The 19 genomic species hypotheses (GSH) were combined into three broader lineages: 1) GSH12 + GSH13a + GSH13c + GSH14 + GSH15, 2) GSH01 + GSH09a + GSH09b + GSH09c-TSP + GSH09c-WIO + GSH13b, and 3) GSH04a + GSH04b + GSH04c + GSH05a-2 + GSH05c-1 + GSH05c-2 + GSH05d + GSH10. The first corresponded to their major Clade 3, the second combined their major Clades 1 and 4, and the third corresponded to their major Clade 2. No individuals showed highly intermediate cluster assignment coefficients. | GSH04a + GSH04b + … + GSH10: K1 100%<br>GSH12 + GSH13a + … + GSH15: K1 100%<br>GSH01 + … + GSH13b: K1 72% (K2 21%, K4 6%) |
| Malagasy *Microcebus* mouse lemurs (van Elst et al., 2024) | 15 | K5 31% (K4 11%, K7 11%, K25 10%, K6 9%): The 15 candidate lineages were assigned to five inferred lineages: 1) *M. arnholdi* + *M. lehilahytsara* + *M. mamiratra* + *M. margotmarshae* + *M. mittermeieri* + *M. myoxinus* + *M.* sp. 1 + *M. tavaratra*, 2) *M. jonahi* + *M. macarthurii*, 3) *M. bongolavensis* + *M. danfossi* + *M. ravelobensis*, 4) *M. murinus* (north), and 5) *M. gerpi*. No individuals showed highly intermediate cluster assignment coefficients. | *M. ar.* + … + *M. ta.*: K3 62% (K1 26%, K4 & K5 5%)[2]<br>*M. jonahi* + *M. macarthurii*: K2 100%[3]<br>*M. bongolavensis* + … + *M. ravel.*: K1 86% (K2 14%)[4] |

Notes for hierarchical analyses: [1]*P. gracilis* and *P. progne* were recovered as separate lineages; [2]the K3 solution recovered *M. tavaratra* as a separate lineage, combined *M. arnholdi* + *M.* sp. 1, and combined *M. lehilahytsara* + *M. mamiratra* + *M. margotmarshae* + *M. mittermeieri* + *M. myoxinus*; [3]*M. jonahi* and *M. macarthurii* were recovered as separate lineages; [4]the K2 solution separated *M. danfossi* from *M. bongolavensis* + *M. ravelobensis*. Abbreviations: K = number of clusters; N = number of, SOM = self-organizing map.

**Supplementary Table S6:** Most important variables for empirical focus study 1: *Polygonia* anglewing butterflies from western Canada (Dupuis et al., 2018)

| Layer | Variables | Variable importance ($\eta^2$ cluster separation) |
|---|---|---|
| Morphology (continuous; RGB wing colors) | R15* | 0.47 |
| | G14* | 0.34 |
| | R12 | 0.19 |
| | B11 | 0.19 |
| | R11 | 0.18 |
| | G12* | 0.14 |
| | B16 | 0.14 |
| | B13 | 0.12 |
| | G11 | 0.09 |
| | B12* | 0.09 |
| | G16 | 0.08 |
| | R13 | 0.06 |
| | G13 | 0.06 |
| | B14 | 0.02 |
| | B15 | 0.01 |
| Morphology (categorical; wing characters) | Wing character 9 | 0.98 |
| | Wing character 8 state 3 | 0.92 |
| | Wing character 5 | 0.91 |
| | Wing character 1 | 0.90 |
| | Wing character 8 state 1 | 0.85 |
| | Wing character 4 | 0.77 |
| | Wing character 6* | 0.76 |
| | Wing character 2* | 0.75 |
| | Wing character 8 state 4 | 0.75 |
| | Wing character 3* | 0.61 |
| | Wing character 10* | 0.49 |
| | Wing character 8 state 2 | 0.22 |
| | Wing character 7 | 0.18 |
| | Morphotype Contrasted | 0.03 |
| | Morphotype Smeared | 0.03 |
| Environment | Burned area in May | 0.18 |
| | Topographic position index | 0.18 |
| | Burned area in June | 0.14 |
| | Mean annual relative humidity | 0.14 |
| | Herbaceous wetland cover | 0.13 |
| | Enhanced Vegetation Index in July–August | 0.12 |
| | Enhanced Vegetation Index in May–June | 0.10 |
| | Precipitation as snow in January* | 0.10 |
| | Snow water equivalent in October* | 0.10 |
| | Snow water equivalent in June | 0.10 |
| | Cropland cover | 0.09 |
| | Precipitation in January* | 0.09 |
| | Precipitation as snow in November | 0.09 |
| | Relative humidity in October* | 0.08 |
| | Hargreaves reference evapotranspiration in April | 0.08 |
| Spatial | Longitude* | 0.03 |
| | Elevation | 0.01 |
| | Latitude | 0.01 |

$\eta^2$ is an ANOVA-like effect size measuring the proportion of weighted variation in SOM codebook values explained by the inferred clusters, with higher values indicating that a variable more strongly separates the inferred lineages. Variables marked with an asterisk are in the upper quartile ($\geq 75^{th}$ percentile) of map-variance variable importance within their respective layer, indicating complementary importance beyond cluster separation. Up to the fifteen most important variables are shown for each layer. Genomic layers are not shown.

**Supplementary Table S7:** Most important variables for empirical focus study 2: seepage salamander *Desmognathus aeneus* from the southeastern United States (Pyron et al., 2024)

| Layer | Variables | Variable importance (map variance) |
|---|---|---|
| Morphology | Tail width at rear of vent | 0.03 |
| | Inter-canthal distance | 0.03 |
| | Humerus length | 0.03 |
| | Orbito-narial distance | 0.02 |
| | Eye–snout distance | 0.02 |
| | Length of third toe | 0.02 |
| | Internarial distance | 0.02 |
| | Inter-orbital distance | 0.02 |
| | Snout-vent length | 0.02 |
| | Eye diameter | 0.01 |
| | Length of third finger | 0.01 |
| | Femur length | 0.01 |
| | Tail length | 0.01 |
| Environment | Months with mean temperature > 10 °C | 0.20 |
| | Precipitation as snow in December | 0.17 |
| | Degree days < 0 °C in April | 0.16 |
| | Climatic moisture deficit in July | 0.14 |
| | Burned area in October | 0.11 |
| | Burned area in January | 0.11 |
| | Burned area in December | 0.11 |
| | May to September precipitation | 0.11 |
| | Degree days > 18 °C in January | 0.11 |
| | Burned area in November | 0.11 |
| | Frost-free days in March | 0.10 |
| | Frost-free days in February | 0.10 |
| | Burned area in February | 0.10 |
| | Climatic moisture deficit in August | 0.10 |
| | Burned area in April | 0.09 |
| Watershed & Ecoregion | HUC-6 031501 (Coosa–Tallapoosa) | 0.18 |
| | HUC-4 0315 (Alabama) | 0.18 |
| | HUC-2 06 (Tennessee) | 0.17 |
| | HUC-2 03 (South Atlantic–Gulf) | 0.17 |
| | Level IV ecoregion 66g (Southern Metasedimentary Mountains) | 0.14 |
| | Level IV ecoregion 66d (Southern Crystalline Ridges and Mountains) | 0.13 |
| | HUC-4 0602 (Middle Tennessee–Hiwassee) | 0.12 |
| | HUC-6 060200 (Middle Tennessee–Hiwassee) | 0.12 |
| | HUC-8 03150104 (Etowah) | 0.10 |
| | Level IV ecoregion 45a (Southern Inner Piedmont) | 0.06 |
| | HUC-8 06020002 (Hiwassee) | 0.06 |
| | Level IV ecoregion 65i (Fall Line Hills) | 0.06 |
| | HUC-8 06020003 (Ocoee) | 0.05 |
| | HUC-10 0602000301 (Toccoa River–Blue Ridge Lake) | 0.05 |
| | HUC-12 060200030102 (Cooper Creek) | 0.05 |
| Spatial | Latitude | 0.07 |
| | Longitude | 0.07 |
| | Elevation | 0.07 |

Variable importance is based on variance across the SOM map, with higher values indicating greater variation in variable codebook values among SOM units. Cluster-separation importance ($\eta^2$) could not be calculated because all retained SOM replicates recovered K = 1. Up to the fifteen most important variables are shown for each layer. The genomic SNP layer is not shown.

**Supplementary Table S8:** Most important variables for the empirical study of Eastern North American *Desmognathus* seal salamanders (*D. monticola* and *D. cheaha*) (Pyron, 2023)

| Layer | Variables | Variable importance ($\eta^2$ cluster separation) |
|---|---|---|
| Morphology | Orbito-narial distance | 0.20 |
| | Inter-orbital distance | 0.12 |
| | Tail length* | 0.05 |
| | Snout-vent length* | 0.02 |
| | Eye diameter | 0.01 |
| | Femur length | 0.01 |
| | Humerus length | 0.00 |
| Environment | Climatic moisture deficit in May* | 0.88 |
| | Degree days > 18 °C in February* | 0.83 |
| | Months with mean temperature > 10 °C | 0.82 |
| | Climatic moisture deficit in September* | 0.80 |
| | Climatic moisture deficit in July* | 0.78 |
| | Frost-free days in March* | 0.77 |
| | Climatic moisture deficit in October* | 0.72 |
| | Surface downwelling shortwave radiation in February | 0.72 |
| | Terrain ruggedness index | 0.70 |
| | Enhanced Vegetation Index in May–June | 0.66 |
| | Soil bulk density | 0.65 |
| | Soil organic carbon | 0.60 |
| | Climatic moisture deficit in August* | 0.59 |
| | Enhanced Vegetation Index in July–August | 0.59 |
| | Enhanced Vegetation Index in September–October | 0.56 |
| Watershed & Ecoregion | HUC-4 0315 (Alabama)* | 0.74 |
| | HUC-2 03 (South Atlantic–Gulf)* | 0.50 |
| | Level IV ecoregion 66d (Southern Crystalline Ridges and Mountains)* | 0.39 |
| | HUC-6 031501 (Coosa–Tallapoosa)* | 0.36 |
| | HUC-6 031502 (Alabama)* | 0.34 |
| | HUC-2 06 (Tennessee)* | 0.33 |
| | Level IV ecoregion 45d (Talladega Upland)* | 0.30 |
| | HUC-4 0601 (Upper Tennessee)* | 0.28 |
| | HUC-6 060101 (French Broad–Holston)* | 0.21 |
| | HUC-8 03150107 (Lower Coosa) | 0.20 |
| | Level IV ecoregion 45a (Southern Inner Piedmont)* | 0.20 |
| | Level IV ecoregion 65q (Buhrstone/Lime Hills)* | 0.20 |
| | Level IV ecoregion 65i (Fall Line Hills) | 0.18 |
| | HUC-8 03150201 (Upper Alabama) | 0.18 |
| | HUC-10 0315020107 (Jones Bluff Reservoir–Alabama) | 0.18 |
| Spatial | Longitude* | 0.73 |
| | Latitude | 0.69 |
| | Elevation | 0.63 |

$\eta^2$ is an ANOVA-like effect size measuring the proportion of weighted variation in SOM codebook values explained by the inferred clusters, with higher values indicating that a variable more strongly separates the inferred lineages. Variables marked with an asterisk are in the upper quartile (≥75th percentile) of map-variance variable importance within their respective layer, indicating complementary importance beyond cluster separation. Up to the fifteen most important variables are shown for each layer. The genomic SNP layer is not shown.

**Supplementary Table S9:** Most important variables for the empirical study of *Desmognathus* dusky salamanders from Mississippi and Alabama (*D. valentinei* and *D. pascagoula*) (Pyron et al., 2022)

| Layer | Variables | Variable importance ($\eta^2$ cluster separation) |
|---|---|---|
| Morphology | Axilla-groin length* | 0.63 |
| | Tail length | 0.37 |
| | Internarial distance* | 0.24 |
| | Humerus length | 0.23 |
| | Tail width at rear of vent | 0.05 |
| | Femur length | 0.04 |
| | Third toe length | 0.01 |
| | Snout-vent length | 0.01 |
| | Third finger length* | 0.00 |
| Environment | Climatic moisture index in November | 0.60 |
| | Burned area in November* | 0.59 |
| | Soil sand fraction | 0.54 |
| | Precipitation in October | 0.50 |
| | Soil organic carbon | 0.49 |
| | Cropland cover* | 0.46 |
| | Burned area in September* | 0.45 |
| | Climatic moisture index | 0.43 |
| | Precipitation in June | 0.40 |
| | Precipitation in March* | 0.37 |
| | Soil organic carbon density | 0.37 |
| | Burned area in October | 0.35 |
| | Climatic moisture deficit in July | 0.33 |
| | Potential evapotranspiration in wettest quarter* | 0.32 |
| | Precipitation in July | 0.31 |
| Watershed & Ecoregion | Level IV ecoregion 65d (Southern Hilly Gulf Coastal Plain) | 0.27 |
| | HUC-8 03170008 (Escatawpa)* | 0.27 |
| | HUC-10 0317000807 (Black Creek–Escatawpa River)* | 0.27 |
| | HUC-12 031700080702 (Franklin Creek)* | 0.27 |
| | HUC-8 03170009 (Mississippi Coastal)* | 0.25 |
| | HUC-10 0317000905 (Little Biloxi River–Biloxi River) | 0.19 |
| | HUC-12 031700090503 (Saucier Creek) | 0.19 |
| | HUC-10 0317000507 (Little Thompson Creek–Thompson Creek) | 0.19 |
| | HUC-12 031700050702 (Bull Branch–Thompson Creek) | 0.19 |
| | Level IV ecoregion 75i (Floodplains and Low Terraces)* | 0.17 |
| | HUC-10 0317000701 (Little Black Creek–Black Creek) | 0.16 |
| | HUC-12 031700070102 (Perkins Creek–Black Creek) | 0.16 |
| | HUC-10 0317000705 (Hickory Creek–Big Black Creek)* | 0.15 |
| | HUC-12 031700070504 (Little Sweetwater Creek–Sweetwater Creek)* | 0.15 |
| | HUC-4 0318 (Pearl) | 0.15 |
| Spatial | Longitude | 0.67 |
| | Elevation* | 0.41 |
| | Latitude | 0.27 |

$\eta^2$ is an ANOVA-like effect size measuring the proportion of weighted variation in SOM codebook values explained by the inferred clusters, with higher values indicating that a variable more strongly separates the inferred lineages. Variables marked with an asterisk are in the upper quartile (≥75th percentile) of map-variance variable importance within their respective layer, indicating complementary importance beyond cluster separation. Up to the fifteen most important variables are shown for each layer. The genomic SNP layer is not shown.

**Supplementary Table S10:** Most important variables for the empirical study of Eastern North American woody shrubs of the *Viburnum nudum* species complex (Spriggs et al., 2019)

| Layer | Variables | Variable importance ($\eta^2$ cluster separation) |
|---|---|---|
| Morphology | Leaf major ellipse axis | 0.63 |
| | Mean peduncle length* | 0.61 |
| | Leaf minor ellipse axis | 0.59 |
| | Mean cyme length | 0.50 |
| | Leaf dark coloration* | 0.42 |
| | Leaf glossy surface | 0.08 |
| | Leaf margin type* | 0.07 |
| | Petiole length | 0.07 |
| | Leaf circularity | 0.01 |

$\eta^2$ is an ANOVA-like effect size measuring the proportion of weighted variation in SOM codebook values explained by the inferred clusters, with higher values indicating that a variable more strongly separates the inferred lineages. Variables marked with an asterisk are in the upper quartile (≥75th percentile) of map-variance variable importance within their respective layer, indicating complementary importance beyond cluster separation. Up to the fifteen most important variables are shown for each layer. The genomic SNP layer is not shown.

**Supplementary Table S11:** Most important variables for the empirical study of Western Atlantic photosynthetic sea slugs of the *Elysia papillosa* species complex (Krug et al., 2026)

| Layer | Variables | Variable importance ($\eta^2$ cluster separation) |
|---|---|---|
| Environment | Mean surface chlorophyll | 0.23 |
| | Mean surface nitrate* | 0.12 |
| | Mean surface phosphate* | 0.09 |
| | Minimum surface temperature | 0.08 |
| | Mean surface silicate | 0.01 |
| Host & development | *Udotea* host* | 0.98 |
| | *Penicillus* host* | 0.97 |
| | Planktotrophic development | 0.93 |
| | *Halimeda* host | 0.84 |
| | Predominantly planktotrophic development | 0.75 |
| | Lecithotrophic development | 0.64 |
| | *Rhipilia* host | 0.07 |
| Spatial | Sea depth | 0.09 |
| | Latitude* | 0.04 |
| | Longitude | 0.04 |

$\eta^2$ is an ANOVA-like effect size measuring the proportion of weighted variation in SOM codebook values explained by the inferred clusters, with higher values indicating that a variable more strongly separates the inferred lineages. Variables marked with an asterisk are in the upper quartile (≥75th percentile) of map-variance variable importance within their respective layer, indicating complementary importance beyond cluster separation. Up to the fifteen most important variables are shown for each layer. The mitochondrial DNA layer is not shown.

**Supplementary Table S12:** Most important variables for the empirical study of Indo-Pacific *Pocillopora* corals (Oury et al., 2023)

| Layer | Variables | Variable importance ($\eta^2$ cluster separation) |
|---|---|---|
| Morphology (continuous) | Septa height* | 0.64 |
| | Maximum columella diameter 1* | 0.57 |
| | Center-to-center corallite distance | 0.31 |
| | Denticle spacing | 0.29 |
| | Maximum calice diameter 1 | 0.25 |
| | Maximum calice diameter 2 | 0.24 |
| Morphology (binary) | Rudimentary septa* | 0.96 |
| | Flat/spinulate columella* | 0.64 |
| | Acicular morphotype | 0.56 |
| | Long/thin septa* | 0.54 |
| | Styliform columella* | 0.51 |
| | Robust/spinulate septa | 0.32 |
| | Effuse morphotype | 0.29 |
| | Digitate morphotype | 0.28 |
| | Granulate surface morphotype* | 0.26 |
| | Lineate morphotype | 0.24 |
| | Oval/convex columella | 0.23 |
| | Meandroid morphotype | 0.21 |
| | Branching morphotype | 0.18 |
| Microsatellites | Pd3-004.1 | 0.64 |
| | Pd3-004.2 | 0.64 |
| | Pd3-009.1 | 0.56 |
| | Pd11.2* | 0.47 |
| | Pd3-008.2* | 0.44 |
| | Pd13.2 | 0.41 |
| | Pd3-008.1 | 0.40 |
| | Pd11.1* | 0.37 |
| | Pd3-005.2 | 0.36 |
| | PV2.2* | 0.35 |
| | Pd4.2 | 0.33 |
| | Pd3-EF35.2 | 0.32 |
| | Pd3-EF35.1 | 0.30 |
| Host marker haplotypes | ORF18* | 0.41 |
| | ORF27* | 0.40 |
| | ORF01* | 0.32 |
| | ORF34* | 0.28 |
| | ORF23 | 0.18 |
| | ORF50 | 0.16 |
| | ORF09* | 0.16 |
| | ORF30 | 0.14 |
| | ORF52 | 0.13 |
| | ORF31 | 0.12 |
| | ORF46 | 0.10 |
| | ORF17 | 0.10 |
| | ORF54 | 0.07 |
| Environmental | Current velocity | 0.38 |
| | Nitrate* | 0.20 |
| | Primary productivity | 0.08 |
| | Silicate | 0.02 |
| Spatial | Latitude | 0.32 |
| | Longitude* | 0.02 |

Variables marked with an asterisk are in the upper quartile (≥75th percentile) of map-variance variable importance within their respective layer, indicating complementary importance beyond cluster separation. Up to the thirteen most important variables are shown for each layer. The genomic SNP and symbiont layers are not shown.

**Supplementary Table S13:** Most important variables for the empirical study of Malagasy mouse lemurs of the genus *Microcebus* (van Elst et al., 2024)

| Layer | Variables | Variable importance ($\eta^2$ cluster separation) |
|---|---|---|
| Morphology | Ear length* | 0.81 |
| | Interorbital distance* | 0.74 |
| | Ear width | 0.72 |
| | Third toe length | 0.64 |
| | Hind foot length | 0.56 |
| | Tail length* | 0.55 |
| | Snout length | 0.52 |
| | Body length | 0.48 |
| | Head length | 0.43 |
| | Lower leg length | 0.39 |
| | Intraorbital distance | 0.33 |
| | Head width | 0.32 |
| | Reproductive period month* | 0.29 |
| | Tail circumference | 0.23 |
| Environment | Potential evapotranspiration in warmest quarter* | 0.87 |
| | Soil sand fraction* | 0.85 |
| | Burned area in September* | 0.83 |
| | Soil pH ($H_2O$)* | 0.83 |
| | Thornthwaite aridity index* | 0.82 |
| | Wetland cover | 0.81 |
| | Climatic moisture index* | 0.81 |
| | Enhanced Vegetation Index in April | 0.79 |
| | Enhanced Vegetation Index in March | 0.78 |
| | Soil organic carbon density* | 0.78 |
| | Minimum solar radiation | 0.78 |
| | Burned area in February | 0.77 |
| | Soil silt fraction | 0.77 |
| | Potential evapotranspiration in wettest quarter | 0.75 |
| | Maximum solar radiation* | 0.74 |
| Microsatellites | Pd3-004.2* | 0.66 |
| | Pd3-004.1 | 0.62 |
| | Pd3-009.1 | 0.58 |
| | Pd3-008.2* | 0.48 |
| | PV2.2* | 0.43 |
| | Pd13.2 | 0.43 |
| | Pd3-005.2 | 0.36 |
| | Pd3-008.1 | 0.34 |
| | Pd11.2* | 0.34 |
| | Pd11.1* | 0.29 |
| | PV2.1* | 0.28 |
| | PV7.1 | 0.26 |
| | Pd3-EF35.2 | 0.23 |
| | PV7.2 | 0.21 |
| | Pd4.2 | 0.19 |
| Spatial | Latitude | 0.79 |
| | Longitude* | 0.75 |
| | Elevation | 0.34 |

Variables marked with an asterisk are in the upper quartile (≥75th percentile) of map-variance variable importance within their respective layer, indicating complementary importance beyond cluster separation. Up to the fifteen most important variables are shown for each layer. The genomic SNP layer is not shown.

**Supplementary Table S14:** Runtime of SOM training and clustering methods for the eight empirical case studies

| Empirical study | Method | Runtime (minutes) |
|---|---|---|
| *Polygonia* anglewing butterflies from western Canada (Dupuis et al., 2018) | SOM training | 4.1 |
| | kmeans + BIC elbow | 3.0 |
| | kmeans + BIC threshold | 3.1 |
| | HDBSCAN | 2.3 |
| | hierarchical clustering + DB | 2.5 |
| | GMM + BIC threshold | 24.9 |
| | OPTICS + Silhouette | 5.9 |
| Seepage salamander *Desmognathus aeneus* from the southeastern United States (Pyron et al., 2024) | SOM training | 2.8 |
| | kmeans + BIC elbow | 3.9 |
| | kmeans + BIC threshold | 4.2 |
| | HDBSCAN | 3.3 |
| | hierarchical clustering + DB | 68.7 |
| | GMM + BIC threshold | 208.6 |
| | OPTICS + Silhouette | 1.6 |
| Eastern North American *Desmognathus* seal salamanders (*D. monticola* & *D. cheaha*) (Pyron, 2023) | SOM training | 14.7 |
| | kmeans + BIC elbow | 15.8 |
| | kmeans + BIC threshold | 17.7 |
| | HDBSCAN | 7.6 |
| | hierarchical clustering + DB | 222.0 |
| | GMM + BIC threshold | 568.7 |
| | OPTICS + Silhouette | 3.1 |
| *Desmognathus* dusky salamanders from Mississippi and Alabama (*D. valentinei* & *D. pascagoula*) (Pyron et al., 2022) | SOM training | 0.7 |
| | kmeans + BIC elbow | 2.6 |
| | kmeans + BIC threshold | 2.6 |
| | HDBSCAN | 2.0 |
| | hierarchical clustering + DB | 29.3 |
| | GMM + BIC threshold | 25.7 |
| | OPTICS + Silhouette | 1.5 |
| Eastern North American woody shrubs of the *Viburnum nudum* complex (Spriggs et al., 2019) | SOM training | 8.4 |
| | kmeans + BIC elbow | 21.7 |
| | kmeans + BIC threshold | 23.5 |
| | HDBSCAN | 10.2 |
| | hierarchical clustering + DB | 230.4 |
| | GMM + BIC threshold | 1321.2 |
| | OPTICS + Silhouette | 7.4 |
| Western Atlantic sea slugs of the *Elysia papillosa* complex (Krug et al., 2026) | SOM training | 1.7 |
| | kmeans + BIC elbow | 1.7 |
| | kmeans + BIC threshold | 1.7 |
| | HDBSCAN | 1.6 |
| | hierarchical clustering + DB | 5.5 |
| | GMM + BIC threshold | 1.9 |
| | OPTICS + Silhouette | 2.1 |
| Indo-Pacific *Pocillopora* corals (Oury et al., 2023) | SOM training | 3.0 |
| | kmeans + BIC elbow | 12.8 |
| | kmeans + BIC threshold | 4.9 |
| | HDBSCAN | 2.0 |
| | hierarchical clustering + DB | 30.0 |
| | GMM + BIC threshold | 65.8 |
| | OPTICS + Silhouette | 1.8 |
| Malagasy *Microcebus* mouse lemurs (van Elst et al., 2024) | SOM training | 2.9 |
| | kmeans + BIC elbow | 7.1 |
| | kmeans + BIC threshold | 10.1 |
| | HDBSCAN | 3.6 |
| | hierarchical clustering + DB | 83.7 |
| | GMM + BIC threshold | 311.3 |
| | OPTICS + Silhouette | 2.1 |

Runtime represents the elapsed computation time for SOM training or clustering across all SOM replicates. Abbreviations: BIC = Bayesian information criterion, DB = Davies–Bouldin index, GMM = Gaussian mixture model, HDBSCAN = hierarchical density-based spatial clustering of applications with noise, OPTICS = ordering points to identify the clustering structure, SOM = self-organizing map.

**Supplementary Table S15:** Runtime comparison between the original *delimSOM* implementation and *delimSOM* 2.0 across increasing numbers of variables

| Number of variables | Original implementation | *delimSOM* 2.0 parallel | *delimSOM* 2.0 non-parallel |
|---|---|---|---|
| 10 | 0.29 | 8.22 | 1.81 |
| 100 | 0.91 | 5.50 | 3.37 |
| 1,000 | 7.85 | 9.40 | 19.21 |
| 10,000 | 80.48 | 54.75 | 196.37 |
| 100,000 | 923.93 | 766.25 | 2,082.97 |

Runtime is reported in seconds for analyses with 50 samples, 110 replicates, 150 SOM training steps, a 4 × 3 SOM grid, and four processing cores for the parallel implementation.

#### Supplementary References


Akman, O., Comar, T., Hrozencik, D., & Gonzales, J. (2019). Data clustering and self-organizing maps in biology. In *Algebraic and Combinatorial Computational Biology* (pp. 351–374). Elsevier. https://doi.org/10.1016/B978-0-12-814066-6.00011-8

Andrews, K. R., Good, J. M., Miller, M. R., Luikart, G., & Hohenlohe, P. A. (2016). Harnessing the power of RADseq for ecological and evolutionary genomics. *Nature Reviews Genetics*, *17*(2), 81–92. https://doi.org/10.1038/nrg.2015.28

Ankerst, M., Breunig, M. M., Kriegel, H.-P., & Sander, J. (1999). OPTICS. *ACM SIGMOD Record*, *28*(2), 49–60. https://doi.org/10.1145/304181.304187

Assis, J., Tyberghein, L., Bosch, S., Verbruggen, H., Serrão, E. A., & De Clerck, O. (2018). Bio-ORACLE v2.0: Extending marine data layers for bioclimatic modelling. *Global Ecology and Biogeography*, *27*(3), 277–284. https://doi.org/10.1111/geb.12693

Bates, O. K., & Bertelsmeier, C. (2021). Climatic niche shifts in introduced species. *Current Biology*, *31*(19), R1252–R1266. https://doi.org/10.1016/j.cub.2021.08.035

Batool, F., & Hennig, C. (2021). Clustering with the average Silhouette width. *Computational Statistics & Data Analysis*, *158*, 107190. https://doi.org/10.1016/j.csda.2021.107190

Becker, R. A., Wilks, A. R., Brownrigg, R., Minka, T. P., & Deckmyn, A. (2025). *maps: Draw geographical maps* (3.4.3).

Bengtsson, H. (2017). *matrixStats: Functions that apply to rows and columns of matrices (and to vectors)*. https://github.com/HenrikBengtsson/matrixStats

Beugin, M., Gayet, T., Pontier, D., Devillard, S., & Jombart, T. (2018). A fast likelihood solution to the genetic clustering problem. *Methods in Ecology and Evolution*, *9*(4), 1006–1016. https://doi.org/10.1111/2041-210X.12968

Brentan, B., Meirelles, G., Luvizotto, E., & Izquierdo, J. (2018). Hybrid SOM+k-means clustering to improve planning, operation and management in water distribution systems. *Environmental Modelling & Software*, *106*, 77–88. https://doi.org/10.1016/j.envsoft.2018.02.013

Broennimann, O., Fitzpatrick, M. C., Pearman, P. B., Petitpierre, B., Pellissier, L., Yoccoz, N. G., Thuiller, W., Fortin, M., Randin, C., Zimmermann, N. E., Graham, C. H., & Guisan, A. (2012). Measuring ecological niche overlap from occurrence and spatial environmental data. *Global Ecology and Biogeography*, *21*(4), 481–497. https://doi.org/10.1111/j.1466-8238.2011.00698.x

Brooks, M. E., Kristensen, K., van Benthem, K. J., Magnusson, A., Berg, C. W., Nielsen, A., Skaug, H. J., Mächler, M., & Bolker, B. M. (2017). glmmTMB balances speed and flexibility among packages for zero-inflated generalized linear mixed modeling. *The R Journal*, *9*(2), 378. https://doi.org/10.32614/RJ-2017-066

Campello, R. J. G. B., Moulavi, D., & Sander, J. (2013). Density-based clustering based on hierarchical density estimates. In *Advances in Knowledge Discovery and Data Mining. PAKDD 2013. Lecture Notes in Computer Science* (Vol. 7819, pp. 160–172). Springer. https://doi.org/10.1007/978-3-642-37456-2_14

Campello, R. J. G. B., Moulavi, D., Zimek, A., & Sander, J. (2015). Hierarchical density estimates for data clustering, visualization, and outlier detection. *ACM Transactions on Knowledge Discovery from Data*, *10*(1), 1–51. https://doi.org/10.1145/2733381

Carscadden, K. A., Emery, N. C., Arnillas, C. A., Cadotte, M. W., Afkhami, M. E., Gravel, D., Livingstone, S. W., & Wiens, J. J. (2020). Niche breadth: Causes and consequences for ecology, evolution, and conservation. *The Quarterly Review of Biology*, *95*(3), 179–214. https://doi.org/10.1086/710388

Cohen, J. (1973). Eta-squared and partial eta-squared in fixed factor ANOVA designs. *Educational and Psychological Measurement*, *33*(1), 107–112. https://doi.org/10.1177/001316447303300111

Costa, I. G., Carvalho, F. de A. T. de, & Souto, M. C. P. de. (2004). Comparative analysis of clustering methods for gene expression time course data. *Genetics and Molecular Biology*, *27*(4), 623–631. https://doi.org/10.1590/S1415-47572004000400025

Cottrell, M., & Letrémy, P. (2005). Missing values: Processing with the Kohonen algorithm. *ASMDA 2005*, 489–496.

Danecek, P., Auton, A., Abecasis, G., Albers, C. A., Banks, E., DePristo, M. A., Handsaker, R. E., Lunter, G., Marth, G. T., Sherry, S. T., McVean, G., & Durbin, R. (2011). The variant call format and VCFtools. *Bioinformatics*, *27*(15), 2156–2158. https://doi.org/10.1093/bioinformatics/btr330

Daniel, F., Microsoft Corporation, & Weston, S. (2022). *doSNOW: Foreach Parallel Adaptor for the "snow" Package*. https://doi.org/10.32614/CRAN.package.doSNOW

Davies, D. L., & Bouldin, D. W. (1979). A cluster separation measure. *IEEE Transactions on Pattern Analysis and Machine Intelligence*, *PAMI-1*(2), 224–227. https://doi.org/10.1109/TPAMI.1979.4766909

Davies, E. (1997). *Machine vision: Theory, algorithms, practicalities* (2nd ed.). Academic Press.

De Kort, H., Vandepitte, K., & Honnay, O. (2013). A meta-analysis of the effects of plant traits and geographical scale on the magnitude of adaptive differentiation as measured by the difference between QST and FST. *Evolutionary Ecology*, *27*(6), 1081–1097. https://doi.org/10.1007/s10682-012-9624-9

de Souto, M. C. P., Costa, I. G., de Araujo, D. S. A., Ludermir, T. B., & Schliep, A. (2008). Clustering cancer gene expression data: A comparative study. *BMC Bioinformatics*, *9*(1), 497. https://doi.org/10.1186/1471-2105-9-497

DeRaad, D. A., McCormack, J. E., Chen, N., Peterson, A. T., & Moyle, R. G. (2022). Combining species delimitation, species trees, and tests for gene flow clarifies complex speciation in scrub-jays. *Systematic Biology*, *71*(6), 1453–1470. https://doi.org/10.1093/sysbio/syac034

Derkarabetian, S., Castillo, S., Koo, P. K., Ovchinnikov, S., & Hedin, M. (2019). A demonstration of unsupervised machine learning in species delimitation. *Molecular Phylogenetics and Evolution*, *139*, 106562. https://doi.org/10.1016/j.ympev.2019.106562

Desgraupes, B. (2023). *clusterCrit: Clustering indices* (1.3). 10.32614/CRAN.package.clusterCrit

Dormann, C. F., Elith, J., Bacher, S., Buchmann, C., Carl, G., Carré, G., Marquéz, J. R. G., Gruber, B., Lafourcade, B., Leitão, P. J., Münkemüller, T., McClean, C., Osborne, P. E., Reineking, B., Schröder, B., Skidmore, A. K., Zurell, D., & Lautenbach, S. (2013). Collinearity: A review of methods to deal with it and a simulation study evaluating their performance. *Ecography*, *36*(1), 27–46. https://doi.org/10.1111/j.1600-0587.2012.07348.x

Duan, J., & Hon, G. C. (2021). FBA: Feature barcoding analysis for single cell RNA-Seq. *Bioinformatics*, *37*(22), 4266–4268. https://doi.org/10.1093/bioinformatics/btab375

Dupuis, J. R., Peigler, R. S., Geib, S. M., & Rubinoff, D. (2018). Phylogenomics supports incongruence between ecological specialization and taxonomy in a charismatic clade of buck moths. *Molecular Ecology*, *27*(22), 4417–4429. https://doi.org/10.1111/mec.14883

Evanno, G., Regnaut, S., & Goudet, J. (2005). Detecting the number of clusters of individuals using the software STRUCTURE: A simulation study. *Molecular Ecology*, *14*(8), 2611–2620. https://doi.org/10.1111/j.1365-294X.2005.02553.x

Excoffier, L., Marchi, N., Marques, D. A., Matthey-Doret, R., Gouy, A., & Sousa, V. C. (2021). fastsimcoal2: Demographic inference under complex evolutionary scenarios. *Bioinformatics*, *37*(24), 4882–4885. https://doi.org/10.1093/bioinformatics/btab468

Falush, D., Stephens, M., & Pritchard, J. K. (2003). Inference of population structure using multilocus genotype data: Linked loci and correlated allele frequencies. *Genetics*, *164*(4), 1567–1587. https://doi.org/10.1093/genetics/164.4.1567

Feder, J. L., Opp, S. B., Wlazlo, B., Reynolds, K., Go, W., & Spisak, S. (1994). Host fidelity is an effective premating barrier between sympatric races of the apple maggot fly. *Proceedings of the National Academy of Sciences*, *91*(17), 7990–7994. https://doi.org/10.1073/pnas.91.17.7990

Flexer, A. (2001). On the use of self-organizing maps for clustering and visualization. *Intelligent Data Analysis*, *5*(5), 373–384. https://doi.org/10.3233/IDA-2001-5502

Folguera, L., Zupan, J., Cicerone, D., & Magallanes, J. F. (2015). Self-organizing maps for imputation of missing data in incomplete data matrices. *Chemometrics and Intelligent Laboratory Systems*, *143*, 146–151. https://doi.org/10.1016/j.chemolab.2015.03.002

Fraley, C., & Raftery, A. (2007). Model-based methods of classification: Using the mclust software in chemometrics. *Journal of Statistical Software*, *18*(6). https://doi.org/10.18637/jss.v018.i06

Fraley, C., & Raftery, A. E. (1998). How many clusters? Which clustering method? Answers via model-based cluster analysis. *The Computer Journal*, *41*(8), 578–588. https://doi.org/10.1093/comjnl/41.8.578

Fraley, C., & Raftery, A. E. (2002). Model-based clustering, discriminant analysis, and density estimation. *Journal of the American Statistical Association*, *97*(458), 611–631. https://doi.org/10.1198/016214502760047131

Frichot, E., & François, O. (2015). LEA: An R package for landscape and ecological association studies. *Methods in Ecology and Evolution*, *6*(8), 925–929. https://doi.org/10.1111/2041-210X.12382

Frichot, E., Mathieu, F., Trouillon, T., Bouchard, G., & François, O. (2014). Fast and efficient estimation of individual ancestry coefficients. *Genetics*, *196*(4), 973–983. https://doi.org/10.1534/genetics.113.160572

Garnier, S., Ross, N., Rudis, B., Sciaini, M., Camargo, A. P., & Scherer, C. (2023). *viridis(Lite) - Colorblind-friendly color maps for R* (0.6.4). https://github.com/sjmgarnier/viridis/

Gaujoux, R. (2026). *doRNG: Generic reproducible parallel backend for "foreach" loops* (1.8.6.3). https://CRAN.R-project.org/package=doRNG

Greenacre, M., & Primicerio, R. (2014). *Multivariate analysis of ecological data*. Fundación BBVA.

Gregorich, M., Strohmaier, S., Dunkler, D., & Heinze, G. (2021). Regression with highly correlated predictors: Variable omission is not the solution. *International Journal of Environmental Research and Public Health*, *18*(8), 4259. https://doi.org/10.3390/ijerph18084259

Guerra, L., Robles, V., Bielza, C., & Larrañaga, P. (2012). A comparison of clustering quality indices using outliers and noise. *Intelligent Data Analysis*, *16*(4), 703–715. https://doi.org/10.3233/IDA-2012-0545

Hahsler, M., & Piekenbrock, M. (2025). *dbscan: Density-based spatial clustering of applications with noise (DBSCAN) and related algorithms* (1.2.4). https://CRAN.R-project.org/package=dbscan

Hall, S. J. G. (2022). Genetic differentiation among livestock breeds—Values for Fst. *Animals*, *12*(9), 1115. https://doi.org/10.3390/ani12091115

Hamerly, G., Perelman, E., Lau, J., & Calder, B. (2005). SimPoint 3.0: Faster and more flexible program phase analysis. *Journal of Instruction-Level Parallelism*, *7*, 1–28.

Hamming, R. W. (1950). Error detecting and error correcting codes. *Bell System Technical Journal*, *29*(2), 147–160. https://doi.org/10.1002/j.1538-7305.1950.tb00463.x

Hasbún, R., González, J., Iturra, C., Fuentes, G., Alarcón, D., & Ruiz, E. (2016). Using genome-wide SNP discovery and genotyping to reveal the main source of population differentiation in Nothofagus dombeyi (Mirb.) Oerst. in Chile. *International Journal of Genomics*, *2016*, 1–10. https://doi.org/10.1155/2016/3654093

Hennig, C. (2015). What are the true clusters? *Pattern Recognition Letters*, *64*, 53–62. https://doi.org/10.1016/j.patrec.2015.04.009

Hickl, O., Queirós, P., Wilmes, P., May, P., & Heintz-Buschart, A. (2022). binny: An automated binning algorithm to recover high-quality genomes from complex metagenomic datasets. *Briefings in Bioinformatics*, *23*(6). https://doi.org/10.1093/bib/bbac431

Hijmans, R. J., Barbosa, M., Ghosh, A., & Mandel, A. (2024). *geodata: Download geographic data* (0.6-2). https://CRAN.R-project.org/package=geodata

Hornik, K. (2005). A CLUE for cluster ensembles. *Journal of Statistical Software*, *14*(12). https://doi.org/10.18637/jss.v014.i12

Hubert, L., & Arabie, P. (1985). Comparing partitions. *Journal of Classification*, *2*(1), 193–218. https://doi.org/10.1007/BF01908075

Jain, A. K. (2010). Data clustering: 50 years beyond K-means. *Pattern Recognition Letters*, *31*(8), 651–666. https://doi.org/10.1016/j.patrec.2009.09.011

Jakobsson, M., & Rosenberg, N. A. (2007). CLUMPP: A cluster matching and permutation program for dealing with label switching and multimodality in analysis of population structure. *Bioinformatics*, *23*(14), 1801–1806. https://doi.org/10.1093/bioinformatics/btm233

Johnston, H. R., Hu, Y., & Cutler, D. J. (2015). Population genetics identifies challenges in analyzing rare variants. *Genetic Epidemiology*, *39*(3), 145–148. https://doi.org/10.1002/gepi.21881

Jombart, T. (2008). adegenet: A R package for the multivariate analysis of genetic markers. *Bioinformatics*, *24*(11), 1403–1405. https://doi.org/10.1093/bioinformatics/btn129

Jombart, T. (2022). *adegenet: Tutorials*. Https://Github.Com/Thibautjombart/Adegenet/Wiki/Tutorials. https://github.com/thibautjombart/adegenet/wiki/Tutorials

Jombart, T., & Ahmed, I. (2011). adegenet 1.3-1: New tools for the analysis of genome-wide SNP data. *Bioinformatics*, *27*(21), 3070–3071. https://doi.org/10.1093/bioinformatics/btr521

Jombart, T., Devillard, S., & Balloux, F. (2010). Discriminant analysis of principal components. A new method for the analysis of genetically structured populations. *BMC Genetics*, *11*(1), 94. https://doi.org/10.1186/1471-2156-11-94

Kass, R. E., & Raftery, A. E. (1995). Bayes factors. *Journal of the American Statistical Association*, *90*(430), 773–795. https://doi.org/10.1080/01621459.1995.10476572

Kiviluoto, K. (1996). Topology preservation in self-organizing maps. *Proceedings of International Conference on Neural Networks (ICNN'96)*, 294–299. https://doi.org/10.1109/ICNN.1996.548907

Knaus, B. J., & Grünwald, N. J. (2017). vcfr: A package to manipulate and visualize variant call format data in R. *Molecular Ecology Resources*, *17*(1), 44–53. https://doi.org/10.1111/1755-0998.12549

Kohonen, T. (1990). The self-organizing map. *Proceedings of the IEEE*, *78*(9), 1464–1480. https://doi.org/10.1109/5.58325

Kohonen, T. (1995). *Self-organizing maps* (Vol. 30). Springer Berlin Heidelberg. https://doi.org/10.1007/978-3-642-97610-0

Kohonen, T. (1998). The self-organizing map. *Neurocomputing*, *21*(1–3), 1–6. https://doi.org/10.1016/S0925-2312(98)00030-7

Kohonen, T. (2001). *Self-organizing maps* (Vol. 30). Springer Berlin Heidelberg. https://doi.org/10.1007/978-3-642-56927-2

Kohonen, T. (2013). Essentials of the self-organizing map. *Neural Networks*, *37*, 52–65. https://doi.org/10.1016/j.neunet.2012.09.018

Kohonen, T. (2014). *MATLAB implementations and applications of the self-organizing map*. Unigrafia Oy.

Kopelman, N. M., Mayzel, J., Jakobsson, M., Rosenberg, N. A., & Mayrose, I. (2015). Clumpak: A program for identifying clustering modes and packaging population structure inferences across K. *Molecular Ecology Resources*, *15*(5), 1179–1191. https://doi.org/10.1111/1755-0998.12387

Krug, P. J., Trathen, D. Y., Retana, J., Rico, D. M., Bronola, J., & Ellingson, R. A. (2026). Integrating host use and dispersal ability with species delimitation to unravel a cryptic radiation of photosynthetic sea slugs. *Biological Journal of the Linnean Society*, *147*(3). https://doi.org/10.1093/biolinnean/blaf111

Kuhn, H. W. (1955). The Hungarian method for the assignment problem. *Naval Research Logistics Quarterly*, *2*(1–2), 83–97. https://doi.org/10.1002/nav.3800020109

La Torre, V., Sajina, A., Goulding, A. D., Marchesini, D., Bezanson, R., Pearl, A. N., & Sodré, L. (2024). Estimating galaxy parameters with self-organizing maps and the effect of missing data. *The Astronomical Journal*, *167*(6), 261. https://doi.org/10.3847/1538-3881/ad3821

Laurie, C. C., Doheny, K. F., Mirel, D. B., Pugh, E. W., Bierut, L. J., Bhangale, T., Boehm, F., Caporaso, N. E., Cornelis, M. C., Edenberg, H. J., Gabriel, S. B., Harris, E. L., Hu, F. B., Jacobs, K. B., Kraft, P., Landi, M. T., Lumley, T., Manolio, T. A., McHugh, C., … Weir, B. S. (2010). Quality control and quality assurance in genotypic data for genome-wide association studies. *Genetic Epidemiology*, *34*(6), 591–602. https://doi.org/10.1002/gepi.20516

Lê, S., Josse, J., & Husson, F. (2008). FactoMineR: An R package for multivariate analysis. *Journal of Statistical Software*, *25*(1). https://doi.org/10.18637/jss.v025.i01

Legendre, P., & Gallagher, E. D. (2001). Ecologically meaningful transformations for ordination of species data. *Oecologia*, *129*(2), 271–280. https://doi.org/10.1007/s004420100716

Leinonen, T., McCairns, R. J. S., O'Hara, R. B., & Merilä, J. (2013). QST–FST comparisons: Evolutionary and ecological insights from genomic heterogeneity. *Nature Reviews Genetics*, *14*(3), 179–190. https://doi.org/10.1038/nrg3395

Lek, S., Delacoste, M., Baran, P., Dimopoulos, I., Lauga, J., & Aulagnier, S. (1996). Application of neural networks to modelling nonlinear relationships in ecology. *Ecological Modelling*, *90*(1), 39–52. https://doi.org/10.1016/0304-3800(95)00142-5

Lek, S., & Guégan, J.-F. (2000). *Artificial neuronal networks*. Springer Berlin Heidelberg. https://doi.org/10.1007/978-3-642-57030-8

Lenth, R. V., & Piaskowski, J. (2025). *emmeans: Estimated marginal means, aka least-squares means* (2.0.1). https://doi.org/10.32614/CRAN.package.emmeans

Licen, S., Di Gilio, A., Palmisani, J., Petraccone, S., de Gennaro, G., & Barbieri, P. (2020). Pattern recognition and anomaly detection by self-organizing maps in a multi month e-nose survey at an industrial site. *Sensors*, *20*(7), 1887. https://doi.org/10.3390/s20071887

Liu, C., Wolter, C., Xian, W., & Jeschke, J. M. (2020). Most invasive species largely conserve their climatic niche. *Proceedings of the National Academy of Sciences*, *117*(38), 23643–23651. https://doi.org/10.1073/pnas.2004289117

Lloyd, S. (1982). Least squares quantization in PCM. *IEEE Transactions on Information Theory*, *28*(2), 129–137. https://doi.org/10.1109/TIT.1982.1056489

MacQueen, J. (1967a). Some methods for classification and analysis of multivariate observations. *Proceedings of the Fifth Berkeley Symposium on Mathematical Statistics and Probability*, *1*, 281–297.

MacQueen, J. (1967b). Some methods for classification and analysis of multivariate observations. In *Proceedings of the Fifth Berkeley Symposium on Mathematical Statistics and Probability* (Vol. 1, pp. 281–297). University of California Press.

Maechler, M., Rousseeuw, P., Struyf, A., Hubert, M., & Hornik, K. (2026). *cluster: Cluster analysis basics and extensions* (2.1.8.2). https://CRAN.R-project.org/package=cluster

Mastretta-Yanes, A., Arrigo, N., Alvarez, N., Jorgensen, T. H., Piñero, D., & Emerson, B. C. (2015). Restriction site-associated DNA sequencing, genotyping error estimation and de novo assembly optimization for population genetic inference. *Molecular Ecology Resources*, *15*(1), 28–41. https://doi.org/10.1111/1755-0998.12291

McInnes, L., Healy, J., & Astels, S. (2017). hdbscan: Hierarchical density based clustering. *The Journal of Open Source Software*, *2*(11), 205. https://doi.org/10.21105/joss.00205

Mehar, A. M., Matawie, K., & Maeder, A. (2013). Determining an optimal value of K in K-means clustering. *2013 IEEE International Conference on Bioinformatics and Biomedicine*, 51–55. https://doi.org/10.1109/BIBM.2013.6732734

Microsoft, W. S. (2022). *foreach: Provides foreach looping construct* (1.5.2). https://CRAN.R-project.org/package=foreach

Miller, A. J., Cullingham, C. I., & Peery, R. M. (2020). The influence of a priori grouping on inference of genetic clusters: Simulation study and literature review of the DAPC method. *Heredity*, *125*, 269–280. https://doi.org/https://doi.org/10.1038/s41437-020-0348-2

Milligan, G. W., & Cooper, M. C. (1985). An examination of procedures for determining the number of clusters in a data set. *Psychometrika*, *50*(2), 159–179. https://doi.org/10.1007/BF02294245

Mirkin, B. (2011). Choosing the number of clusters. *WIREs Data Mining and Knowledge Discovery*, *1*(3), 252–260. https://doi.org/10.1002/widm.15

Munkres, J. (1957). Algorithms for the assignment and transportation problems. *Journal of the Society for Industrial and Applied Mathematics*, *5*(1), 32–38. https://doi.org/10.1137/0105003

Murtagh, F., & Legendre, P. (2014). Ward's hierarchical agglomerative clustering method: Which algorithms implement Ward's criterion? *Journal of Classification*, *31*(3), 274–295. https://doi.org/10.1007/s00357-014-9161-z

Nazareno, A. G., & Knowles, L. L. (2021). There is no 'Rule of Thumb': Genomic filter settings for a small plant population to obtain unbiased gene flow estimates. *Frontiers in Plant Science*, *12*. https://doi.org/10.3389/fpls.2021.677009

Nielsen, R., Paul, J. S., Albrechtsen, A., & Song, Y. S. (2011). Genotype and SNP calling from next-generation sequencing data. *Nature Reviews Genetics*, *12*(6), 443–451. https://doi.org/10.1038/nrg2986

O'Leary, S. J., Puritz, J. B., Willis, S. C., Hollenbeck, C. M., & Portnoy, D. S. (2018). These aren't the loci you'e looking for: Principles of effective SNP filtering for molecular ecologists. *Molecular Ecology*, *27*(16), 3193–3206. https://doi.org/10.1111/mec.14792

Omran, M. G. H., Engelbrecht, A. P., & Salman, A. (2007). An overview of clustering methods. *Intelligent Data Analysis*, *11*(6), 583–605. https://doi.org/10.3233/IDA-2007-11602

Opedal, Ø. H., Armbruster, W. S., Hansen, T. F., Holstad, A., Pélabon, C., Andersson, S., Campbell, D. R., Caruso, C. M., Delph, L. F., Eckert, C. G., Lankinen, Å., Walter, G. M., Ågren, J., & Bolstad, G. H. (2023). Evolvability and trait function predict phenotypic divergence of plant populations. *Proceedings of the National Academy of Sciences*, *120*(1). https://doi.org/10.1073/pnas.2203228120

Oury, N., Noël, C., Mona, S., Aurelle, D., & Magalon, H. (2023). From genomics to integrative species delimitation? The case study of the Indo-Pacific Pocillopora corals. *Molecular Phylogenetics and Evolution*, *184*, 107803. https://doi.org/10.1016/j.ympev.2023.107803

Paradis, E., & Schliep, K. (2019). ape 5.0: An environment for modern phylogenetics and evolutionary analyses in R. *Bioinformatics*, *35*, 526–528. https://doi.org/10.1093/bioinformatics/bty633

Park, S.-H., Hosoishi, S., Ogata, K., & Kuboki, Y. (2014). Clustering of ant communities and indicator species analysis using self-organizing maps. *Comptes Rendus. Biologies*, *337*(9), 545–552. https://doi.org/10.1016/j.crvi.2014.07.003

Park, Y.-S., Tison, J., Lek, S., Giraudel, J.-L., Coste, M., & Delmas, F. (2006). Application of a self-organizing map to select representative species in multivariate analysis: A case study determining diatom distribution patterns across France. *Ecological Informatics*, *1*(3), 247–257. https://doi.org/10.1016/j.ecoinf.2006.03.005

Pearson, K. (1896). VII. Mathematical contributions to the theory of evolution.—III. Regression, heredity, and panmixia. *Philosophical Transactions of the Royal Society of London. Series A, Containing Papers of a Mathematical or Physical Character*, *187*, 253–318. https://doi.org/10.1098/rsta.1896.0007

Pfeifer, S. P. (2017). From next-generation resequencing reads to a high-quality variant data set. *Heredity*, *118*(2), 111–124. https://doi.org/10.1038/hdy.2016.102

Pinheiro, J., & Bates, D. (2006). *Mixed-effects models in S and S-PLUS*. Springer Science & Business Media.

Pinheiro, J., Bates, D., & R Core Team. (2025). *nlme: Linear and nonlinear mixed effects models* (3.1-168). https://doi.org/10.32614/CRAN.package.nlme

Pompanon, F., Bonin, A., Bellemain, E., & Taberlet, P. (2005). Genotyping errors: Causes, consequences and solutions. *Nature Reviews Genetics*, *6*(11), 847–859. https://doi.org/10.1038/nrg1707

Pritchard, J. K., Stephens, M., & Donnelly, P. (2000). Inference of population structure using multilocus genotype data. *Genetics*, *155*(2), 945–959. https://doi.org/10.1093/genetics/155.2.945

Pyron, R. A. (2023). Unsupervised machine learning for species delimitation, integrative taxonomy, and biodiversity conservation. *Molecular Phylogenetics and Evolution*, *189*, 107939. https://doi.org/10.1016/j.ympev.2023.107939

Pyron, R. A., Kakkera, A., Beamer, D. A., & O'Connell, K. A. (2024). Discerning structure versus speciation in phylogeographic analysis of seepage salamanders (Desmognathus aeneus) using demography, environment, geography, and phenotype. *Molecular Ecology*, *33*(2). https://doi.org/10.1111/mec.17219

Pyron, R. A., O'Connel, K. A., Lamb, J. Y., & Beamer, D. A. (2022). A new, narrowly endemic species of swamp-dwelling dusky salamander (Plethodontidae: Desmognathus) from the Gulf Coastal Plain of Mississippi and Alabama. *Zootaxa*, *5133*(1), 53–82. https://doi.org/10.11646/zootaxa.5133.1.3

Pyron, R. A., O'Connell, K. A., Duncan, S. C., Burbrink, F. T., & Beamer, D. A. (2023). Speciation hypotheses from phylogeographic delimitation yield an integrative taxonomy for seal salamanders (Desmognathus monticola). *Systematic Biology*, *72*(1), 179–197. https://doi.org/10.1093/sysbio/syac065

R Core Team. (2025). *R: A language and environment for statistical computing* (4.4.1). R Foundation for Statistical Computing. https://www.r-project.org/.

Raivio, K., Simula, O., & Laiho, J. (2001). Neural analysis of mobile radio access network. *Proceedings 2001 IEEE International Conference on Data Mining*, 457–464. https://doi.org/10.1109/ICDM.2001.989552

Ramírez-Martínez, M. M., & Tlapaya-Romero, L. (2023). Association of bat flies (Diptera: Streblidae) and bats: Richness and host specificity in Western Mexico. *International Journal for Parasitology: Parasites and Wildlife*, *21*, 160–167. https://doi.org/10.1016/j.ijppaw.2023.05.001

Rejeb, S., Duveau, C., & Rebafka, T. (2022). Self-organizing maps for exploration of partially observed data and imputation of missing values. *Chemometrics and Intelligent Laboratory Systems*, *231*, 104653. https://doi.org/10.1016/j.chemolab.2022.104653

Richardson, J. T. E. (2011). Eta squared and partial eta squared as measures of effect size in educational research. *Educational Research Review*, *6*(2), 135–147. https://doi.org/10.1016/j.edurev.2010.12.001

Rodriguez, M. Z., Comin, C. H., Casanova, D., Bruno, O. M., Amancio, D. R., Costa, L. da F., & Rodrigues, F. A. (2019). Clustering algorithms: A comparative approach. *PLOS ONE*, *14*(1), e0210236. https://doi.org/10.1371/journal.pone.0210236

Rodríguez-Echeverría, J., Semanjski, I., Van Gheluwe, C., Ochoa, D., IJben, H., & Gautama, S. (2020). Density-based spatial clustering and ordering points approach for characterizations of tourist behaviour. *ISPRS International Journal of Geo-Information*, *9*(11), 686. https://doi.org/10.3390/ijgi9110686

Rousseeuw, P. J. (1987). Silhouettes: A graphical aid to the interpretation and validation of cluster analysis. *Journal of Computational and Applied Mathematics*, *20*, 53–65. https://doi.org/10.1016/0377-0427(87)90125-7

Samad, T., & Harp, S. (1992). Self–organization with partial data. *Network: Computation in Neural Systems*, *3*(2), 205–212. https://doi.org/10.1088/0954-898X/3/2/008

Schönberger, D., MacDonald, Z. G., Schmidt, B. C., & Dupuis, J. R. (2026). NicheDiv: A DAPC framework to quantify niche divergence across highly multivariate environmental space. *BioRxiv*. https://doi.org/10.64898/2026.06.19.733388

Scrucca, L., Fop, M., Murphy, T. B., & Raftery, A. E. (2016). mclust 5: clustering, classification and density estimation using Gaussian finite mixture models. *R Journal*, *8*(1), 289–317.

Scrucca, L., Fraley, C., Murphy, T. B., & Adrian E., R. (2023). *Model-based clustering, classification, and density estimation using mclust in R*. Chapman and Hall/CRC. https://doi.org/10.1201/9781003277965

Siefert, A., Violle, C., Chalmandrier, L., Albert, C. H., Taudiere, A., Fajardo, A., Aarssen, L. W., Baraloto, C., Carlucci, M. B., Cianciaruso, M. V., de L. Dantas, V., de Bello, F., Duarte, L. D. S., Fonseca, C. R., Freschet, G. T., Gaucherand, S., Gross, N., Hikosaka, K., Jackson, B., … Wardle, D. A. (2015). A global meta-analysis of the relative extent of intraspecific trait variation in plant communities. *Ecology Letters*, *18*(12), 1406–1419. https://doi.org/10.1111/ele.12508

Smith, M. L., & Carstens, B. C. (2020). Process-based species delimitation leads to identification of more biologically relevant species. *Evolution*, *74*(2), 216–229. https://doi.org/10.1111/evo.13878

Smith, M. L., & Hahn, M. W. (2024). Selection leads to false inferences of introgression using popular methods. *Genetics*, *227*(4). https://doi.org/10.1093/genetics/iyae089

Sokal, R. R., & Rohlf, F. J. (2012). *Biometry: The principles and practice of statistics in biological research* (4th ed.). W. H. Freeman.

Spearman, C. (1904). The proof and measurement of association between two things. *The American Journal of Psychology*, *15*(1), 72. https://doi.org/10.2307/1412159

Spriggs, E. L., Eaton, D. A. R., Sweeney, P. W., Schlutius, C., Edwards, E. J., & Donoghue, M. J. (2019). Restriction-site-associated DNA sequencing reveals a cryptic Viburnum species on the North American coastal plain. *Systematic Biology*, *68*(2), 187–203. https://doi.org/10.1093/sysbio/syy084

Steinley, Douglas. (2006). K-means clustering: A half-century synthesis. *British Journal of Mathematical and Statistical Psychology*, *59*(1), 1–34. https://doi.org/10.1348/000711005X48266

Sun, Y. (2000). On quantization error of self-organizing map network. *Neurocomputing*, *34*(1–4), 169–193. https://doi.org/10.1016/S0925-2312(00)00292-7

Thorndike, R. L. (1953). Who belongs in the family? *Psychometrika*, *18*(4), 267–276. https://doi.org/10.1007/BF02289263

Tibshirani, R., Walther, G., & Hastie, T. (2001). Estimating the number of clusters in a data set via the Gap statistic. *Journal of the Royal Statistical Society Series B: Statistical Methodology*, *63*(2), 411–423. https://doi.org/10.1111/1467-9868.00293

Ultsch, A. (1993). Self-organizing neural networks for visualisation and classification. In *Information and classification. Studies in classification, data analysis and knowledge organization* (pp. 307–313). Springer. https://doi.org/10.1007/978-3-642-50974-2_31

Valente, F., & Wellekens, C. (2004). Scoring unknown speaker clustering: VB vs. BIC. *Interspeech 2004*, 593–596. https://doi.org/10.21437/Interspeech.2004-248

van Elst, T., Sgarlata, G. M., Schüßler, D., Tiley, G. P., Poelstra, J. W., Scheumann, M., Blanco, M. B., Aleixo-Pais, I. G., Rina Evasoa, M., Ganzhorn, J. U., Goodman, S. M., Hasiniaina, A. F., Hending, D., Hohenlohe, P. A., Ibouroi, M. T., Iribar, A., Jan, F., Kappeler, P. M., Le Pors, B., … Salmona, J. (2024). Integrative taxonomy clarifies the evolution of a cryptic primate clade. *Nature Ecology & Evolution*. https://doi.org/10.1038/s41559-024-02547-w

Vatcheva, K. P., & Lee, M. (2016). Multicollinearity in regression analyses conducted in epidemiologic studies. *Epidemiology: Open Access*, *06*(02). https://doi.org/10.4172/2161-1165.1000227

Vesanto, J. (1999). SOM-based data visualization methods. *Intelligent Data Analysis*, *3*(2), 111–126.

Vesanto, J., & Alhoniemi, E. (2000). Clustering of the self-organizing map. *IEEE Transactions on Neural Networks*, *11*(3), 586–600. https://doi.org/10.1109/72.846731

Ward, J. H. (1963). Hierarchical grouping to optimize an objective function. *Journal of the American Statistical Association*, *58*(301), 236. https://doi.org/10.2307/2282967

Wehmeyer, M. L., Jaworski, L., Jöst, H., Șuleșco, T., Rauhöft, L., Afonso, S. M. M., Neumann, M., Kliemke, K., Lange, U., Kiel, E., Schmidt-Chanasit, J., Sauer, F. G., & Lühken, R. (2024). Host attraction and host feeding patterns indicate generalist feeding of Culex pipiens s.s. and Cx. torrentium. *Parasites & Vectors*, *17*(1), 369. https://doi.org/10.1186/s13071-024-06439-7

Wehrens, R., & Buydens, L. M. C. (2007). Self- and super-organizing maps in R: The kohonen package. *Journal of Statistical Software*, *21*(5). https://doi.org/10.18637/jss.v021.i05

Wehrens, R., & Kruisselbrink, J. (2018). Flexible self-organizing maps in kohonen 3.0. *Journal of Statistical Software*, *87*(7). https://doi.org/10.18637/jss.v087.i07

Weir, B. S., & Cockerham, C. C. (1984). Estimating F-Statistics for the analysis of population structure. *Evolution*, *38*(6), 1358. https://doi.org/10.2307/2408641

Westerband, A. C., Funk, J. L., & Barton, K. E. (2021). Intraspecific trait variation in plants: A renewed focus on its role in ecological processes. *Annals of Botany*, *127*(4), 397–410. https://doi.org/10.1093/aob/mcab011

Wu, X., Kumar, V., Ross Quinlan, J., Ghosh, J., Yang, Q., Motoda, H., McLachlan, G. J., Ng, A., Liu, B., Yu, P. S., Zhou, Z.-H., Steinbach, M., Hand, D. J., & Steinberg, D. (2008). Top 10 algorithms in data mining. *Knowledge and Information Systems*, *14*(1), 1–37. https://doi.org/10.1007/s10115-007-0114-2

Zamani, N., Russell, P., Lantz, H., Hoeppner, M. P., Meadows, J. R. S., Vijay, N., Mauceli, E., di Palma, F., Lindblad-Toh, K., Jern, P., & Grabherr, M. G. (2013). Unsupervised genome-wide recognition of local relationship patterns. *BMC Genomics*, *14*(1), 347. https://doi.org/10.1186/1471-2164-14-347

Zar, J. H. (2010). *Biostatistical analysis*. Pearson.

Zhengtian, Z., Zhiyuan, R., & Xiaoyan, D. (2023). Feature selection for binary classification based on class labeling, SOM, and hierarchical clustering. *Measurement and Control*, *56*(9–10), 1649–1669. https://doi.org/10.1177/00202940231173748
